# A system for phenotype harmonization in the NHLBI Trans-Omics for Precision Medicine (TOPMed) Program

**DOI:** 10.1101/2020.06.18.146423

**Authors:** Adrienne M. Stilp, Leslie S. Emery, Jai G. Broome, Erin J. Buth, Alyna T. Khan, Cecelia A. Laurie, Fei Fei Wang, Quenna Wong, Dongquan Chen, Catherine M. D’Augustine, Nancy L. Heard-Costa, Chancellor R. Hohensee, William Craig Johnson, Lucia D. Juarez, Jingmin Liu, Karen M. Mutalik, Laura M. Raffield, Kerri L. Wiggins, Paul S. de Vries, Tanika N. Kelly, Charles Kooperberg, Pradeep Natarajan, Gina M. Peloso, Patricia A. Peyser, Alex P. Reiner, Donna K. Arnett, Stella Aslibekyan, Kathleen C. Barnes, Lawrence F. Bielak, Joshua C. Bis, Brian E. Cade, Ming-Huei Chen, Adolfo Correa, L. Adrienne Cupples, Mariza de Andrade, Patrick T. Ellinor, Myriam Fornage, Nora Franceschini, Weiniu Gan, Santhi K. Ganesh, Jan Graffelman, Megan L. Grove, Xiuqing Guo, Nicola L. Hawley, Wan-Ling Hsu, Rebecca D. Jackson, Cashell E. Jaquish, Andrew D. Johnson, Sharon LR Kardia, Shannon Kelly, Jiwon Lee, Rasika A. Mathias, Stephen T. McGarvey, Braxton D. Mitchell, May E. Montasser, Alanna C. Morrison, Kari E. North, Seyed Mehdi Nouraie, Elizabeth C. Oelsner, Nathan Pankratz, Stephen S. Rich, Jerome I. Rotter, Jennifer A. Smith, Kent D. Taylor, Ramachandran S. Vasan, Daniel E. Weeks, Scott T. Weiss, Carla G. Wilson, Lisa R. Yanek, Bruce M. Psaty, Susan R. Heckbert, Cathy C. Laurie

## Abstract

Genotype-phenotype association studies often combine phenotype data from multiple studies to increase power. Harmonization of the data usually requires substantial effort due to heterogeneity in phenotype definitions, study design, data collection procedures, and data set organization. Here we describe a centralized system for phenotype harmonization that includes input from phenotype domain and study experts, quality control, documentation, reproducible results, and data sharing mechanisms. This system was developed for the National Heart, Lung and Blood Institute’s Trans-Omics for Precision Medicine (TOPMed) program, which is generating genomic and other omics data for >80 studies with extensive phenotype data. To date, 63 phenotypes have been harmonized across thousands of participants from up to 17 TOPMed studies per phenotype. We discuss the challenges faced in this undertaking and how they were addressed. The harmonized phenotype data and associated documentation have been submitted to National Institutes of Health data repositories for controlled-access by the scientific community. We also provide materials to facilitate future harmonization efforts by the community, which include (1) the code used to generate the 63 harmonized phenotypes, enabling others to reproduce, modify or extend these harmonizations to additional studies; and (2) results of labeling thousands of phenotype variables with controlled vocabulary terms.

---

To increase power in epidemiological analyses, multiple studies are often combined for pooled- or meta-analysis. In both cases, heterogeneities among studies are generally addressed by careful selection and harmonization of study data to include in analyses. In this report, we describe a system for phenotype harmonization, which was developed for the National Heart, Lung and Blood Institute’s (NHLBI) Trans-Omics for Precision Medicine (TOPMed) program (https://www.nhlbiwgs.org/). We define phenotype harmonization as the process by which data variables, each representing a specified phenotype concept, are selected from multiple studies and transformed as needed so that they can be combined and analyzed together. In principle, phenotype harmonization can be achieved prospectively when all contributing studies use the same protocols, an endeavor facilitated by methodological standards such as those developed by the PhenX consortium (1). However, retrospective harmonization is often needed in order to use valuable data that have previously been collected by multiple studies using different phenotype definitions, study designs, data collection procedures, and data structures.

A key goal of the TOPMed program is to identify genetic risk factors for heart, lung, blood, and sleep disorders. To date, the program has generated whole genome sequence data for over 140,000 participants from over 80 different studies, as well as other omics data from many of the same participants. The participating studies have previously gathered extensive phenotype data, including physical measurements, clinical chemistry, questionnaires, clinical registries, and medical imaging. Many of the same phenotypes have been collected in multiple studies, which provides the potential for combined analyses to increase power for detecting the effects of low frequency and rare sequence variants. However, due to substantial heterogeneity in phenotype data among studies and within studies over time, harmonization is required for combined analyses.

Our system for retrospective harmonization of phenotype data in TOPMed includes a collaborative framework, domain expertise, high-quality data inputs, validation of data outputs, rigorous documentation, and respect for stakeholders (i.e., features of the Maelstrom Research guidelines (2)), as well as reproducibility, updating, and sharing of harmonized results derived from controlled-access human data. We describe these features in detail, along with examples of applications to TOPMed study data. We also describe a system for tagging study variables with phenotype concepts for use in future harmonization efforts.

## METHODS

### Overview

The TOPMed Data Coordinating Center (DCC) developed a centralized system for phenotype harmonization, which is outlined in Figure 1. Although we describe the harmonization process as a linear sequence of steps, it was frequently iterative such that the results of later steps required going back and modifying earlier steps. Table 1 provides definitions of terms used here and in the Web Appendix.

**Table 1.**
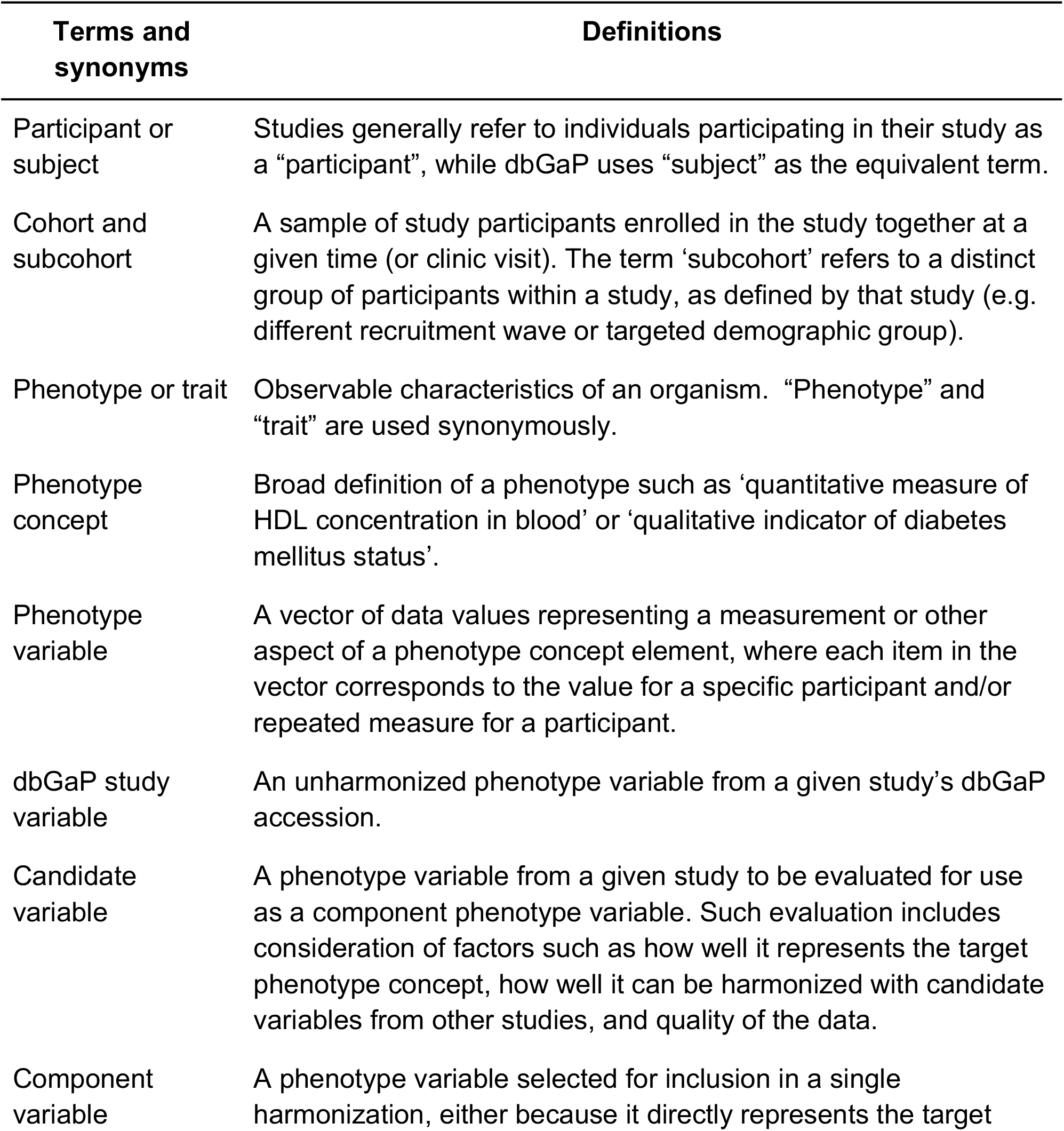
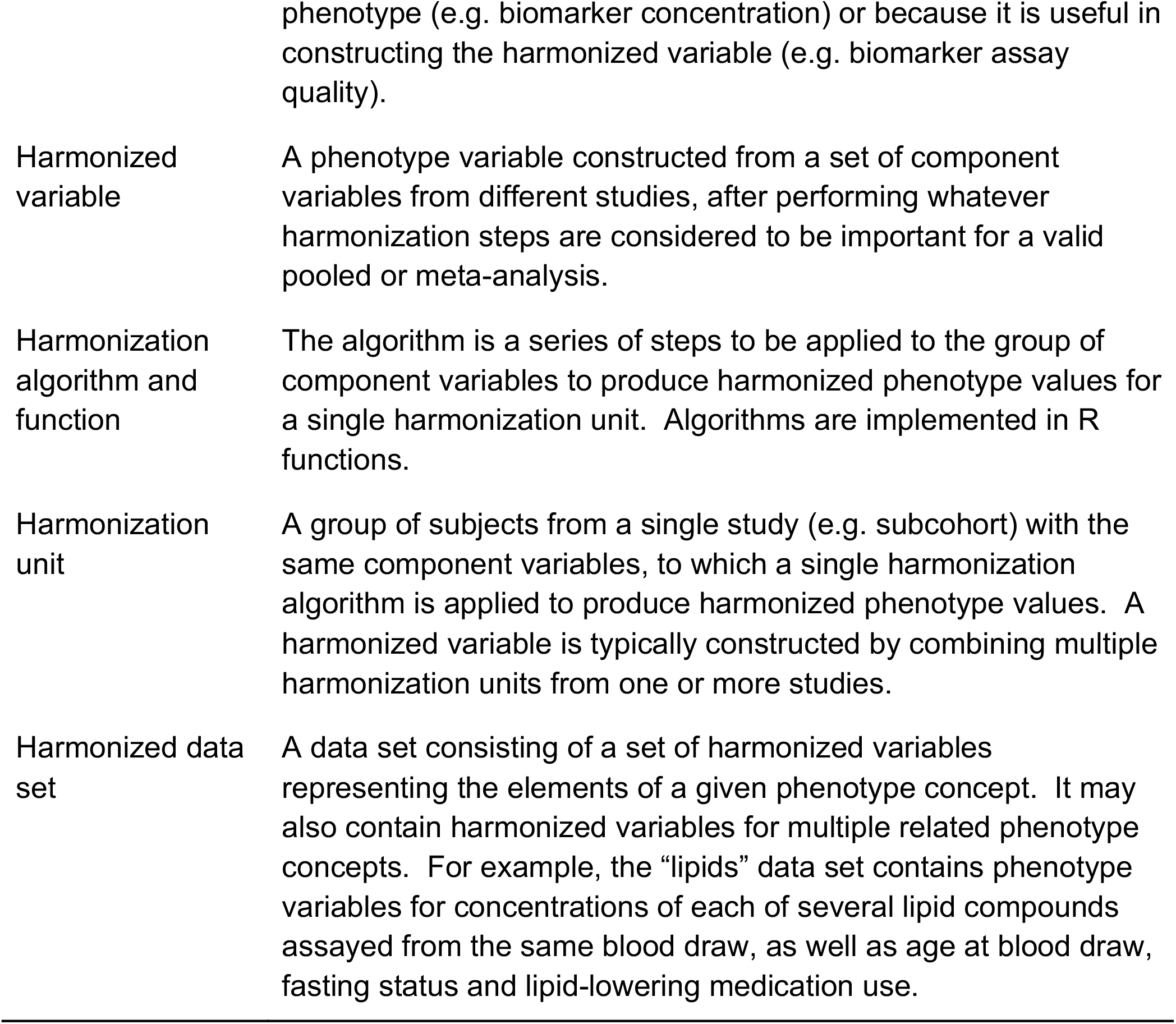
Terminology. Specific terminology used in this report, in the Supplement and in documentation distributed with harmonized phenotype data.

**Figure 1.**
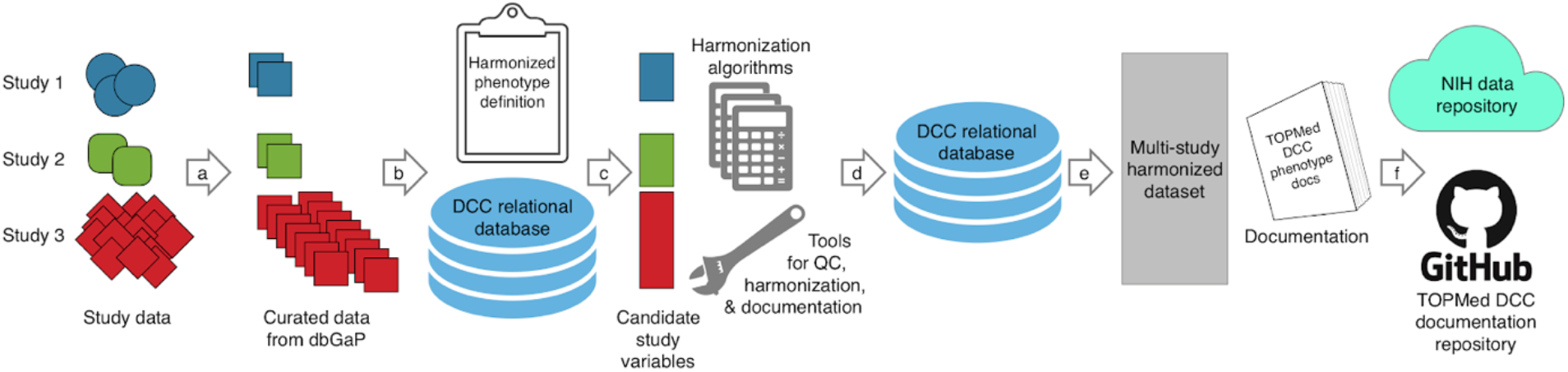
Harmonization system overview. (a) Existing study data in diverse formats are curated by dbGaP, including accessioning and conversion to a consistent file format. (b) Formatted data and associated metadata (e.g. variable descriptions) are stored in a TOPMed DCC relational database. (c) The harmonized phenotype variable is defined, and metadata for multiple studies are searched to identify candidate phenotypic variables that potentially can be harmonized together to produce the desired harmonized variable (harmonization steps 1 and 2). (d) Analytical tools that interact with the DCC database are used for quality control of study variables, implementation of harmonization algorithms, and documentation; harmonized results are added to the same DCC database shown in step b (harmonization steps 3-5). (e) Files containing a multi-study, harmonized data set and associated documentation are produced. (f) Data, metadata and documentation are submitted to an NIH repository for controlled access by the scientific community, while JSON-formatted documentation files containing code and provenance tracking are submitted to a publicly available GitHub repository.

The system tracked the harmonization of each phenotype separately, along with the age at measurement or biosample collection. Each harmonized phenotype variable is assigned a controlled-vocabulary term from the Unified Medical Language System (UMLS) (3). Analysts worked on a group of related phenotype variables at the same time (e.g., high-density lipoprotein cholesterol, low-density lipoprotein cholesterol (LDL-C), total cholesterol, triglycerides, fasting status, and lipid-lowering medication use) which were generally released together in a single dataset (e.g., “Lipids”) -see Table 2. When harmonizing a group of related phenotypes, it is important to use phenotype variables that were measured or collected from a participant at the same time point.

**Table 2.** Harmonized Variables Produced by the DCC. Harmonized variables produced by the DCC. Additional documentation about each variable can be found in the GitHub repository (https://github.com/UW-GAC/topmed-dcc-harmonized-phenotypes). The ‘concept variant number’ at the end of each harmonized variable name differentiates among different implementations of harmonization for the same basic phenotype concept (e.g. cimt_1 and cimt_2 are names for cIMT variables calculated with slightly different harmonization algorithms).

| **Dataset** | **Phenotype concept** | **Harmonized variable name** | **Number of participants** | **Number of studies** |
| --- | --- | --- | --- | --- |
| Atherosclerosis | Coronary artery calcium volume | cac_volume_1 | 11,098 | 2 |
| Atherosclerosis | Coronary artery calcium score | cac_score_1 | 15,042 | 6 |
| Atherosclerosis | Common carotid intima-media thickness | cimt_1 | 35,420 | 6 |
| Atherosclerosis | Common carotid intima-media thickness | cimt_2 | 30,473 | 5 |
| Atherosclerosis | Carotid stenosis | carotid_stenosis_1 | 15,098 | 3 |
| Atherosclerosis | Presence of carotid plaque | carotid_plaque_1 | 27,344 | 5 |
| Baseline Common Covariates | Standing body height | height_baseline_1 | 230,287 | 16 |
| Baseline Common Covariates | Body weight | weight_baseline_1 | 230,657 | 16 |
| Baseline Common Covariates | Ever smoker status | ever_smoker_baseline_1 | 225,271 | 14 |
| Baseline Common Covariates | Current smoker status | current_smoker_baseline_1 | 228,688 | 16 |
| Baseline<br>Common<br>Covariates | Body mass index | bmi_baseline_1 | 230,918 | 17 |
| Blood Cell<br>Count | Basophil<br>concentration in<br>blood | basophil_ncnc_bld_1 | 36,586 | 7 |
| Blood Cell<br>Count | Eosinophil<br>concentration in<br>blood | eosinophil_ncnc_bld_1 | 37,426 | 7 |
| Blood Cell<br>Count | Lymphocyte<br>concentration in<br>blood | lymphocyte_ncnc_bld_1 | 39,702 | 7 |
| Blood Cell<br>Count | Hematocrit level<br>in blood | hematocrit_vfr_bld_1 | 193,469 | 9 |
| Blood Cell<br>Count | Hemoglobin<br>concentration in<br>blood | hemoglobin_mcnc_bld_1 | 193,367 | 9 |
| Blood Cell<br>Count | Monocyte<br>concentration in<br>blood | monocyte_ncnc_bld_1 | 39,647 | 7 |
| Blood Cell<br>Count | Neutrophil<br>concentration in<br>blood | neutrophil_ncnc_bld_1 | 38,285 | 7 |
| Blood Cell<br>Count | Mean corpuscular<br>volume in blood | mcv_entvol_rbc_1 | 44,593 | 7 |
| Blood Cell<br>Count | Mean corpuscular<br>hemoglobin<br>concentration in<br>blood | mchc_mcnc_rbc_1 | 51,293 | 8 |
| Blood Cell<br>Count | Mean corpuscular<br>hemoglobin in<br>blood | mch_entmass_rbc_1 | 39,649 | 7 |
| Blood Cell<br>Count | Platelet<br>concentration in<br>blood | platelet_ncnc_bld_1 | 190,177 | 9 |
| Blood Cell<br>Count | Mean platelet<br>volume in blood | pmv_entvol_bld_1 | 13,816 | 3 |
| Blood Cell Count | Red blood cell concentration in blood | rbc_ncnc_bld_1 | 39,710 | 7 |
| Blood Cell Count | Red cell distribution width | rdw_ratio_rbc_1 | 28,034 | 4 |
| Blood Cell Count | White blood cell concentration in blood | wbc_ncnc_bld_1 | 192,346 | 9 |
| Blood Pressure | Systolic blood pressure | bp_systolic_1 | 225,934 | 14 |
| Blood Pressure | Diastolic blood pressure | bp_diastolic_1 | 225,934 | 14 |
| Blood Pressure | Antihypertensive medication use | antihypertensive_meds_1 | 207,130 | 12 |
| Demographic | Hispanic subgroup | hispanic_subgroup_1 | 18,612 | 4 |
| Demographic | Subcohort identifier | subcohort_1 | 218,747 | 15 |
| Demographic | Reported race | race_1 | 230,994 | 17 |
| Demographic | Reported sex | annotated_sex_1 | 233,030 | 17 |
| Demographic | Reported Hispanic/Latino indicator | ethnicity_1 | 188,905 | 11 |
| Demographic | Geographic recruitment site | geographic_site_1 | 212,529 | 12 |
| Inflammation | CD40 protein concentration in blood | cd40_1 | 4,238 | 2 |
| Inflammation | CRP concentration in blood | crp_1 | 49,536 | 10 |
| Inflammation | E-selectin concentration in blood | eselectin_1 | 1,215 | 1 |
| Inflammation | ICAM1<br>concentration in<br>blood | icam1_1 | 15,876 | 5 |
| Inflammation | IL1β<br>concentration in<br>blood | il1_beta_1 | 708 | 1 |
| Inflammation | IL6 concentration<br>in blood | il6_1 | 20,390 | 5 |
| Inflammation | IL10<br>concentration in<br>blood | il10_1 | 3,455 | 2 |
| Inflammation | IL18<br>concentration in<br>blood | il18_1 | 3,159 | 1 |
| Inflammation | Isoprostane 8-epi-PGF2α<br>concentration in<br>urine | isoprostane_8_epi_pgf2a_1 | 7,523 | 1 |
| Inflammation | Activity of LP-PLA2 in blood | lppla2_act_1 | 18,117 | 3 |
| Inflammation | Mass of LP-PLA2<br>in blood | lppla2_mass_1 | 18,049 | 3 |
| Inflammation | MCP1<br>concentration in<br>blood | mcp1_1 | 7,557 | 1 |
| Inflammation | MMP9<br>concentration in<br>blood | mmp9_1 | 964 | 1 |
| Inflammation | MPO<br>concentration in<br>blood | mpo_1 | 3,162 | 1 |
| Inflammation | OPG<br>concentration in<br>blood | opg_1 | 7,648 | 1 |
| Inflammation | P-selectin<br>concentration in<br>blood | pselectin_1 | 8,037 | 1 |
| Inflammation | TNFα<br>concentration in<br>blood | tnfa_1 | 5,075 | 3 |
| Inflammation | TNFα receptor 1<br>concentration in<br>blood | tnfa_r1_1 | 2,802 | 1 |
| Inflammation | TNF receptor 2<br>concentration in<br>blood | tnfr2_1 | 7,962 | 1 |
| Lipids | Fasting status | fasting_lipids_1 | 64,895 | 11 |
| Lipids | High density<br>lipoprotein<br>concentration in<br>blood | hdl_1 | 65,676 | 11 |
| Lipids | Total cholesterol<br>concentration in<br>blood | total_cholesterol_1 | 65,707 | 11 |
| Lipids | Triglyceride<br>concentration in<br>blood | triglycerides_1 | 65,706 | 11 |
| Lipids | Low density<br>lipoprotein<br>concentration in<br>blood | ldl_1 | 64,715 | 11 |
| Lipids | Lipid-lowering<br>medication use | lipid_lowering_medication_1 | 58,962 | 9 |
| VTE | Age at beginning<br>of followup | vte_followup_start_age_1 | 61,692 | 4 |
| VTE | Prior history of<br>venous<br>thromboembolism | vte_prior_history_1 | 62,445 | 5 |
| VTE | Venous<br>thromboembolism<br>case status | vte_case_status_1 | 63,092 | 6 |

Cluster of differentiation 40 (CD40); C-reactive protein (CRP); Intercellular adhesion molecule 1 (ICAM1); Interleukin 1 beta (IL1 $\beta$ ); Interleukin 6 (IL6); Interleukin 10 (IL10);
Interleukin 18 (IL18); Lipoprotein-associated phospholipase A2 (LP-PLA2); Monocyte chemoattractant protein-1 (MCP1); Matrix metalloproteinase 9 (MMP9); Myeloperoxidase (MPO); Osteoprotegerin (OPG); Tumor necrosis factor (TNF); 8-epi-prostaglandin F2 $\alpha$ (8-epi-PGF2 $\alpha$ )

Phenotype concepts and harmonization algorithms were developed in collaboration with phenotype experts in TOPMed Working Groups (WGs). Study-specific issues were resolved in collaboration with investigators and/or data managers from the relevant study. The harmonization system described here was implemented primarily by DCC scientists, with advice from WGs and study experts. This report does not document other harmonization efforts involving TOPMed studies that were performed independently of the DCC (e.g., Oelsner et al. (4) or the independent efforts of various TOPMed WGs).

The information technology supporting phenotype harmonization consisted of a locally-hosted relational database and associated applications. A custom R package was used to interact with the database and a series of Python and R scripts were run by analysts to perform harmonization. The codebase also allowed addition of new study and harmonized data to the database; retrieval of existing study data in their original structure; and production of harmonized datasets and documentation for distribution to investigators. A custom web application was used to search the publicly-available metadata for relevant study variables.

### Obtaining and processing study data

All study phenotype data and associated metadata were obtained from the National Institutes of Health (NIH) database of Genotypes and Phenotypes (dbGaP; https://www.ncbi.nlm.nih.gov/gap/) (5), which provides controlled access for the scientific community. Use of dbGaP data provides a mechanism to track the provenance of a harmonized phenotype variable using dbGaP accession numbers assigned to multiple data entities, including studies, datasets, and individual variables within datasets. The harmonization system leverages work already done by dbGaP to curate data into a consistent file format, include metadata (e.g., variable descriptions and types), and perform some value-checking based on the data type. Use of dbGaP data enables reproducibility of harmonized phenotypes, as scientific investigators can obtain the same datasets (via controlled access). For each study, the harmonization process included all participants with data available on dbGaP, rather than only those being sequenced in TOPMed.

After obtaining approval for access to a study’s dbGaP accession, all available phenotype data and associated metadata were imported into a relational database. An overview of the database is provided in the Web Appendix (section S2).

Studies participating in TOPMed were approved by institutional review boards and participants provided informed consent, including information regarding data use limitations and guidelines for sharing data on dbGaP. Even though the DCC harmonized data for all participants available on dbGaP, the resulting harmonized phenotypes may only be analyzed for participants whose dbGaP consent group allows research in that area.

### Harmonization steps

The following harmonization steps are focused on producing each individual harmonized phenotype variable (although, as noted above, several related phenotypes may be harmonized in parallel and provided to users within a single dataset). The Web Appendix (section S3) provides detailed examples of these steps.

#### Step 1: Define the harmonized phenotype variable

The first step was to develop a precise definition of the target harmonized phenotype variable, often including references to specific assay or measurement methods, time points in longitudinal studies, and other relevant factors. For example, for LDL-C concentration in blood, the definition might specify calculation according to the Friedewald equation (6) using high-density lipoprotein cholesterol, total cholesterol, and triglycerides measurements, all from the same blood sample drawn at the baseline clinic visit. The initial definition was sometimes modified to accommodate heterogeneities in the data available in different studies as they were discovered in subsequent steps.

#### Step 2: Identify ‘candidate’ phenotype variables across contributing studies

The next step was to identify candidate dbGaP study variables that could potentially be used for calculating the target harmonized phenotype variable, as well as corresponding variables containing age at measurement or biosample collection. Because controlled vocabulary usage is limited in dbGaP datasets, this process consisted of searching variable names, descriptions, and encoded values. The tagging project described below was implemented to facilitate this process for both DCC harmonization and other harmonization efforts by the scientific community.

Once an initial set of candidate variables was identified, the selection was refined by assessing compatibility with the definition of the target harmonized phenotype and for methodological equivalence across studies. This process often involved selecting among different methods of measuring the phenotype and/or choosing the most appropriate variable from a set of repeated measurements. Analysts generally consulted publicly-available study protocols, phenotype domain experts in the relevant WG, and study liaisons, who know the intricacies of their study’s data.

In some cases, a new harmonized variable was constructed from previously harmonized component variables (e.g., a harmonized body mass index variable from previously harmonized height and weight variables).

#### Step 3: Perform quality control (QC) on candidate variables

QC on selected candidate variables was performed to check whether the observed values were consistent with expected ranges, investigate any unexpected distributions, and check that the data were internally consistent with other related study variables.

Batch effects were also evaluated when relevant batching information was available. If QC issues were identified for a candidate variable, analysts decided, in consultation with the WG and study liaisons, whether an alternative variable from the same study could be used, or whether the study should be excluded from the harmonization for this phenotype. Individual data points with impossible values (such as a negative analyte concentration) were excluded from the harmonized phenotype variable. Extreme but theoretically possible values were noted in the documentation, but were not excluded because (1) the definition of extremity is often difficult and subjective; (2) TOPMed whole genome sequencing has discovered millions of rare variants, some of which may be causing extreme phenotypic values; and (3) users may prefer to handle extreme values differently (e.g., by excluding or winsorizing at different values). Therefore, the decision about how to handle extreme values in analyses was left to downstream users of the data.

QC results for candidate study variables were used to determine which ones would be used as ‘component’ variables in subsequent harmonization steps. The final set of component variables was chosen only after QC of the multi-study harmonized variable (see Step 5).

#### Step 4: Construct harmonization algorithms

The next step was to specify the algorithms to be used in transforming component variables into the harmonized variable. An algorithm was developed for each ‘harmonization unit’, which consists of a group of participants from a single study with component variables that can be harmonized in the same way. Each algorithm was implemented as an R function that accepts the component variables as input and returns the harmonized values and age at measurement. The algorithm might be as simple as giving each component variable a consistent name across studies or converting to a common unit of measurement, but often included more complicated steps, such as averaging repeated measurements or creating a smoking status variable from multiple questionnaire responses.

#### Step 5: Produce and QC multi-study harmonized phenotype

After harmonization algorithms were implemented for each contributing study, the harmonized values were calculated and combined across harmonization units and studies using in-house R scripts.

This draft of the multi-study harmonized variable was then assessed for homogeneity of values among studies and harmonization units within studies. This process included a comparison of means and standard deviations of continuous variables or frequencies of categorical variables, by study, subcohort, and other relevant subgroups within each study. For continuous variables, we also inspected the distributions of residuals after fitting a linear model with age, sex, and harmonized race. If any issues arose in this process, analysts evaluated whether the harmonization unit in question should be excluded or whether different component variables should be used for harmonization.

When QC checks were complete and the set of contributing studies was finalized, analysts summarized the results and any additional information relevant for analysis in a free text document referred to as ‘harmonization comments’. This document may include notes about the presence of a notable cluster of outliers; differences among studies that were not considered important enough for removal of a study from harmonization; or variation among studies or subcohorts in assay methodology. These notes allow users to flexibly choose how to account for potential effects in analysis. See Boxes in the Web Appendix section S3.5 for examples of these comments.

The final multi-study harmonized variable was then added to the DCC’s phenotype database. The information added included metadata and data values for the harmonized variable, as well as the set of component variables and harmonization functions used to generate the harmonized data values.

### Distributing harmonization results to the scientific community

The DCC provides a package of datasets and documentation using information stored in the database. Each dataset generally includes a group of related harmonized variables plus age at measurement for each variable. The documentation includes files in JavaScript Object Notation (JSON) format containing code that allows a user to reproduce or modify harmonized variables once they obtain access to the specified study data from dbGaP (see Web Appendix section S6 for details). In addition, the harmonized variables described in Table 2 have been submitted to dbGaP and to the NHLBI BioData Catalyst repository for distribution to the scientific community via application to dbGaP.

### Updating harmonized variables

A harmonized phenotype variable often needed to be updated to include additional studies and/or to incorporate dbGaP updates to the component study variables from previously included studies. These updates were semi-automated because the relational database contained all of the information necessary to recreate the harmonized phenotype. Updates of all variables in a dataset were typically made at the same time.

### Tagging phenotype variables to facilitate future harmonization

While the detailed harmonization process described above produces well-documented, reproducible, and updateable harmonized phenotype variables, other investigators may want to carry out harmonization differently (e.g., using different component variables; a different harmonization algorithm; a different harmonized phenotype definition; or additional or different timepoints). They may also need to harmonize a phenotype the DCC has not worked on yet. To facilitate identification of candidate variables for harmonization, we worked with study and domain experts to label TOPMed dbGaP study variables with controlled vocabulary terms to indicate the phenotype concept they represent (i.e., ‘variable tagging’). Study variables were tagged with 65 important phenotype concepts from heart, lung, blood, sleep and demographic domains (Web Table S7). Harmonized phenotype variables for 27 of the 65 concepts have been constructed already, but many more are possible, even for the same concept. The remaining harmonized variables represent phenotype concepts that were not directly included among the 65 originally identified concepts.

Study variable tagging was done via a database-backed web application with built-in data validation. The DCC worked with representatives from seven large cohort studies to identify all of their studies’ dbGaP study variables that fit one or more of the 65 phenotype concepts, and to label them with the appropriate phenotype concept tag(s) and corresponding UMLS term(s). DCC phenotype team members also tagged variables for the remaining studies available at the time. We performed careful quality review of all tagged variables to ensure consistency and accuracy of the tagging across studies. Details of this process are described in the Web Appendix (section S7).

## RESULTS

### Phenotypes harmonized

A total of 63 harmonized variables were constructed across multiple TOPMed studies (up to 17 for some variables) belonging to 8 phenotype datasets (Table 2). Within each dataset, the variables generally represent related phenotypes that are analyzed together (except for common covariates and demographic variables). Web Figure S4 shows histograms of the harmonized variables.

### QC issues in harmonization

Four types of issues arose frequently during QC of study and harmonized phenotype variables: (1) notable differences among studies/subcohorts in the distributions of quantitative measures or frequencies of categorical phenotypes; (2) variation among studies/subcohorts in methods for how the same phenotype was assessed or measured; (3) extreme (sometimes impossible) values of quantitative measures; and (4) inconsistencies in the values of related phenotypes. In general, the resolution of these issues was highly phenotype-dependent and relied on expertise from the WG members and study liaisons. Here we give some examples of how these issues were detected and resolved, with more detail and examples in the Web Appendix (sections S3.3 and S3.5).

When producing a harmonized variable, we compared distributions across studies and subcohorts within studies to identify differences that might be due to errors or unusual features of a given study. We show an example of this type of comparison in Figure 2 for the ever smoker harmonized variable. It is clear that two study/subcohorts, F and G1, had a much higher proportion of smokers, while a third study/subcohort, E, had a much lower proportion of smokers than the average. In two cases, the proportions can be explained by the studies’ recruitment strategies; study/subcohort F targeted smokers for enrollment into the study (7), while study/subcohort E included children (8). Because these differences can be explained by recruitment strategy, no modification to the harmonization was needed. Further exploration of subcohort G1 showed that this high proportion was due to an unlabeled missing value code in one of the component variables. We corrected the harmonization algorithm to account for this missing code, and the differences in the proportions of smokers by subcohort after this correction were much smaller.

**Figure 2.**
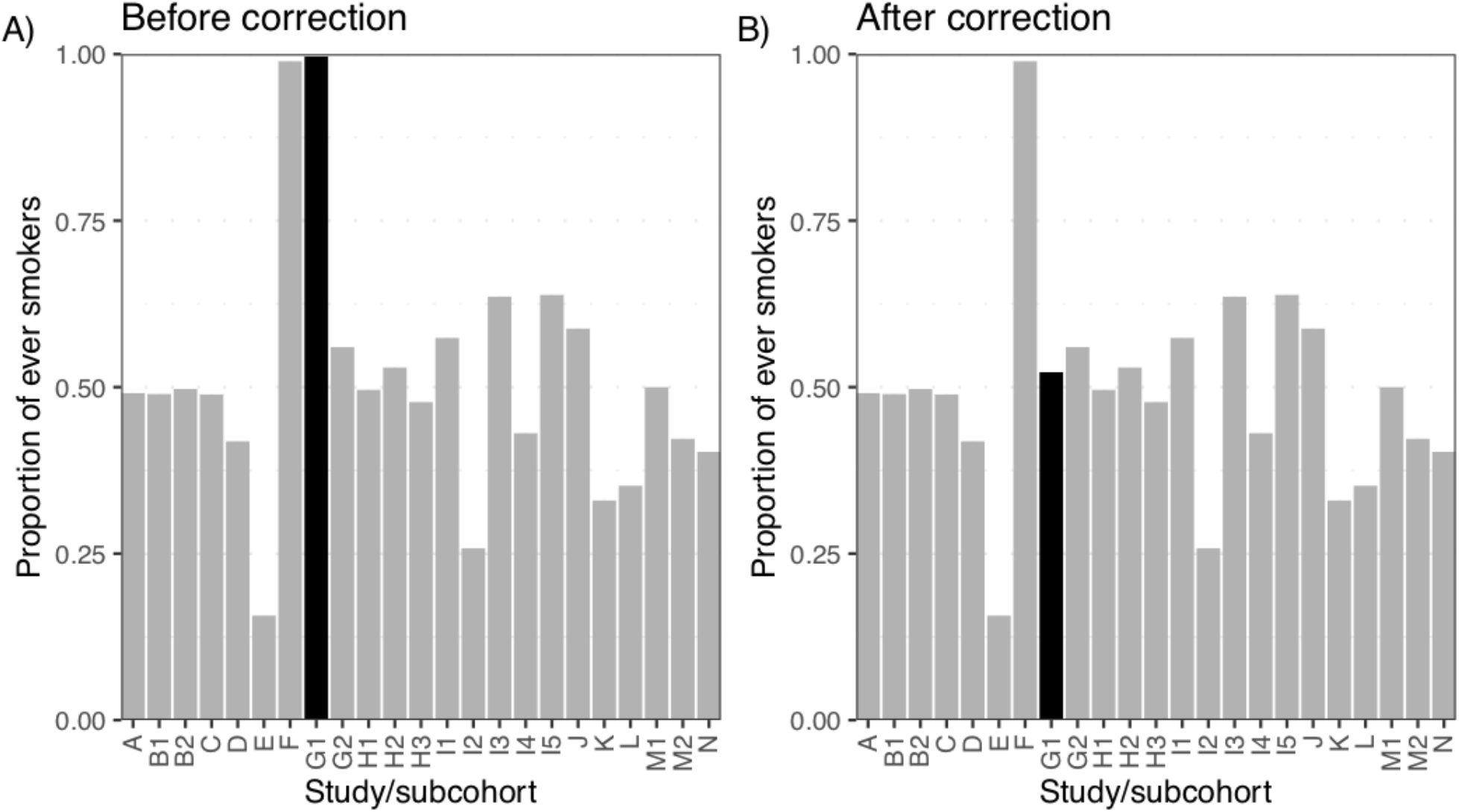
Proportion of ever smokers from the harmonized ever_smoker_baseline_1 variable by (anonymized) study subcohort. In both plots, different studies are labeled by a letter (e.g., B), and different subcohorts within each study (if applicable) are labeled by appending a number to the study letter (e.g., B1 and B2). A) Proportion of smokers by study/subcohort after initial harmonization. Three study/subcohorts (E, F, and G1) have much smaller or larger proportions compared to the majority of other studies. B) Proportion of smokers by study/subcohort after correcting study/subcohort G1 (shown in black) for an unlabeled missing value code.

A second example of harmonized phenotype QC is shown in Figure 3. The final QC for interleukin-6 concentration included inspection of the distribution of values by study and subcohort as well as the residuals after adjusting for age, sex, and race. The distribution for one study was much wider in range and generally had higher values compared with the other study/subcohorts (Study E in Figure 3). These differences remained even after adjusting for age, sex, and race. The DCC consulted with study liaisons, and decided to remove that study from harmonization because the reason for the unusual distribution could not be determined.

**Figure 3.**
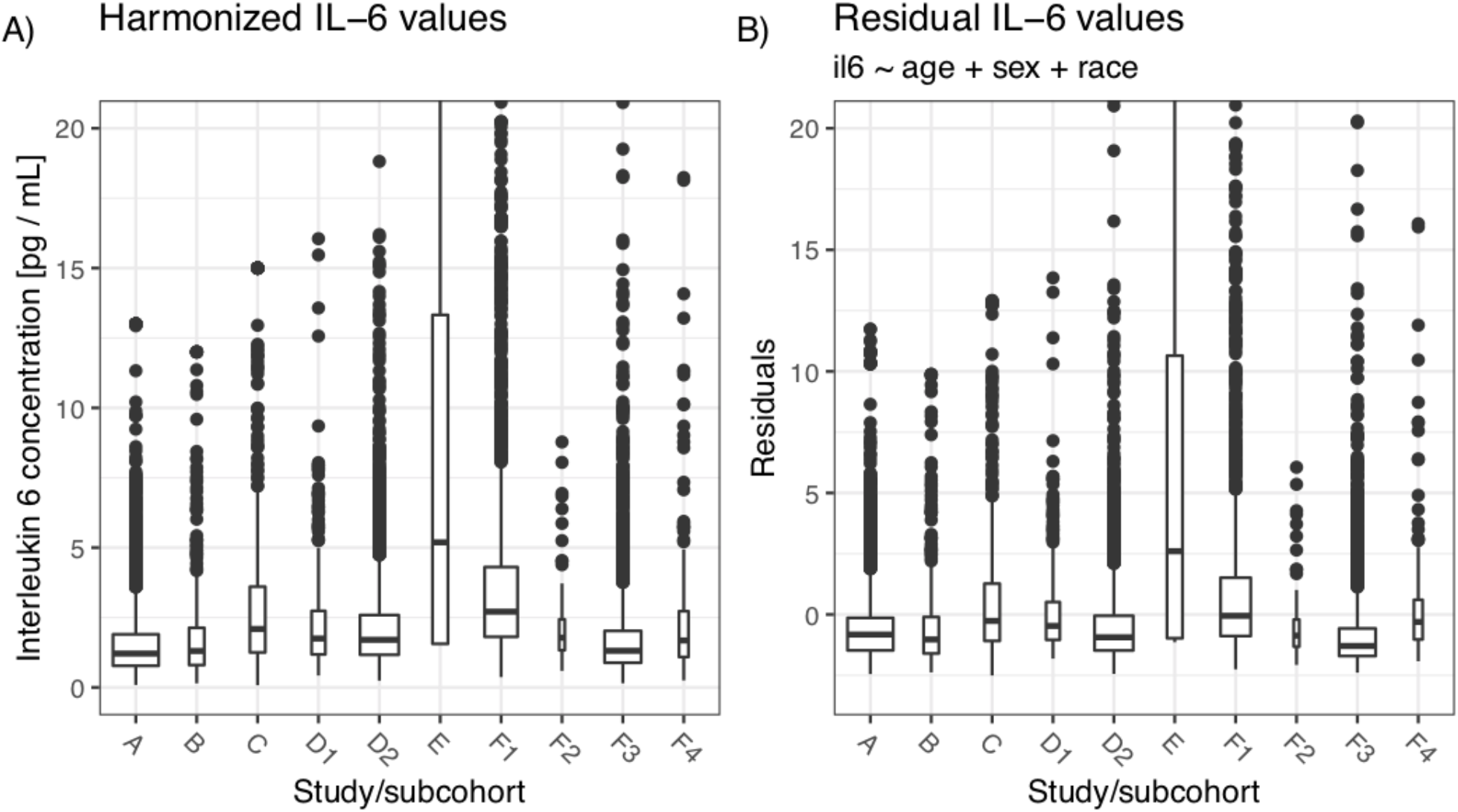
Distribution of harmonized IL-6 values by study/subcohort. In both plots, different studies are labeled by a single letter (e.g., D), and different subcohorts within each study (if applicable) are labeled by appending a number to the study letter (e.g., D1 and D2). A) Harmonized IL-6 values. The interquartile range for Study E is much larger than the other study/subcohorts. B) Residuals of a linear model (il6 ∼ age + sex + race) by study/subcohort. The large differences between Study E and the other study/subcohorts remain after adjusting the values for age, sex, and race.

There is often a trade-off between the homogeneity of a harmonized variable and achieving a large sample size by including many studies (9–11). This issue generally arose when studies measured different aspects of a harmonized variable (e.g., measurements of the thickness of different carotid artery walls for calculating common carotid intima-media thickness) or used different methods to collect a similar measurement (e.g., different assay methods for inflammation phenotypes). In these cases, WG and study members were involved in decisions about whether to exclude studies or modify the definition of the harmonized phenotype.

We sometimes found biologically invalid data points, such as diastolic greater than systolic blood pressure for some participants, or unexpected relationships between variable values, such as white blood cell subtype counts not adding up to the total count. Other inconsistencies were found in participant responses to questionnaires (e.g., participants who report that they have never smoked but also report smoking a non-zero number of cigarettes per day). As noted in the Methods section, impossible data values were typically not included in the harmonized variable, while the potentially valid but extreme values are retained but noted in the harmonization comments.

### Reproducibility of harmonized phenotype variables

We have successfully reproduced several of our harmonized variables exactly using only the JSON documentation provided in our public GitHub repository (https://github.com/UW-GAC/topmed-dcc-harmonized-phenotypes), along with the specified study data files from dbGaP (via controlled access). The repository also includes a fully reproducible example using simulated dbGaP data that instructs users about how to reproduce the harmonized variables using the documentation.

### DCC phenotype tagging results

We tagged dbGaP study variables with UMLS terms representing 65 phenotype concepts in 16 domains. Web Table S7 provides descriptions, detailed tagging instructions, and UMLS terms for each phenotype concept. A total of 16,671 dbGaP study variables from 17 studies were tagged with relevant UMLS phenotype terms.

Because some variables were tagged with multiple phenotype terms, there are 17,063 unique pairings of dbGaP variable and UMLS phenotype term. Table 3 shows the number of variables available in each study, the number tagged, and the proportion tagged. The latter varies according to variation among studies in the breadth and depth of phenotype domains for which they have collected data. For example, the Framingham Heart Study has many variables in domains that are not part of the 65 phenotype concepts chosen for tagging, such as bone mineral density measurements.

**Table 3.**
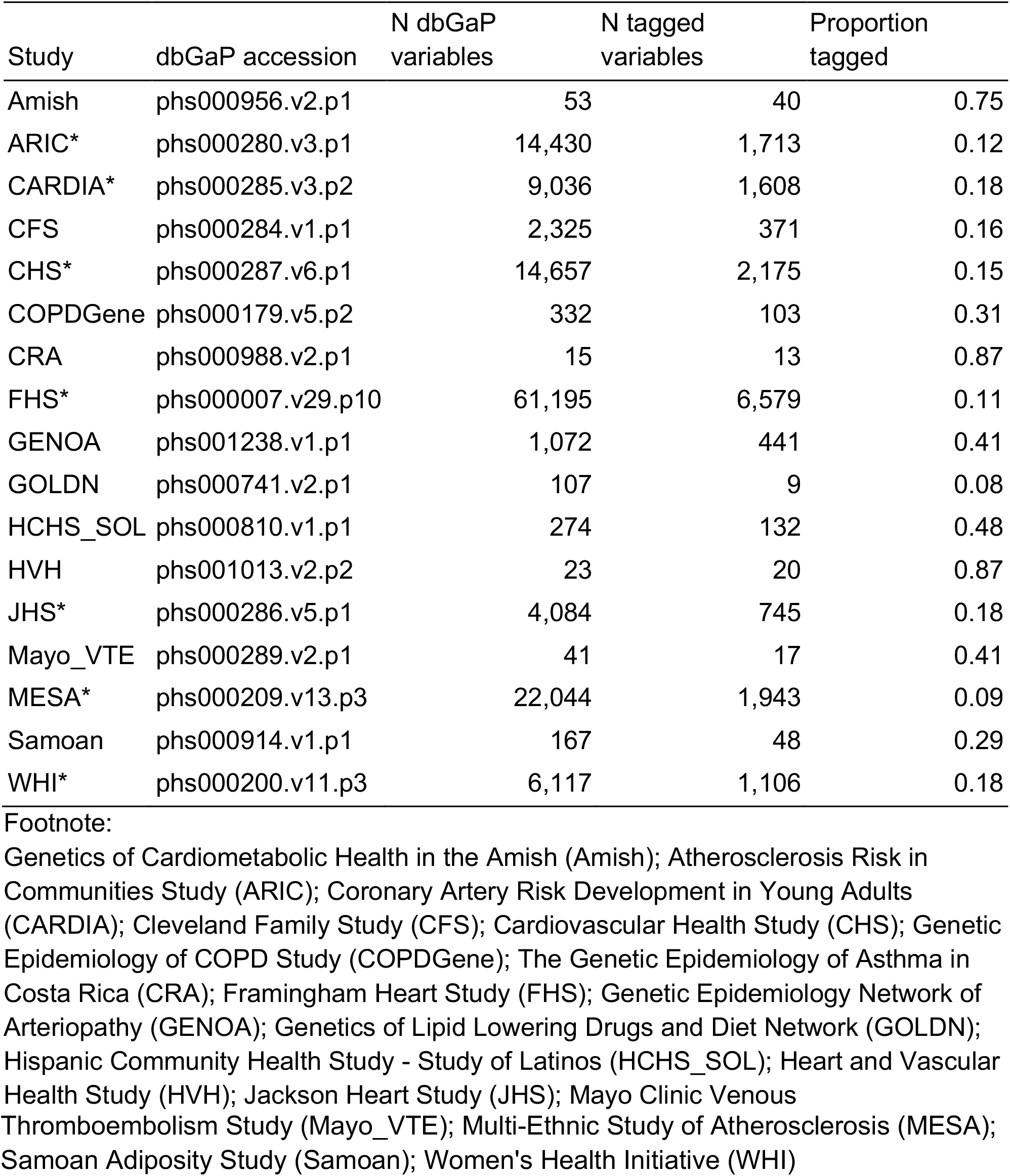
Variables Tagged With Controlled Vocabulary Phenotype Concepts. The number and proportion of variables tagged with controlled vocabulary phenotype concepts for each study. Study names marked with an asterisk indicate that initial tagging was completed by study data experts; those without an asterisk were tagged by DCC analysts.

### Data availability

The study data used as input for harmonization are available to the scientific community from dbGaP via controlled-access. In a single application, a user can apply for access to all dbGaP study accessions provided in the documentation. In addition, the harmonized data in Table 2 have been submitted to dbGaP and to a new NHLBI data repository, BioData Catalyst (https://biodatacatalyst.nhlbi.nih.gov/). In both cases, access will be through application to dbGaP.

We worked with dbGaP scientists to make the tagging results available in dbGaP searches and visible on dbGaP variable pages. Detailed information on how to access and use this information is available on the TOPMed website (https://www.nhlbiwgs.org/dcc-pheno).

## DISCUSSION

The TOPMed program was designed to add cutting-edge genomics and other omics data to over 80 studies with extensive characterization of heart, lung, blood and sleep phenotypes. Because phenotype data in the contributing studies are quite heterogeneous, retrospective harmonization is critical to achieving the program’s goals. The harmonization system described in this report has been used to harmonize 63 phenotypes for several WGs, members of which are using them in many different analyses, primarily genotype-phenotype association studies. Some of these studies have been published (e.g., 12–15) and many others are in preparation.

An important consideration in the design of our harmonization system was the ability to share harmonized phenotypes with the broader scientific community. This goal is challenging because the study data, and any individual-level derivations thereof, require controlled access due to human subject privacy and consent restrictions. We addressed this problem by obtaining the study data for harmonization from dbGaP, which can be accessed by the scientific community; by providing detailed documentation about the component variables and algorithms for each variable; and by returning the harmonized data to NIH-designated repositories. The consent type for a given participant’s harmonized data is inherited from the dbGaP data used as components for the harmonization.

Reproducibility of results is difficult to ensure with confidential data (16). Harmonized data produced by our system are fully reproducible because of the availability of study data, provenance tracking, harmonization code, and other documentation. However, exact reproducibility can only be ensured if a user has access to the same version of the data that was used in harmonization, as studies can update or even remove variables from their dbGaP accessions.

A limitation of our process for phenotype harmonization is that it was very labor-intensive and does not scale readily to the thousands of phenotypes available in TOPMed and other similar programs. Identification and QC of study and harmonized variables are largely manual and would be very difficult to automate, but without performing the steps described here, the utility of results may be compromised. Because of these scalability issues, we provide the following materials to help other investigators perform their own harmonizations:

1. Code and documentation sufficient to allow others to reproduce, modify or expand upon our harmonizations.
2. Detailed examples of the types of QC performed, issues that arose, and how they were resolved (see Web Appendix sections S3.3 and S3.5). We expect this information will prove useful to investigators working on a broad range of phenotypes and may also be helpful in applications to funding agencies regarding the level of resources required for useful harmonization efforts.
3. Thousands of dbGaP variables tagged with 65 phenotype concepts, which can be used directly by other investigators for the largely manual and time-consuming step of identifying the study variables needed for harmonization. The tagging data also provide a gold standard, human-curated data set for developing automated approaches to identifying variables that fit a specific phenotype concept.

We also suggest the following for future phenotype harmonization efforts:

1. Full documentation of harmonization including code, procedures and input data provenance so that others can reproduce and extend their work. Sharing this documentation can benefit the scientific community without sharing the actual harmonized phenotype values (which requires complicated data-sharing arrangements).
2. Studies sharing phenotype data with the community (a) structure data tables so that each phenotype variable (i.e., table column) contains data corresponding to only one phenotype concept and (b) provide controlled vocabulary term(s) from UMLS or its component ontologies for each phenotype variable.
3. Studies currently collecting data consider use of standardized protocols, such as developed by the PhenX consortium (1), to reduce the need for retrospective harmonization in the future.

## Supporting information

Web Appendix

Web Table S7

## ACKNOWLEDGMENTS

Adrienne M. Stilp and Leslie S. Emery contributed equally to the work.

This work was funded by numerous grants and contracts from the National Institutes of Health.

The Trans-Omics in Precision Medicine (TOPMed) program was supported by the National Heart, Lung and Blood Institute (NHLBI), with core services provided by the TOPMed Informatics Research Center (3R01HL-117626-02S1; contract HHSN268201800002I) and the TOPMed Data Coordinating Center (R01HL-120393; U01HL-120393; contract HHSN268201800001I). We gratefully acknowledge the studies and participants who provided biological samples and data for TOPMed. Phenotype harmonization activities were funded in part with Federal funds from the National Heart, Lung, and Blood Institute (NHLBI), National Institutes of Health (NIH), Department of Health and Human Services, under Contract No. HHSN26820180001I. Additional harmonization funding was provided by NHLBI grant 5 U01 HL 120393-04. Phenotype variable tagging was funded by grant supplement 3 U01 HL 120393-04S2 from NHLBI and the NIH Office of the Director, as part of the NIH Data Commons Pilot Phase Consortium.

NHLBI TOPMed: Genetics of Cardiometabolic Health in the Amish (Amish)

We gratefully thank our Amish community and research volunteers for their long-standing partnership in research, and acknowledge the dedication of our Amish liaisons, field workers and the Amish Research Clinic staff, without which these studies would not have been possible. The TOPMed component of the Amish Research Program was supported by NIH grants R01 HL121007, U01 HL072515, and R01 AG18728.

NHLBI TOPMed: Atherosclerosis Risk in Communities Study (ARIC)

The Atherosclerosis Risk in Communities study has been funded in whole or in part with Federal funds from the National Heart, Lung, and Blood Institute, National Institutes of Health, Department of Health and Human Services (contract numbers HHSN268201700001I, HHSN268201700002I, HHSN268201700003I, HHSN268201700004I and HHSN268201700005I). The authors thank the staff and participants of the ARIC study for their important contributions.

NHLBI TOPMed: Coronary Artery Risk Development in Young Adults (CARDIA) The Coronary Artery Risk Development in Young Adults Study (CARDIA) is conducted and supported by the National Heart, Lung, and Blood Institute (NHLBI) in collaboration with the University of Alabama at Birmingham (HHSN268201800005I & HHSN268201800007I), Northwestern University (HHSN268201800003I), University of Minnesota (HHSN268201800006I), and Kaiser Foundation Research Institute (HHSN268201800004I). CARDIA was also partially supported by the Intramural Research Program of the National Institute on Aging (NIA) and an intra-agency agreement between NIA and NHLBI (AG0005).

NHLBI TOPMed: Cleveland Family Study -WGS Collaboration (CFS) The Cleveland Family Study has been supported in part by National Institutes of Health grants (R01-HL046380, KL2-RR024990, R35-HL135818, and R01-HL113338).

NHLBI TOPMed: Cardiovascular Health Study (CHS)

Cardiovascular Health Study: This research was supported by contracts HHSN268201200036C, HHSN268200800007C, HHSN268201800001C, N01HC55222, N01HC85079, N01HC85080, N01HC85081, N01HC85082, N01HC85083, N01HC85086, and grants U01HL080295 and U01HL130114 from the National Heart, Lung, and Blood Institute (NHLBI), with additional contribution from the National Institute of Neurological Disorders and Stroke (NINDS). Additional support was provided by R01AG023629 from the National Institute on Aging (NIA). A full list of principal CHS investigators and institutions can be found at CHS-NHLBI.org. The content is solely the responsibility of the authors and does not necessarily represent the official views of the National Institutes of Health.

NHLBI TOPMed: Genetic Epidemiology of COPD Study (COPDGene)

The COPDGene project described was supported by Award Number U01 HL089897 and Award Number U01 HL089856 from the National Heart, Lung, and Blood Institute. The content is solely the responsibility of the authors and does not necessarily represent the official views of the National Heart, Lung, and Blood Institute or the National Institutes of Health. The COPDGene project is also supported by the COPD Foundation through contributions made to an Industry Advisory Board comprised of AstraZeneca, Boehringer Ingelheim, GlaxoSmithKline, Novartis, Pfizer, Siemens and Sunovion. A full listing of COPDGene investigators can be found at: http://www.copdgene.org/directory

NHLBI TOPMed: The Genetic Epidemiology of Asthma in Costa Rica -Asthma in Costa Rica cohort (CRA)

The Genetic Epidemiology of Asthma in Costa Rica (CRA) was funded by Grant R37 HL066289-14 and Grant P01 HL132825 from the National Heart, Lung and Blood Institute.

NHLBI TOPMed: Framingham Heart Study (FHS)

The Framingham Heart Study (FHS) acknowledges the support of contracts NO1-HC-25195, HHSN268201500001I and 75N92019D00031 from the National Heart, Lung and Blood Institute and grant supplement R01 HL092577-06S1 for this research. We also acknowledge the dedication of the FHS study participants without whom this research would not be possible.

NHLBI TOPMed: Genetic Epidemiology Network of Arteriopathy (GENOA)

Support for GENOA was provided by the National Heart, Lung and Blood Institute (HL054457, HL054464, HL054481, HL119443, HL087660, and HL085571) of the National Institutes of Health.

NHLBI TOPMed: Genetics of Lipid Lowering Drugs and Diet Network (GOLDN) GOLDN biospecimens, baseline phenotype data, and intervention phenotype data were collected with funding from National Heart, Lung and Blood Institute (NHLBI) grant U01 HL072524. Whole-genome sequencing in GOLDN was funded by NHLBI grant R01 HL104135 and supplement R01 HL104135-04S1.

NHLBI TOPMed: Hispanic Community Health Study -Study of Latinos (HCHS_SOL) The Hispanic Community Health Study/Study of Latinos is a collaborative study supported by contracts from the National Heart, Lung, and Blood Institute (NHLBI) to the University of North Carolina (HHSN268201300001I / N01-HC-65233), University of Miami (HHSN268201300004I / N01-HC-65234), Albert Einstein College of Medicine (HHSN268201300002I / N01-HC-65235), University of Illinois at Chicago – HHSN268201300003I / N01-HC-65236 Northwestern Univ), and San Diego State University (HHSN268201300005I / N01-HC-65237). The following Institutes/Centers/Offices have contributed to the HCHS/SOL through a transfer of funds to the NHLBI: National Institute on Minority Health and Health Disparities, National Institute on Deafness and Other Communication Disorders, National Institute of Dental and Craniofacial Research, National Institute of Diabetes and Digestive and Kidney Diseases, National Institute of Neurological Disorders and Stroke, NIH Institution-Office of Dietary Supplements.

NHLBI TOPMed: Heart and Vascular Health Study (HVH)

The Heart and Vascular Health Study was supported by grants HL068986, HL085251, HL095080, and HL073410 from the National Heart, Lung, and Blood Institute.

NHLBI TOPMed: Jackson Heart Study (JHS)

The Jackson Heart Study (JHS) is supported and conducted in collaboration with Jackson State University (HHSN268201800013I), Tougaloo College (HHSN268201800014I), the Mississippi State Department of Health (HHSN268201800015I) and the University of Mississippi Medical Center (HHSN268201800010I, HHSN268201800011I and HHSN268201800012I) contracts from the National Heart, Lung, and Blood Institute (NHLBI) and the National Institute on Minority Health and Health Disparities (NIMHD). The authors also wish to thank the staffs and participants of the JHS.

NHLBI TOPMed: Mayo Clinic Venous Thromboembolism Study (Mayo_VTE) Funded, in part, by grants from the National Institutes of Health, National Heart, Lung and Blood Institute (HL66216 and HL83141). the National Human Genome Research Institute (HG04735, HG06379), and research support provided by Mayo Foundation.

NHLBI TOPMed: Multi-Ethnic Study of Atherosclerosis (MESA)

Whole genome sequencing (WGS) for the Trans-Omics in Precision Medicine (TOPMed) program was supported by the National Heart, Lung and Blood Institute (NHLBI). WGS for “ NHLBI TOPMed: Multi-Ethnic Study of Atherosclerosis (MESA)” (phs001416.v1.p1) was performed at the Broad Institute of MIT and Harvard (3U54HG003067-13S1). Centralized read mapping and genotype calling, along with variant quality metrics and filtering were provided by the TOPMed Informatics Research Center (3R01HL-117626-02S1). Phenotype harmonization, data management, sample-identity QC, and general study coordination, were provided by the TOPMed Data Coordinating Center (3R01HL-120393-02S1).

MESA and the MESA SHARe project are conducted and supported by the National Heart, Lung, and Blood Institute (NHLBI) in collaboration with MESA investigators. Support for MESA is provided by contracts 75N92020D00001, HHSN268201500003I, N01-HC-95159, 75N92020D00005, N01-HC-95160, 75N92020D00002, N01-HC-95161, 75N92020D00003, N01-HC-95162, 75N92020D00006, N01-HC-95163, 75N92020D00004, N01-HC-95164, 75N92020D00007, N01-HC-95165, N01-HC-95166, N01-HC-95167, N01-HC-95168, N01-HC-95169, UL1-TR-000040, UL1-TR-001079, UL1-TR-001420, UL1-TR-001881, and DK063491.

NHLBI TOPMed: Samoan Adiposity Study (Samoan)

Data collection was funded by NIH grant R01-HL093093. We thank the Samoan participants of the study and local village authorities. We acknowledge the Samoan Ministry of Health and the Samoa Bureau of Statistics for their support of this research.

NHLBI TOPMed: Women’s Health Initiative (WHI)

The WHI program is funded by the National Heart, Lung, and Blood Institute, National Institutes of Health, U.S. Department of Health and Human Services through contracts HHSN268201600018C, HHSN268201600001C, HHSN268201600002C, HHSN268201600003C, and HHSN268201600004C.

Additional support was provided to some authors:

Nora Franceschini (NIH grants R01-MD012765, R01-DK117445, R21-HL140385); Patrick Ellinor (NIH grants R01HL092577, R01HL128914, K24HL105780);

Alex Reiner (NIH grants R01HL130733);

Paul de Vries (American Heart Association grant 18CDA34110116);

Elizabeth C Oelsner (NHLBI Pooled Cohorts Study and NIH grants R21-HL129924, K23**-**HL130627);

Santhi Ganesh (NIH grants R01HL122684, R01HL139672); Brian E. Cade (NIH grant K01-HL135405);

Pradeep Natarajan and Gina Peloso (NIH grant R01HL142711);

Dr. Vasan is supported in part by the Evans Medical Foundation and the Jay and Louis Coffman Endowment from the Department of Medicine, Boston University School of Medicine.

We acknowledge Drs. Mike Feolo and Masato Kimura for making the harmonized phenotype data and the phenotype tagging data available on dbGaP. We also acknowledge contributors to the overall TOPMed project, which can be found on the DCC website (https://www.nhlbiwgs.org/topmed-banner-authorship).

The views expressed in this manuscript are those of the authors and do not necessarily represent the views of the National Heart, Lung, and Blood Institute; the National Institutes of Health; or the U.S. Department of Health and Human Services.

Three authors report conflicts of interest:

Bruce Psaty serves on the Steering Committee of the Yale Open Data Access Project funded by Johnson & Johnson.

Pradeep Natarajan received grant support from Amgen, Apple, and Boston Scientific, and consulting fees from Apple, all unrelated to the present work.

Stella Aslibekyan is currently employed by and holds equity in 23andMe, Inc.

## ABBREVIATIONS

DCC: Data Coordinating Center
dbGaP: database for Genotypes and Phenotypes
JSON: JavaScript Object Notation
LDL-C: Low-density lipoprotein cholesterol
NHLBI: National Heart, Lung, and Blood Institute
NIH: National Institutes of Health
QC: quality control
TOPMed: Trans-Omics for Precision Medicine
UMLS: Unified Medical Language System
WG: Working Group

