## Supplementary material for "A system for phenotype harmonization in the NHLBI Trans-Omics for Precision Medicine (TOPMed) Program": Web Appendix

[1]Department of Biostatistics, School of Public Health, University of Washington, Seattle, Washington [2]Department of Medicine, School of Medicine, University of Alabama at Birmingham, Birmingham, Alabama [3]Data Management, Framingham Heart Study, Framingham, Massachusetts [4]Genetic Data Management, Framingham Heart Study, Framingham, Massachusetts [5]Department of Medicine, School of Medicine, Boston University, Boston, Massachusetts [6]Division of Public Health Sciences, Fred Hutchinson Cancer Research Center, Seattle, Washington [7]Division of Preventive Medicine, School of Medicine, University of Alabama at Birmingham, Birmingham, Alabama [8]Department of Genetics, School of Medicine, University of North Carolina at Chapel Hill, Chapel Hill, North Carolina [9]Cardiovascular Health Research Unit, Department of Medicine, University of Washington, Seattle, Washington [10]Human Genetics Center, Department of Epidemiology, School of Public Health, The University of Texas Health Science Center at Houston, Houston, Texas [11]Department of Epidemiology, School of Public Health and Tropical Medicine, Tulane University, New Orleans, Louisiana [12]Cardiovascular Research Center, Massachusetts General Hospital, Boston, Massachusetts [13]Center for Genomic Medicine, Massachusetts General Hospital, Boston, Massachusetts [14]Department of Medicine, Harvard Medical School, Harvard University, Boston, Massachusetts [15]Program in Medical and Population Genetics, Broad Institute of MIT and Harvard, Cambridge, Massachusetts [16]Department of Biostatistics, School of Public Health, Boston University, Boston, Massachusetts [17]Department of Epidemiology, School of Public Health, University of Michigan, Ann Arbor, Michigan [18]Department of Epidemiology, School of Public Health, University of Washington, Seattle, Washington [19]Fred Hutchinson Cancer Research Center, Seattle, Washington [20]Dean's Office, University of Kentucky, Lexington, Kentucky [21]Department of Epidemiology, School of Public Health, University of Alabama at Birmingham, Birmingham, Alabama [22]Colorado Center for Personalized Medicine, School of Medicine, University of Colorado, Aurora, Colorado [23]Division of Sleep and Circadian Disorders, Department of Medicine, Brigham and Women's Hospital, Boston, Massachusetts [24]Division of Sleep Medicine, Harvard Medical School, Harvard University, Boston, Massachusetts [25]National Heart, Lung, and Blood Institute, National Institutes of Health, Bethesda, Maryland [26]Framingham Heart Study, Framingham, Massachusetts [27]Division of Intramural Research, Population Sciences Branch, National Heart, Lung and Blood Institute, Bethesda [28]Department of Medicine, University of Mississippi Medical Center, Jackson, Mississippi [29]Department of Pediatrics, University of Mississippi Medical Center, Jackson, Mississippi [30]Department of Population Health Science, John D. Bower School of Population Health, University of Mississippi Medical Center, Jackson, Mississippi [31]Division of Biomedical Statistics and Informatics, Mayo Clinic, Rochester, MN, Rochester, Minnesota [32]Cardiovascular Disease Initiative, Broad Institute of MIT and Harvard, Cambridge, Massachusetts

[33]Brown Foundation Institute of Molecular Medicine, McGovern Medical School, The University of Texas Health Science Center at Houston, Houston, Texas [34]Department of Epidemiology, Gillings School of Global Public Health, University of North Carolina at Chapel Hill, Chapel Hill, North Carolina [35]Division Of Lung Diseases, National Heart, Lung, and Blood Institute, National Institutes of Health, Bethesda, Maryland [36]Division of Cardiovascular Medicine, Department of Internal Medicine, Michigan Medicine, University of Michigan, Ann Arbor, Michigan [37]Department of Human Genetics, Michigan Medicine, University of Michigan, Ann Arbor, Michigan [38]Department of Statistics and Operations Research, Technical University of Catalonia, Barcelona, Spain [39]Department of Epidemiology, Human Genetics and Environmental Sciences, School of Public Health, The University of Texas Health Science Center at Houston, Houston, Texas [40]The Institute for Translational Genomics and Population Sciences, Department of Pediatrics, The Lundquist Institute for Biomedical Innovation at Harbor-UCLA Medical Center, Torrance, California [41]Department of Chronic Disease Epidemiology, Yale School of Public Health, New Haven, Connecticut [42]Division of Endocrinology, Diabetes and Metabolism, Department of Internal Medicine, The Ohio State University, Columbus, Ohio [43]Department of Epidemiology, Vitalant Research Institute, San Francisco, California [44]UCSF Benioff Children's Hospital Oakland, Oakland, California [45]GeneSTAR Research Center, Division of General Internal Medicine, School of Medicine, Johns Hopkins University, Baltimore, Maryland [46]International Health Institute, School of Public Health, Brown University, Providence, Rhode Island [47]Department of Epidemiology, School of Public Health, Brown University, Providence, Rhode Island [48]Division of Endocrinology, Diabetes and Nutrition and Program for Personalized and Genomic Medicine, Department of Medicine, University of Maryland School of Medicine, Baltimore, Maryland [49]Geriatrics Research and Education Clinical Center, Baltimore VA Medical Center, Baltimore, Maryland [50]Department of Medicine, School of Medicine, University of Pittsburgh, Pittsburgh, Pennsylvania [51]Department of Medicine, Columbia University, New York, New York [52]Department of Laboratory Medicine and Pathology, University of Minnesota Medical School, Minneapolis, Minnesota [53]Center for Public Health Genomics, School of Medicine, University of Virginia, Charlottesville, Virginia [54]Department of Public Health Sciences, School of Medicine, University of Virginia, Charlottesville, Virginia [55]Survey Research Center, Institute for Social Research, University of Michigan, Ann Arbor, Michigan [56]Department of Epidemiology, School of Public Health, Boston University, Boston, Massachusetts [57]Department of Human Genetics, Graduate School of Public Health, University of Pittsburgh, Pittsburgh, Pennsylvania [58]Department of Biostatistics, Graduate School of Public Health, University of Pittsburgh, Pittsburgh, Pennsylvania [59]Channing Division of Network Medicine, Department of Medicine, Brigham and Women's Hospital, Boston, Massachusetts [60]Harvard Medical School, Harvard University, Boston, Massachusetts [61]Research Informatics Services, National Jewish Health, Denver, Colorado [62]Department of Health Services, School of Public Health, University of Washington, Seattle, Washington [63]Kaiser Permanente Washington Health Research Institute, Seattle, Washington

**S4 Updating harmonized variables** **32**

**S5 Distributing harmonization results to the scientific community** **32**

**S6 Harmonized phenotype documentation and reproducibility** **32**

**S7 Phenotype tagging detailed methods** **37**

**References** **46**

Table S1: The studies included in the DCC's harmonized phenotypes.

| Study | Name | dbGaP TOPMed accession | dbGaP phenotype accession(s) |
|---|---|---|---|
| Amish | Genetics of Cardiometabolic Health in the Amish | phs000956 | phs000956 |
| ARIC | Atherosclerosis Risk in Communities Study | phs001211 | phs000280 |
| CARDIA | Coronary Artery Risk Development in Young Adults | phs001612 | phs000285 |
| CFS | Cleveland Family Study | phs000954 | phs000284 |
| CHS | Cardiovascular Health Study | phs001368 | phs000287 |
| COPDGene | Genetic Epidemiology of COPD Study | phs000951 | phs000179 |
| CRA | The Genetic Epidemiology of Asthma in Costa Rica | phs000988 | phs000988 |
| FHS | Framingham Heart Study | phs000974 | phs000007 |
| GENOA | Genetic Epidemiology Network of Arteriopathy | phs001345 | phs001238 |
| GOLDN | Genetics of Lipid Lowering Drugs and Diet Network | phs001359 | phs000741 |
| HCHS_SOL | Hispanic Community Health Study - Study of Latinos | phs001395 | phs000810 |
| HVH | Heart and Vascular Health Study | phs000993 | phs001013 |
| JHS | Jackson Heart Study | phs000964 | phs000286 |
| Mayo_VTE | Mayo Clinic Venous Thromboembolism Study | phs001402 | phs000289, phs001402 |
| MESA | Multi-Ethnic Study of Atherosclerosis | phs001416 | phs000209 |
| Samoan | Samoan Adiposity Study | phs000972 | phs000914 |
| WHI | Women's Health Initiative | phs001237 | phs000200 |

### S1 TOPMed study data on dbGaP

The National Heart, Lung and Blood Institute's (NHLBI) TOPMed program is providing genomic data for over 80 studies that had previously recruited participants and collected phenotypic data. For many of these studies, phenotypic data were previously available in the database for Genotypes and Phenotypes (dbGaP; https://www.ncbi.nlm.nih.gov/gap/) for controlled-access by the scientific community (along with other non-TOPMed genomic data). For these studies, phenotype data are largely included in the pre-existing 'parent' study accession on dbGaP, while the TOPMed genomic data are currently in a separate 'TOPMed' accession (although there is a plan to eventually merge the two accessions). Some studies did not have prior data in dbGaP and, in these cases, both phenotypic and TOPMed genomic data are in the TOPMed accession. Web Table S1 provides TOPMed and parent study accessions for the studies used in the harmonizations reported in this paper.

The TOPMed Data Coordinating Center (DCC) harmonization process uses phenotype variables submitted by studies to dbGaP as the source data for harmonization. The provenance of harmonized variables is tracked through accession identifiers assigned by dbGaP. These include unique identifiers for each study ("phs" prefix), phenotypic data set within study ("pht") and variable within data set ("phv") (1). A given study, phenotypic data set, or variable accession can be found on the dbGaP website by searching for the accession number (e.g., phs000007), and then selecting the record for that accession in the appropriate results tab.

### S2 Phenotype database overview

The TOPMed DCC's relational database stores both study phenotype data, which are used as components in harmonization, and harmonized data. The database contains two central tables that store phenotype data values in "entity-attribute-value" format, one for study data and the other for harmonized data. In these tables, "entity" is an internal participant identifier (unique within and among studies), "attribute" is a unique phenotype variable identifier, and "value" is the value of the attribute for a given participant. For the harmonized data values, we also track age at measurement in this table. The internal participant identifier is called "topmed_subject_id" and is used when processing study data during harmonization. Additional database tables track metadata associated with the actual data values, including provenance and other information needed for documentation of harmonized phenotype values. We currently use MariaDB version 10.2.11 (2).

### S3 Harmonization Steps

In this section, we supplement the main text with more details about the DCC's harmonization process and illustrate the steps using four phenotype variables: ever_smoker_baseline_1 (ever-smoker status), bp_systolic_1 (systolic blood pressure, SBP), il6_1 (interleukin 6 concentration in blood), and cimt_2 (common carotid intima media thickness, cIMT).

#### S3.1 Step 1: Define the harmonized phenotype variable

The following examples provide definitions of the target harmonized variable, which include units for continuous variables and definitions of encoded values for categorical variables. As noted below, some of the variable definitions were modified during the harmonization process. Each harmonized variable is given an intermediate name, indicating the phenotype concept and sometimes a modifier related to time point (or some other feature). This intermediate name is then converted into the final name by appending a concept variant number to differentiate among different implementations of harmonization for the same basic phenotype concept. For example, cimt_1 and cimt_2 are names for common carotid intima media thickness variables calculated with slightly different harmonization algorithms. Another example is "ever_smoker_baseline_1" to indicate ever smoker status assessed at the baseline clinic visit. Each harmonized phenotype variable is paired with age at measurement, assessment or biosample collection, with the exception of certain demographic variables (e.g., subcohort code within a study). The age variables are named by pre-appending "age_at_" to the harmonized variable name (e.g. "age_at_cimt_1").

##### S3.1.1 Step 1 example: ever_smoker_baseline_1

The harmonized "ever_smoker_baseline_1" variable was defined as an indicator of whether a participant had ever regularly smoked cigarettes prior to the time when they enrolled in the study. To determine whether smoking was "regular", we relied on wording provided in each study's assessment of smoking behavior. For example, one study phrased their question as: "have you ever smoked cigarettes regularly? (no means less than 20 packs of cigarettes or 12 oz. of tobacco in a lifetime or less than 1 cigarette a day for a year.)" ("G3A070"; phv00020925). Studies that did not ask specifically about regular smoking were excluded. The values of this variable were encoded as 0="never a cigarette smoker" and 1="current or former cigarette smoker".

##### S3.1.2 Step 1 example: bp_systolic_1

The harmonized "bp_systolic_1" variable was defined as SBP in units of mmHg, measured from the upper arm in a clinical setting while the participant is in a resting position.

The Working Group's (WG) analysis plan suggested that the SBP variable should be increased by a fixed amount (15 mmHg) if the participant had been taking blood pressure-lowering medication. This suggestion was accommodated by providing a separate harmonized variable for whether or not a participant was taking such medication, so that users can decide exactly how to account for medication usage.

##### S3.1.3 Step 1 example: il6_1

The harmonized variable il_6 is defined as the concentration of interleukin 6 (IL6) in pg/mL measured in either blood serum or plasma. The measurement methodology was specified as enzyme-linked immunosorbent assay. Participants whose assay measurements were marked as failed by the study were set to missing.

Study documentation about laboratory protocols for IL6 measurements indicated differences among studies in the upper and lower limits of detection (LOD) for their assays. Furthermore, studies handled values outside the LOD in various ways, including setting these values to the LOD plus or minus 1; setting them to the

LOD; providing an indicator variable; or providing no information about whether samples were outside the LOD. To harmonize these variations, participants with IL6 concentration values that were below or above the LOD were set to the lower LOD or upper LOD, respectively, in all studies.

###### S3.1.4 Step 1 example: cimt_2

The harmonized variable for common carotid intima media thickness (cIMT) was initially defined as the thickness of the common carotid intima media as measured using ultrasound.

Because of study differences in the availability of near and far wall measurements, we harmonized two different variables for cIMT; each was assigned a different concept variant number. The two definitions are as follows:

1. cimt_1: the mean of two values: mean of multiple thickness estimates from the left far wall and from the right far wall
2. cimt_2: the mean of four values: maximum of multiple thickness estimates from the left far wall, left near wall, right far wall and right near wall.

Studies that had taken measurements fitting both definitions were included in both variables, while those that had measurements for just one definition would only be included in that one variable. The choice of two cIMT variables allows analysts to use their preferred definition in their analysis, or even to combine values from the two variables if they desire.

In the following examples for cIMT, we focus on the cimt_2 variable.

##### S3.2 Step 2: Identify candidate phenotype variables across contributing studies

The main task in this step is to find the candidate variables for the phenotype being harmonized. The lack of controlled vocabulary applied to study variables on dbGaP means that manual searches for relevant keywords are required to obtain a full set of candidate variables to consider for inclusion in harmonization. We have implemented a number of strategies to accomplish this task, such as contacting WG and study representatives for more information about the data, searching and browsing the available data using an internal web app, or using the results of the phenotype tagging project.

Analysts encounter various challenges when identifying candidate variables. One difficulty is cryptic variable names, which often use either some abbreviation of what's being measured (e.g., "cursmk1" for whether the participant smoked cigarettes in the last 30 days; phv00085572) or an administrative naming scheme to track the original questionnaire and field from which the variable was derived (e.g. "A99" for whether a participant smokes cigarettes; phv00007612). The process is also hampered by inconsistencies in variable descriptions across studies, which may use abbreviations (e.g., "AVZMSYS" contains systolic blood pressure but has a description "AVE ZERO MUD SYSTOL"; phv00100435), use synonyms for the same phenotype in different data sets (e.g., "sugar" vs. "glucose" for blood glucose measurements in "MF65" and "GLUSIU21"; phv00000537 and phv00204643), or give incomplete information (e.g., the variable description for "MF256" consists only of "4"; phv00000711). Organizational differences between studies can also add complexity to identifying potential component variables. For example, some datasets use an entity-attribute-value structure to report phenotype datasets, which means that the contents of a variable must be examined before determining whether it contains relevant phenotype data (e.g., variables "TESTNAME" and "TESTVAL" for laboratory test names and values; phv00282937 and phv00193862).

Studies may have measured the same phenotype multiple times in the same participant, either as repeated measures at the same timepoint or longitudinal measurements collected over many years. Analysts must decide how to handle these multiple measurements. At the DCC, this also means choosing a single measurement for each participant due to analysis models commonly used in genetic association testing, which are generally run without repeated measures. To select the timepoint to use for a given phenotype, DCC analysts weigh multiple factors. One consideration is how the study protocol for collecting each measurement compares with that used for other studies. In some cases, more recent measurements may be chosen to reduce heterogeneity among studies, as earlier measurements are more likely to have been collected using older protocols that differed from more modern methods. A second option is to choose the timepoint with the largest set of non-missing values to provide larger power in analyses. The specific strategy chosen depends on the intended analysis plan for the harmonized variable.

After selecting the timepoint to include in harmonization, analysts must identify variables that contain the age of measurement of the phenotype at the chosen timepoint. This process is highly dependent on each study's organization strategy. The most common strategy used by studies is to provide age at a given exam as a separate variable that applies to all other variables from that dataset (e.g., "AGEBL"; phv00100487). A related case is to provide an age variable that applies to all other variables from a given exam (e.g., "age1"; phv00177930). A more complicated strategy is to provide an age at a given point (e.g., "AGE"; phv00078437) plus the time from that date to the date of measurement (e.g., "F2DAYS" and "F34DAYS"; phv00078436 and phv00078773); these three variables must be used together to calculate the appropriate age at measurement.

In some cases, studies provide variables for both the original data values and for a derived variable calculated from the original values. Studies can calculate these derived variables differently, or the derived variables may not match the original definition of the harmonized variable. When studies provide both the original and derived data values, we prefer to use the original data values to recalculate the derived variable. For example, when harmonizing low density lipoprotein cholesterol (LDL-C), we used total cholesterol, triglycerides, and high density lipoprotein cholesterol as component variables to compute harmonized LDL-C using the Friedewald equation (3) instead of using a study's derived LDL-C variable directly.

##### S3.2.1 Step 2 example: ever_smoker_baseline_1

For this phenotype, we searched phenotype variable descriptions from each study for terms related to smoking (e.g., "smok*", "cigarette", "tob*", etc.). We attempted to identify all variables indicating cigarette smoking status, which include both direct indicators such as "Have you ever smoked cigarettes?" ("EverSmokedCig"; phv00159747), as well as other indicators of lifetime smoking history such as "How old were you when you first started regular cigarette smoking?" ("H065"; phv00072093) or "On average how many cigarettes per day do/did you usually smoke?" ("AVGCIGDY"; phv00307905). Studies generally obtained this information via self-report using questionnaires. Variables were limited to those collected at the baseline visit for each study. We then identified variables related to the age at measurement of the "ever_smoker_baseline_1" variable.

We highlight some examples of the difficulty of identifying relevant candidate variables for ever_smoker_baseline_1. Some studies' questions did not exactly fit the definition of the harmonized phenotype and, when this occurs, analysts must make a decision about whether the variable is close enough to the definition to include. For example, in the Framingham Heart Study (FHS) "Original Cohort", we did not find an ever-smoker question specifically about cigarette smoking, but did find a more general one about tobacco use. Because of the importance of this subcohort, which was enrolled in 1948 (4), we decided to include this variable, considering that users may decide whether or not to include this sample set in their analyses. Finally, we note that some studies separate data for different participants into different datasets (e.g., repeated "evsmk1" variables from different datasets in MESA; phv00083243 and phv00085570), which both need to be processed to derive a harmonized variable that includes all study participants.

The study variables selected for inclusion in the ever_smoker_baseline_1 variable harmonization after quality control (QC) are shown in WebTable S2 and for the associated age variable in Web Table S3.

##### S3.2.2 Step 2 example: bp_systolic_1

For this variable, we searched for phenotype variable descriptions using keywords like "bp", "systol*", "sys*", and "sbp". Due to the paired nature of the systolic and diastolic measurements, analysts also identified variables that measured diastolic blood pressure (DBP) at the same timepoint, which was done using additional keywords such as "diastol*", "dia*", and "dbp". We also looked for dataset names and descriptions using similar keywords.

In variables identified in these searches, we found that several types of instruments were used to make blood pressure measurements. After consultation with members in the Blood Pressure WG, a decision was made to only include variables with data collected using some type of sphygmomanometer. When a random-zero sphygmomanometer was used and the zero readings were available in dbGaP, the zero reading adjustments were applied in the harmonization.

We selected measurements from the baseline visit for most studies. A small number of studies did not have information about antihypertensive medication use at the baseline exam, so in these cases, measurements from the earliest exam with both blood pressure measurements and information about antihypertensive medication use were chosen.

Some studies provide each blood pressure measurement from a repeat set of measurements as separate variables (e.g., “SBPA13”, phv00128370; “SBPA16”, phv00128373), while others provide only an average that they computed (e.g., “Systolic_BP”, phv00258701). When possible, we recalculated the average using the individual measurements, but we used the average if it was the only variable available for a given study.

Two studies provided only one blood pressure measurement instead of a repeated set of measures or the average of those measures. Even though the original definition of the variable required that bp_systolic_1 be calculated as an average of two measures, we decided to include these studies in the harmonized variable to increase the sample size. The difference from the original definition was noted in the harmonization comments so that blood pressure measurements from these studies can be excluded if desired.

The study variables selected for inclusion in the bp_systolic_1 variable harmonization after QC are shown in Web Table S4. This table includes both SBP and DBP variables (e.g., “Systolic_BP” and “Diastolic_BP”; phv00258701 and phv00258703) due to QC steps discussed in section S3.3.2.

###### S3.2.3 Step 2 example: il6_1

For this variable, DCC analysts searched for phenotype variable descriptions using keywords like “il6”, “il-6”, “il 6”, or “interleukin”. Because inflammation biomarkers are not a commonly-measured phenotype, variables from any visit were considered, but a single visit for each study or subcohort within a study was chosen such that it provided the maximum sample size for that study or subcohort. When repeated measures were made from sample(s) taken during a single visit, the first measurement was used for consistency with studies in which only one measurement was made.

In addition to the variables measuring IL6, supporting variables to indicate sample quality were also used for harmonization when available, such as an indicator of whether a sample assay failed or was outside the upper or lower LOD. These quality indicators may be difficult to find because they could have descriptions that are vague outside the context of the dataset or related study documentation (e.g., “flag”; phv00081000). Analysts typically identified these variables by reading study documentation and inspecting variables in the same dbGaP dataset as the IL6 measurement variables.

The study variables selected for inclusion in the il6_1 variable harmonization after QC are shown in Web Table S5 The components include the variables measuring IL6 as well as supporting variables about sample quality required for harmonization. Some studies (e.g., FHS) have multiple variables representing measurements from participants in different subcohorts.

###### S3.2.4 Step 2 example: cimt_2

For this variable, we searched for strings like “intima media thickness”, “imt”, “cimt”, “common carotid artery”, and “intima media thickness” in dbGaP study variable descriptions. To minimize heterogeneity within a given study or subcohort within a study, variables only from a specific visit for each study or subcohort were included. The visit used for each study or subcohort was chosen by reading study documentation and consulting studies for recommendations. Sample size and number of non-missing cIMT measurements were also taken into account when selecting the visit to use.

Some studies provided measurements of the common cIMT in specific regions of the carotid artery in separate variables, while others provided only the target cIMT quantity (i.e., mean-of-max cIMT). When possible, measurements of specific regions were used to derive the cIMT values instead of using study-derived mean-of-max cIMT values, but the study-derived variables were used if individual measurements were not available.

Most studies provided only one cIMT measurement for each participant at either systole or diastole, but one study provided cIMT measurements at both systole and diastole for some participants. After discussion with the study and informed research about the phenotype, we included the cIMT measurements at diastole in the harmonization to match previous investigations of cIMT (5). This decision was noted in the harmonization comments.

In the CHS study, ultrasound images were acquired for the Original subcohort in year 1 and for the New subcohort in year 5. We found two sets of cIMT measurements for the Original subcohort and, after consultation with the study, determined that this was due to the fact that the original images for this subcohort were reread in year 5 when the New subcohort ultrasounds were performed. In order to increase consistency across both baseline ultrasound readings, we used the reread measurements from year 5.

The study variables selected for inclusion in the cimt_2 variable harmonization after QC are shown in Web Table S6.

Table S2: Component study variables used to harmonize ever_smoker_baseline_1.

| Variable accession | Variable name | Variable description |
|---|---|---|
| **ARIC** | | |
| phs000280.v4.pht004111.v2.phv00207368.v1 | HOM28 | Have you ever smoked cigarettes? Q28 [Home Interview, exam 1] |
| phs000280.v4.pht004111.v2.phv00207369.v1 | HOM29 | How old were you when you first started regular cigarette smoking? Q29 [Home Interview, exam 1] |
| phs000280.v4.pht004111.v2.phv00207370.v1 | HOM30 | Do you now smoke cigarettes? Q30 [Home Interview, exam 1] |
| phs000280.v4.pht004111.v2.phv00207375.v1 | HOM35 | On the average of the entire time you smoked, how many cigarettes did you usually smoke per day? Q35 [Home Interview, exam 1] |
| phs000280.v4.pht004111.v2.phv00207376.v1 | HOM36 | (Do/did) you inhale the cigarette smoke? Q36 [Home Interview, exam 1] |
| **CARDIA** | | |
| phs000285.v3.pht001573.v2.phv00113213.v2 | A10CIGS | SUBJECT HAS SMOKED CIGARETTES. Q 2 |
| **CFS** | | |
| phs000284.v1.pht001902.v1.phv00122012.v1 | visit | Visit Number |
| phs000284.v1.pht001902.v1.phv00122340.v1 | SMOKED | Ever smoked cigarettes (A) |
| phs000284.v1.pht001902.v1.phv00122341.v1 | AGESMOK | Age when first smoked cigarettes (A) |
| phs000284.v1.pht001902.v1.phv00122342.v1 | AVGSMOK | Average number of cigarettes smoke per day (A) |
| phs000284.v1.pht001902.v1.phv00122343.v1 | MONSMOKE | Past month, smoke >=1 cigarettes/day (A) |
| phs000284.v1.pht001902.v1.phv00122344.v1 | NOWSMOKE | Number of cigarettes currently smoke/day (A) |
| **CHS** | | |
| phs000287.v6.pht001450.v1.phv00098844.v1 | SMOKE101 | SMOKED IN LIFETIME |
| phs000287.v6.pht001450.v1.phv00098845.v1 | SMOKE201 | SMOKED CIGARETTES LAST 30 DAYS |
| phs000287.v6.pht001450.v1.phv00099157.v1 | SMKAGE08 | HOW OLD WHEN YOU STARTED TO SMOKE |
| phs000287.v6.pht001450.v1.phv00099159.v1 | AMOUNT08 | HOW MANY DID YOU SMOKE PER DAY ON AVER (99=UNKNOWN) |
| phs000287.v6.pht001490.v1.phv00105143.v1 | SMOKE101 | SMOKED IN LIFETIME |
| phs000287.v6.pht001490.v1.phv00105144.v1 | SMOKE201 | SMOKED CIGARETTES LAST 30 DAYS |
| phs000287.v6.pht001490.v1.phv00106198.v1 | SMKAGE58 | HOW OLD WHEN YOU STARTED TO SMOKE |
| phs000287.v6.pht001490.v1.phv00106200.v1 | AMOUNT58 | HOW MANY DID YOU SMOKE PER DAY ON AVER. |
| **COPDGene** | | |
| phs000179.v5.pht002239.v4.phv00159636.v4 | HowSoonSmoke | How soon after waking do you smoke first cigarette |
| phs000179.v5.pht002239.v4.phv00159637.v4 | SmokeMore2hrs | Smoke more during first 2 hours of day than rest of day |
| phs000179.v5.pht002239.v4.phv00159638.v4 | CigHateGiveUp | Which cigarette would you hate most to give up |
| phs000179.v5.pht002239.v4.phv00159639.v4 | FindHardNotSmoke | Do you find it hard to not smoke in forbidden places |
| phs000179.v5.pht002239.v4.phv00159640.v4 | SmokeSickBed | Smoke when so ill you are in bed most of day |
| phs000179.v5.pht002239.v4.phv00159641.v4 | SmokeMenthol | Do you now or did you smoke menthol cigarettes |

Table S2: *(continued)*

| Study | Variable accession | Variable name | Variable description |
|---|---|---|---|
| | phs000179.v5.pht002239.v4.phv00159747.v4 | EverSmokedCig | Have you ever smoked cigarettes? |
| | phs000179.v5.pht002239.v4.phv00159748.v4 | SmokStartAge | How old were you when you first started cigarette smoking? [Years old] |
| | phs000179.v5.pht002239.v4.phv00159749.v4 | SmokCigNow | Do you now smoke cigarettes [as of one month ago]? |
| | phs000179.v5.pht002239.v4.phv00159750.v4 | CigPerDaySmokNow | How many cigarettes do you smoke per day now? [Cigarettes/day] |
| | phs000179.v5.pht002239.v4.phv00159752.v4 | CigPerDaySmokAvg | Average for entire time how many cigarettes smoked per day [cigarettes/day] |
| | phs000179.v5.pht002239.v4.phv00159754.v4 | CigSmok24hrs | How many cigarettes have you smoked in the past 24 hours [cigarettes] |
| | phs000179.v5.pht002239.v4.phv00159755.v4 | CigSmok2hrs | How many cigarettes have you smoked in the past 2 hours [cigarettes] |
| | phs000179.v5.pht002239.v4.phv00159756.v4 | CigSmokHalfHr | How many cigarettes have you smoked in the past half hour [cigarettes] |
| | phs000179.v5.pht002239.v4.phv00169388.v3 | Duration_Smoking | Duration of smoking [yrs] |
| **CRA** | | | |
| | phs000988.v2.pht005248.v2.phv00267374.v2 | ever_Smoker | Ever smoked |
| | phs000988.v2.pht005248.v2.phv00267375.v2 | Current_Smoker | Current smoking status |
| | phs000988.v2.pht005248.v2.phv00267376.v2 | former_Smoker | Former smoking status |
| | phs000988.v2.pht005248.v2.phv00267378.v2 | cigsperday | Number of cigarettes smoked per day |
| | phs000988.v2.pht005248.v2.phv00267379.v2 | cigsperday_average | Number of cigarettes smoked per day, averaged over all years of smoking |
| **FHS** | | | |
| | phs000007.v29.pht000009.v2.phv00000543.v1 | MF71 | TOBACCO USED "NOW" OR "EVER" |
| | phs000007.v29.pht000030.v7.phv00007612.v5 | A99 | SMOKES CIGARETTES |
| | phs000007.v29.pht000074.v10.phv00020925.v4 | G3A070 | HAVE YOU EVER SMOKED CIGARETTES REGULARLY? (NO MEANS LESS THAN |
| | phs000007.v29.pht006005.v1.phv00273759.v1 | g3a070 | Have you ever smoked cigarettes regularly? (no means less than 20 packs of cigarettes or 12 oz. of tobacco in a lifetime or less than 1 cigarette a day for a year.) |
| | phs000007.v29.pht006006.v1.phv00274252.v1 | g3a070 | Have you ever smoked cigarettes regularly? (no means less than 20 packs of cigarettes or 12 oz. of tobacco in a lifetime or less than 1 cigarette a day for a year.) |
| **GENOA** | | | |
| | phs001238.v1.pht006043.v1.phv00277618.v1 | SMOKE100 | Have you smoked more than 100 cigarettes in your entire life? |
| | phs001238.v1.pht006043.v1.phv00277621.v1 | CIGARETT | Do you now smoke cigarettes? |
| | phs001238.v1.pht006043.v1.phv00277624.v1 | AVGCIGDY | On average how many cigarettes per day do/did you usually smoke? |
| | phs001238.v1.pht006657.v1.phv00307899.v1 | SMOKE100 | Have you smoked more than 100 cigarettes in your entire life? |
| | phs001238.v1.pht006657.v1.phv00307902.v1 | CIGARETT | Do you now smoke cigarettes? |
| | phs001238.v1.pht006657.v1.phv00307905.v1 | AVGCIGDY | On average how many cigarettes per day do/did you usually smoke? |
| **HCHS_SOL** | | | |
| | phs000810.v1.pht004715.v1.phv00258106.v1 | TBEA1 | Smoke at least 100 cigs in lifetime (TBEA1) |
| | phs000810.v1.pht004715.v1.phv00258107.v1 | TBEA3 | Present smoking status (TBEA3) |
| | phs000810.v1.pht004715.v1.phv00258108.v1 | TBEA4 | Daily: cigs per day - present (TBEA4) |
| | phs000810.v1.pht004715.v1.phv00258110.v1 | TBEA5A | Some: cigarettes per day on days you smoked during past 30 days - original desc: some: past 30 days - quit smoking 6 months or longer (TBEA5A) |
| **HVH** | | | |
| | phs001013.v3.pht005311.v2.phv00259376.v2 | ccs | Case-control status |
| | phs001013.v3.pht005311.v2.phv00259377.v2 | indexy | Index Year |
| | phs001013.v3.pht005311.v2.phv00259394.v2 | smoke | Smoking status at index date |
| **JHS** | | | |
| | phs000286.v5.pht001977.v1.phv00128496.v1 | TOBA1 | 1: Smoked at least 400 cigarettes |
| | phs000286.v5.pht001977.v1.phv00128498.v1 | TOBA3 | 3: Do you now smoke cigarettes |
| | phs000286.v5.pht001977.v1.phv00128502.v1 | TOBA6 | 6: Smoke more first few hrs after wake |
| | phs000286.v5.pht001977.v1.phv00128503.v1 | TOBA7 | 7: How soon do you smoke? |
| | phs000286.v5.pht001977.v1.phv00128506.v1 | TOBA10 | 10: Smoke when ill? |
| | phs000286.v5.pht001977.v1.phv00128507.v1 | TOBA11 | 11: Cigarettes smoke usually per day |
| **MESA** | | | |

Table S2: *(continued)*

| Study | Variable accession | Variable name | Variable description |
|---|---|---|---|
| | phs000209.v13.pht001111.v4.phv00083243.v1 | evsmk1 | SMOKED AT LEAST 100 CIGARETTES IN LIFETIME |
| | phs000209.v13.pht001111.v4.phv00083245.v1 | cursmk1 | CIGARETTES: SMOKED IN LAST 30 DAYS |
| | phs000209.v13.pht001111.v4.phv00083247.v1 | cigsday1 | CIGARETTES: AVERAGE # SMOKED PER DAY |
| | phs000209.v13.pht001116.v10.phv00085570.v2 | evsmk1 | SMOKED AT LEAST 100 CIGARETTES IN LIFETIME |
| | phs000209.v13.pht001116.v10.phv00085572.v2 | cursmk1 | CIGARETTES: SMOKED IN LAST 30 DAYS |
| | phs000209.v13.pht001116.v10.phv00085574.v2 | cigsday1 | CIGARETTES: AVERAGE # SMOKED PER DAY |
| | phs000209.v13.pht001121.v3.phv00087252.v1 | evsmkf | SMOKED 100+ CIGARETTES IN LIFETIME |
| | phs000209.v13.pht001121.v3.phv00087254.v1 | cursmkf | SMOKED CIGARETTES IN THE LAST 30 DAYS |
| | phs000209.v13.pht001121.v3.phv00087256.v1 | cigsdayf | AVERAGE NUMBER OF CIGARETTES SMOKED PER DAY |
| **Samoan** | | | |
| | phs000914.v1.pht005253.v1.phv00258705.v1 | Current_smoke | Current Smoker |
| | phs000914.v1.pht005253.v1.phv00258713.v1 | Past_smoker | Past Smoker |
| | phs000200.v11.pht001003.v6.phv00078774.v6 | SMOKEVR | Smoked at least 100 cigarettes ever |

Table S3: Component study variables used to harmonize age at measurement of ever_smoker_baseline_1.

| | Variable accession | Variable name | Variable description |
|---|---|---|---|
| **ARIC** | | | |
| | phs000280.v4.pht004063.v2.phv00204712.v1 | V1AGE01 | Age at visit 1 [Cohort, Exam 1] |
| **CARDIA** | | | |
| | phs000285.v3.pht001559.v2.phv00112439.v2 | A01AGE2 | AGE VERIFY |
| **CFS** | | | |
| | phs000284.v1.pht001902.v1.phv00122015.v1 | age | Subject age at time of study |
| **CHS** | | | |
| | phs000287.v6.pht001452.v1.phv00100487.v1 | AGEBL | CALCULATED AGE AT BASELINE |
| **COPDGene** | | | |
| | phs000179.v5.pht002239.v4.phv00159836.v4 | Age_Enroll | Age at enrollment |
| **CRA** | | | |
| | phs000988.v2.pht005248.v2.phv00258650.v2 | age | Subject age |
| **FHS** | | | |
| | phs000007.v29.pht003099.v4.phv00177930.v4 | age1 | Age at Exam 1 |
| **GENOA** | | | |
| | phs001238.v1.pht006039.v1.phv00277507.v1 | AGE | Age at time of examination in years |
| | phs001238.v1.pht006653.v1.phv00307788.v1 | AGE | Age at time of examination in years |
| **HCHS_SOL** | | | |
| | phs000810.v1.pht004715.v1.phv00226251.v1 | AGE | Age |
| **HVH** | | | |
| | phs001013.v3.pht005311.v2.phv00259378.v2 | age | Age at index date |
| **JHS** | | | |
| | phs000286.v5.pht001949.v1.phv00126009.v1 | AGE01 | Age(yrs) at baseline clinic visit |
| **MESA** | | | |
| | phs000209.v13.pht001111.v4.phv00082639.v2 | age1c | AGE |
| | phs000209.v13.pht001116.v10.phv00084442.v3 | age1c | AGE |
| | phs000209.v13.pht001121.v3.phv00087071.v1 | agefc | AGE |
| **Samoan** | | | |
| | phs000914.v1.pht005253.v1.phv00258680.v1 | Dec_Age | Age at enrollment |
| **WHI** | | | |
| | phs000200.v11.pht000998.v6.phv00078436.v6 | F2DAYS | F2 Days since randomization |
| | phs000200.v11.pht000998.v6.phv00078437.v6 | AGE | Age at screening |
| | phs000200.v11.pht001003.v6.phv00078773.v6 | F34DAYS | F34 Days since randomization/enrollment |

Table S4: Component study variables used to harmonize bp_systolic_1.

| Variable accession | Variable name | Variable description |
|---|---|---|
| **Amish** | | |
| phs000956.v2.pht005002.v1.phv00252995.v1 | sbp__baseline | Systolic blood pressure at baseline visit |
| phs000956.v2.pht005002.v1.phv00252996.v1 | dbp__baseline | Diastolic blood pressure at baseline visit |
| **ARIC** | | |
| phs000280.v4.pht004192.v2.phv00210284.v1 | SBPA15 | [Second blood pressure measurement]. 2nd systolic. Q15 [Siting Blood Pressure, exam 1] |
| phs000280.v4.pht004192.v2.phv00210285.v1 | SBPA16 | [Second blood pressure measurement]. 2nd diastolic. Q16 [Siting Blood Pressure, exam 1] |
| phs000280.v4.pht004192.v2.phv00210286.v1 | SBPA17 | [Second blood pressure measurement]. 2nd zero reading. Q17 [Siting Blood Pressure, exam 1] |
| phs000280.v4.pht004192.v2.phv00210287.v1 | SBPA18 | [Third blood pressure measurement]. 3rd systolic. Q18 [Siting Blood Pressure, exam 1] |
| phs000280.v4.pht004192.v2.phv00210288.v1 | SBPA19 | [Third blood pressure measurement]. 3rd diastolic. Q19 [Siting Blood Pressure, exam 1] |
| phs000280.v4.pht004192.v2.phv00210289.v1 | SBPA20 | [Third blood pressure measurement]. 3rd zero reading. Q20 [Siting Blood Pressure, exam 1] |
| **CARDIA** | | |
| phs000285.v3.pht001560.v2.phv00112481.v2 | A02R2S | SECOND READING SBP |
| phs000285.v3.pht001560.v2.phv00112482.v2 | A02R2D | SECOND READING DBP |
| phs000285.v3.pht001560.v2.phv00112483.v2 | A02RZ2S | RZ2 SBP |
| phs000285.v3.pht001560.v2.phv00112484.v2 | A02RZ2D | RZ2 DBP |
| phs000285.v3.pht001560.v2.phv00112487.v2 | A02R3S | THIRD READING SBP |
| phs000285.v3.pht001560.v2.phv00112488.v2 | A02R3D | THIRD READING DBP |
| phs000285.v3.pht001560.v2.phv00112489.v2 | A02RZ3S | RZ3 SBP |
| phs000285.v3.pht001560.v2.phv00112490.v2 | A02RZ3D | RZ3 DBP |
| **CFS** | | |
| phs000284.v1.pht001902.v1.phv00122012.v1 | visit | Visit Number |
| phs000284.v1.pht001902.v1.phv00123001.v1 | sbp | Mean Systolic BP |
| phs000284.v1.pht001902.v1.phv00123002.v1 | dbp | Mean Diastolic BP |
| **CHS** | | |
| phs000287.v6.pht001452.v1.phv00100435.v1 | AVZMSYS | AVE ZERO MUD SYSTOL (mm Hg) |
| phs000287.v6.pht001452.v1.phv00100436.v1 | AVZMDIA | AVE ZERO MUD DIASTOL-adj (mm Hg) |
| **COPDGene** | | |
| phs000179.v5.pht002239.v4.phv00159583.v4 | diasBP | Diastolic blood pressure [mmHg] |
| phs000179.v5.pht002239.v4.phv00159590.v4 | sysBP | Systolic blood pressure [mmHg] |
| **FHS** | | |
| phs000007.v29.pht000009.v2.phv00000719.v1 | MF264 | BLOOD PRESSURE: FIRST EXAMINER, SYSTOLIC, EXAM 4 |
| phs000007.v29.pht000009.v2.phv00000720.v1 | MF265 | BLOOD PRESSURE: FIRST EXAMINER, DIASTOLIC, EXAM 4 |
| phs000007.v29.pht000009.v2.phv00000721.v1 | MF266 | BLOOD PRESSURE: SECOND EXAMINER, SYSTOLIC, EXAM 4 |
| phs000007.v29.pht000009.v2.phv00000722.v1 | MF267 | BLOOD PRESSURE: SECOND EXAMINER, DIASTOLIC, EXAM 4 |
| phs000007.v29.pht004813.v1.phv00250561.v1 | e485 | Physical Exam - Physician Blood Pressure First Reading - Systolic (nearest 2mm Hg) |
| phs000007.v29.pht004813.v1.phv00250562.v1 | e486 | Physical Exam - Physician Blood Pressure First Reading - Diastolic (nearest 2mm Hg) |
| phs000007.v29.pht004813.v1.phv00250652.v1 | e581 | Physical Exam - Physician Blood Pressure Second Reading - Systolic (nearest 2mm Hg) |
| phs000007.v29.pht004813.v1.phv00250653.v1 | e582 | Physical Exam - Physician Blood Pressure Second Reading - Diastolic (nearest 2mm Hg) |
| phs000007.v29.pht006026.v1.phv00277034.v1 | DBP1 | Average diastolic blood pressure, Exam 1 |
| phs000007.v29.pht006026.v1.phv00277045.v1 | SBP1 | Average systolic blood pressure, Exam 1 |
| phs000007.v29.pht006027.v1.phv00277137.v1 | DBP1 | Average diastolic blood pressure, Exam 1 |
| phs000007.v29.pht006027.v1.phv00277185.v1 | SBP1 | Average systolic blood pressure, Exam 1 |
| **GENOA** | | |
| phs001238.v1.pht006039.v1.phv00277520.v1 | RAND_SYS2 | Random-zero sphygmomanometer: Systolic; 2nd of 3 readings |
| phs001238.v1.pht006039.v1.phv00277521.v1 | RAND_DIA2 | Random-zero sphygmomanometer: Diastolic; 2nd of 3 readings |
| phs001238.v1.pht006039.v1.phv00277522.v1 | RAND_SYS3 | Random-zero sphygmomanometer: Systolic; 3rd of 3 readings |

Table S4: *(continued)*

| Study | Variable accession | Variable name | Variable description |
|---|---|---|---|
| | phs001238.v1.pht006039.v1.phv00277523.v1 | RAND_DIA3 | Random-zero sphygmomanometer: Diastolic; 3rd of 3 readings |
| | phs001238.v1.pht006653.v1.phv00307801.v1 | RAND_SYS2 | Random-zero sphygmomanometer: Systolic; 2nd of 3 readings |
| | phs001238.v1.pht006653.v1.phv00307802.v1 | RAND_DIA2 | Random-zero sphygmomanometer: Diastolic; 2nd of 3 readings |
| | phs001238.v1.pht006653.v1.phv00307803.v1 | RAND_SYS3 | Random-zero sphygmomanometer: Systolic; 3rd of 3 readings |
| | phs001238.v1.pht006653.v1.phv00307804.v1 | RAND_DIA3 | Random-zero sphygmomanometer: Diastolic; 3rd of 3 readings |
| **GOLDN** | | | |
| | phs000741.v2.pht003918.v2.phv00259052.v1 | SBP | Systolic Blood pressure |
| | phs000741.v2.pht003918.v2.phv00259053.v1 | DBP | Diastolic blood pressure |
| **HCHS_SOL** | | | |
| | phs000810.v1.pht004715.v1.phv00226390.v1 | SBPA5 | Average systolic blood pressure (SBPA5) |
| | phs000810.v1.pht004715.v1.phv00226391.v1 | SBPA6 | Average diastolic blood pressure (SBPA6) |
| **JHS** | | | |
| | phs000286.v5.pht001974.v1.phv00128370.v1 | SBPA13 | 13: Systolic (first BP) |
| | phs000286.v5.pht001974.v1.phv00128371.v1 | SBPA14 | 14: Diastolic (first BP) |
| | phs000286.v5.pht001974.v1.phv00128372.v1 | SBPA15 | 15: Zero reading (first BP) |
| | phs000286.v5.pht001974.v1.phv00128373.v1 | SBPA16 | 16: Systolic (second BP) |
| | phs000286.v5.pht001974.v1.phv00128374.v1 | SBPA17 | 17: Diastolic (second BP) |
| | phs000286.v5.pht001974.v1.phv00128375.v1 | SBPA18 | 18: Zero Reading (second BP) |
| **MESA** | | | |
| | phs000209.v13.pht001111.v4.phv00083403.v1 | s2bp1 | SEATED BP: SYSTOLIC 2ND READING (mmHg) |
| | phs000209.v13.pht001111.v4.phv00083404.v1 | d2bp1 | SEATED BP: DIASTOLIC 2ND READING (mmHg) |
| | phs000209.v13.pht001111.v4.phv00083406.v1 | s3bp1 | SEATED BP: SYSTOLIC 3RD READING (mmHg) |
| | phs000209.v13.pht001111.v4.phv00083407.v1 | d3bp1 | SEATED BP: DIASTOLIC 3RD READING (mmHg) |
| | phs000209.v13.pht001116.v10.phv00085735.v2 | s2bp1 | SEATED BP: SYSTOLIC 2ND READING (mmHg) |
| | phs000209.v13.pht001116.v10.phv00085736.v2 | d2bp1 | SEATED BP: DIASTOLIC 2ND READING (mmHg) |
| | phs000209.v13.pht001116.v10.phv00085737.v2 | s3bp1 | SEATED BP: SYSTOLIC 3RD READING (mmHg) |
| | phs000209.v13.pht001116.v10.phv00085738.v2 | d3bp1 | SEATED BP: DIASTOLIC 3RD READING (mmHg) |
| | phs000209.v13.pht001121.v3.phv00087509.v1 | s2bpf | 2ND READING: SEATED SYSTOLIC BP (mmHg) |
| | phs000209.v13.pht001121.v3.phv00087510.v1 | d2bpf | 2ND READING: SEATED DIASTOLIC BP (mmHg) |
| | phs000209.v13.pht001121.v3.phv00087512.v1 | s3bpf | 3RD READING: SEATED SYSTOLIC BP (mmHg) |
| | phs000209.v13.pht001121.v3.phv00087513.v1 | d3bpf | 3RD READING: SEATED DIASTOLIC BP (mmHg) |
| **Samoan** | | | |
| | phs000914.v1.pht005253.v1.phv00258701.v1 | Systolic_BP | Systolic blood pressure (average of last two measurements) |
| | phs000914.v1.pht005253.v1.phv00258703.v1 | Diastolic_BP | Diastolic blood pressure (average of last two measurements) |
| **WHI** | | | |
| | phs000200.v11.pht001019.v6.phv00079850.v6 | F80VTYP | Visit Type |
| | phs000200.v11.pht001019.v6.phv00079852.v6 | F80DAYS | F80 Days since randomization/enrollment |
| | phs000200.v11.pht001019.v6.phv00079854.v6 | SYSTBP1 | Systolic blood pressure (1st reading) |
| | phs000200.v11.pht001019.v6.phv00079855.v6 | DIASBP1 | Diastolic blood pressure (1st reading) |
| | phs000200.v11.pht001019.v6.phv00079856.v6 | SYSTBP2 | Systolic blood pressure (2nd reading) |
| | phs000200.v11.pht001019.v6.phv00079857.v6 | DIASBP2 | Diastolic blood pressure (2nd reading) |

Table S5: Component study variables used to harmonize il6_1.

| | Variable accession | Variable name | Variable description |
|---|---|---|---|
| **CARDIA** | | | |
| | phs000285.v3.pht001862.v2.phv00121064.v2 | FL6IL6 | IL6 PG/ML |
| | phs000285.v3.pht001862.v2.phv00121065.v2 | FL6IL6CM | IL6 COMMENTS |
| **CFS** | | | |
| | phs000284.v1.pht001902.v1.phv00122012.v1 | visit | Visit Number |
| | phs000284.v1.pht001902.v1.phv00124021.v1 | il6am | Il6 am (pg/mL) |
| **CHS** | | | |

Table S5: *(continued)*

| Study | Variable accession | Variable name | Variable description |
|---|---|---|---|
| | phs000287.v6.pht001452.v1.phv00100500.v1 | IL6BL | IL-6 at baseline (pg/ml) |
| **FHS** | | | |
| | phs000007.v29.pht000161.v6.phv00023796.v5 | il6 | INTERLEUKIN-6 FROM SERUM (PG/ML) |
| | phs000007.v29.pht001043.v4.phv00080999.v3 | il6 | Interleukin-6 concentration |
| | phs000007.v29.pht001043.v4.phv00081000.v3 | flag | Data type indicator |
| | phs000007.v29.pht002891.v4.phv00172223.v4 | il6 | Interleukin-6 |
| **MESA** | | | |
| | phs000209.v13.pht001116.v10.phv00085009.v2 | il61 | INTERLEUKIN-6 (IL-6) (pg/mL) |
| | phs000209.v13.pht001116.v10.phv00085010.v2 | il61M | EXCEPTIONAL MISSING IL61 |

Table S6: Component study variables used to harmonize cimt_2.

| | Variable accession | Variable name | Variable description |
|---|---|---|---|
| **ARIC** | | | |
| | phs000280.v3.pht004207.v1.phv00211053.v1 | LOPAMX23 | Maximum near wall width, left common carotid: optimal angle [Ultrasound Derived Data, exam 1] |
| | phs000280.v3.pht004207.v1.phv00211054.v1 | LANAMX23 | Maximum near wall width, left common carotid: anterior angle [Ultrasound Derived Data, exam 1] |
| | phs000280.v3.pht004207.v1.phv00211055.v1 | LPOAMX23 | Maximum near wall width, left common carotid: posterior angle [Ultrasound Derived Data, exam 1] |
| | phs000280.v3.pht004207.v1.phv00211059.v1 | ROPAMX23 | Maximum near wall width, right common carotid: optimal angle [Ultrasound Derived Data, exam 1] |
| | phs000280.v3.pht004207.v1.phv00211060.v1 | RANAMX23 | Maximum near wall width, right common carotid: anterior angle [Ultrasound Derived Data, exam 1] |
| | phs000280.v3.pht004207.v1.phv00211061.v1 | RPOAMX23 | Maximum near wall width, right common carotid: posterior angle [Ultrasound Derived Data, exam 1] |
| | phs000280.v3.pht004207.v1.phv00211081.v1 | LOPAMX45 | Maximum far wall width, left common carotid: optimal angle [Ultrasound Derived Data, exam 1] |
| | phs000280.v3.pht004207.v1.phv00211082.v1 | LANAMX45 | Maximum far wall width, left common carotid: anterior angle [Ultrasound Derived Data, exam 1] |
| | phs000280.v3.pht004207.v1.phv00211083.v1 | LPOAMX45 | Maximum far wall width, left common carotid: posterior angle [Ultrasound Derived Data, exam 1] |
| | phs000280.v3.pht004207.v1.phv00211087.v1 | ROPAMX45 | Maximum far wall width, right common carotid: optimal angle [Ultrasound Derived Data, exam 1] |
| | phs000280.v3.pht004207.v1.phv00211088.v1 | RANAMX45 | Maximum far wall width, right common carotid: anterior angle [Ultrasound Derived Data, exam 1] |
| | phs000280.v3.pht004207.v1.phv00211089.v1 | RPOAMX45 | Maximum far wall width, right common carotid: posterior angle [Ultrasound Derived Data, exam 1] |
| **CHS** | | | |
| | phs000287.v6.pht001452.v1.phv00100290.v1 | PERSTAT | COHORT |
| | phs000287.v6.pht001473.v1.phv00101238.v1 | NMAX155 | BL REREAD NEAR WALL MAX, R. COMMON |
| | phs000287.v6.pht001473.v1.phv00101239.v1 | FMAX155 | BL REREAD FAR WALL MAX, R. COMMON |
| | phs000287.v6.pht001473.v1.phv00101250.v1 | NMAX555 | BL REREAD NEAR WALL MAX, L. COMMON |
| | phs000287.v6.pht001473.v1.phv00101251.v1 | FMAX555 | BL REREAD FAR WALL MAX, L. COMMON |
| | phs000287.v6.pht001473.v1.phv00101264.v1 | NMAX141 | YEAR 5 NEAR WALL MAX, R. COMMON |
| | phs000287.v6.pht001473.v1.phv00101265.v1 | FMAX141 | YEAR 5 FAR WALL MAX, R. COMMON |
| | phs000287.v6.pht001473.v1.phv00101276.v1 | NMAX541 | YEAR 5 NEAR WALL MAX, L. COMMON |
| | phs000287.v6.pht001473.v1.phv00101277.v1 | FMAX541 | YEAR 5 FAR WALL MAX, L. COMMON |
| **FHS** | | | |
| | phs000007.v29.pht000083.v6.phv00021728.v5 | CCD_MEMX | MEAN OF MAX IMT FOR BOTH LEFT AND RIGHT COMMON CAROTID ARTERIES IN DIASTOLE (MM) |
| **JHS** | | | |
| | phs000286.v5.pht001978.v1.phv00128541.v1 | lcl_mx45 | Left common lateral maximum far wall in millimeters |
| | phs000286.v5.pht001978.v1.phv00128542.v1 | lca_mx45 | Left common anterior maximum far wall in millimeters |
| | phs000286.v5.pht001978.v1.phv00128543.v1 | lcp_mx45 | Left common posterior maximum far wall in millimeters |
| | phs000286.v5.pht001978.v1.phv00128544.v1 | rcl_mx45 | Right common lateral maximum far wall in millimeters |
| | phs000286.v5.pht001978.v1.phv00128545.v1 | rca_mx45 | Right common anterior maximum far wall in millimeters |
| | phs000286.v5.pht001978.v1.phv00128546.v1 | rcp_mx45 | Right common posterior maximum far wall in millimeters |
| | phs000286.v5.pht001978.v1.phv00128561.v1 | lcl_mx23 | Left common lateral maximum near wall in millimeters |

Table S6: *(continued)*

| Study | Variable accession | Variable name | Variable description |
|---|---|---|---|
| | phs000286.v5.pht001978.v1.phv00128562.v1 | lca_mx23 | Left common anterior maximum near wall in millimeters |
| | phs000286.v5.pht001978.v1.phv00128563.v1 | lcp_mx23 | Left common posterior maximum near wall in millimeters |
| | phs000286.v5.pht001978.v1.phv00128564.v1 | rcl_mx23 | Right common lateral maximum near wall in millimeters |
| | phs000286.v5.pht001978.v1.phv00128565.v1 | rca_mx23 | Right common anterior maximum near wall in millimeters |
| | phs000286.v5.pht001978.v1.phv00128566.v1 | rcp_mx23 | Right common posterior maximum near wall in millimeters |
| **MESA** | | | |
| | phs000209.v13.pht001116.v10.phv00084877.v2 | lcfwmax1 | LEFT COMMON CAROTID FAR WALL MAX (mm) |
| | phs000209.v13.pht001116.v10.phv00084881.v2 | lcnwmax1 | LEFT COMMON CAROTID NEAR WALL MAX (mm) |
| | phs000209.v13.pht001116.v10.phv00084956.v2 | rcfwmax1 | RIGHT COMMON CAROTID FAR WALL MAX (mm) |
| | phs000209.v13.pht001116.v10.phv00084959.v2 | rcnwmax1 | RIGHT COMMON CAROTID NEAR WALL MAX (mm) |
| | phs000209.v13.pht001121.v3.phv00087557.v1 | rcfwmaxf | RIGHT COMMON CAROTID FAR WALL MAX (mm) |
| | phs000209.v13.pht001121.v3.phv00087558.v1 | rcnwmaxf | RIGHT COMMON CAROTID NEAR WALL MAX (mm) |
| | phs000209.v13.pht001121.v3.phv00087559.v1 | lcfwmaxf | LEFT COMMON CAROTID FAR WALL MAX (mm) |
| | phs000209.v13.pht001121.v3.phv00087560.v1 | lcnwmaxf | LEFT COMMON CAROTID NEAR WALL MAX (mm) |
| | phs000209.v13.pht001528.v1.phv00111971.v1 | rcfwmax4 | RIGHT COMMON CAROTID FAR WALL MAX |
| | phs000209.v13.pht001528.v1.phv00111975.v1 | rcnwmax4 | RIGHT COMMON CAROTID NEAR WALL MAX |
| | phs000209.v13.pht001528.v1.phv00112047.v1 | lcfwmax4 | LEFT COMMON CAROTID FAR WALL MAX |
| | phs000209.v13.pht001528.v1.phv00112051.v1 | lcnwmax4 | LEFT COMMON CAROTID NEAR WALL MAX |

#### S3.3 Step 3: Perform QC on candidate variables

A primary goal of the QC step is to verify that study variables selected for harmonization are consistent with the study-specified metadata and do not contain impossible values. We implement a number of general checks as well as checks that are specific to each harmonized variable. Many of these checks require specific knowledge about properties of and measurement techniques for the phenotype being harmonized. DCC analysts acquire this knowledge by reading published and online descriptions of relevant techniques, consulting study protocols, and consulting with WG experts.

The general checks of candidate variables for each study include the following:

1. Are there a large number of missing values?
   a. If yes, can the missingness be explained by other factors, such as a questionnaire skip pattern?
   b. Were all missing codes recorded by the study, or are there values in the data that could represent unrecorded missing codes (e.g., “9” or “99”)?
2. Does the distribution of values fit within the expected range for the phenotype being harmonized?
   a. Is this distribution affected by participant ascertainment for this study, such as a study that primarily recruited a specific population or specific disease cases?
   b. Were extreme values winsorized?
   c. Are there any impossible values, such as negative analyte concentrates or composition fractions over 100%?
   d. Are there any batch effects that could introduce heterogeneities?
3. Are the data values generally consistent with other related variables measured at the same time point?

Many of these steps require knowledge of how the phenotype to be harmonized relates to other phenotypes (e.g., SBP and DBP) and whether each study has measured the related variables at the same time point. These specific checks vary from phenotype to phenotype; we give more detail in the four examples below.

If QC issues are discovered in a candidate variable, DCC analysts decide if those differences can be corrected using other related variables in the study accession, if a different study variable can be used, or if the study should be excluded from harmonization. These decisions are made in consultation with the WG and with the study liaisons on a case-by-case basis for each phenotype and study. After QC-related decisions have been made, the selected candidate variables are referred to as "component variables" and are used in the next step to calculate the desired harmonized variable.

In the following subsections, we address any substantial QC issues identified for each of the four example harmonized variables. We also discuss how missing values were handled.

###### S3.3.1 Step 3 example: ever_smoker_baseline_1

The component phenotype variables for the ever_smoker_baseline_1 variable originated from questionnaires, where participants were asked about their smoking habits.

Participants who replied that they have never smoked were generally not asked more detailed questions about smoking habits, such as the number of cigarettes smoked per day or the age at which they started smoking. These skip patterns led to large numbers of expected missing values in the component variables selected for this phenotype in many studies. Participants who have missing responses for all questions about smoking history were given a missing value for ever_smoker_baseline_1. Participants who responded to a direct question about whether they ever smoked, but have missing values for indirect questions about smoking behavior were coded using the direct response only. Those who have a missing value for whether they ever smoked, but have responses to other questions that clearly indicate a smoking history (such as number of cigarettes per day) were assigned a value of 1 (current or former smoker status).

Variables were also assessed to identify responses where participants gave conflicting responses, such as responding that they have never smoked to one question but smoked two packs of cigarettes per day to another. When discrepant responses were identified, if any response indicated use of cigarettes then we assigned a value of 1 (current or former smoker status).

###### S3.3.2 Step 3 example: bp_systolic_1

This variable was harmonized by averaging multiple blood pressure measurements collected at a single clinic visit, so QC was performed on each individual measurement as well as the average. The QC process for bp_systolic_1 is further complicated because blood pressure measurements are taken as a pair of systolic and diastolic measurements, so they must be QC'd together. We first checked each paired SBP or DBP measurement for a given participant; both measurements in the pair were set to missing if either measurement was missing; if either measurement was negative; or if the DBP measurement was larger than the SBP measurement. The averages were then calculated using the remaining sets of paired SBP and DBP measurements. After the average SBP and DBP values were calculated, these checks were performed again on the average, and any failed results were set to missing for both bp_systolic_1 and the related harmonized variable bp_diastolic_1.

Some studies provided one or more sets of paired SBP/DBP measurements for each participant plus a study-calculated average of these measurements. In these cases, DCC analysts checked for discrepancies between DCC-calculated and study-calculated measurements. In some cases, such discrepancies were due only to the handling of missing and biologically impossible values as described above. No additional discrepancies were identified in the studies processed thus far, but would have been noted in the harmonization comments if they existed.

###### S3.3.3 Step 3 example: il6_1

The standard checks described above were performed for IL6. Additional QC was possible for this variable because some studies provided information about the batch or processing plate on which the IL6 assay was run for a set of participants. If available, this information was used to test for plate-associated batch effects by performing an F-test on IL6 values adjusted for age, sex, and (if applicable) subcohort. If the F statistic was significant (p-value < 0.05), a Wilcoxon rank sum test was performed to investigate the robustness of the apparent effect. For this phenotype variable, no study had significant F-test p-values for plate (Web Figure S1).

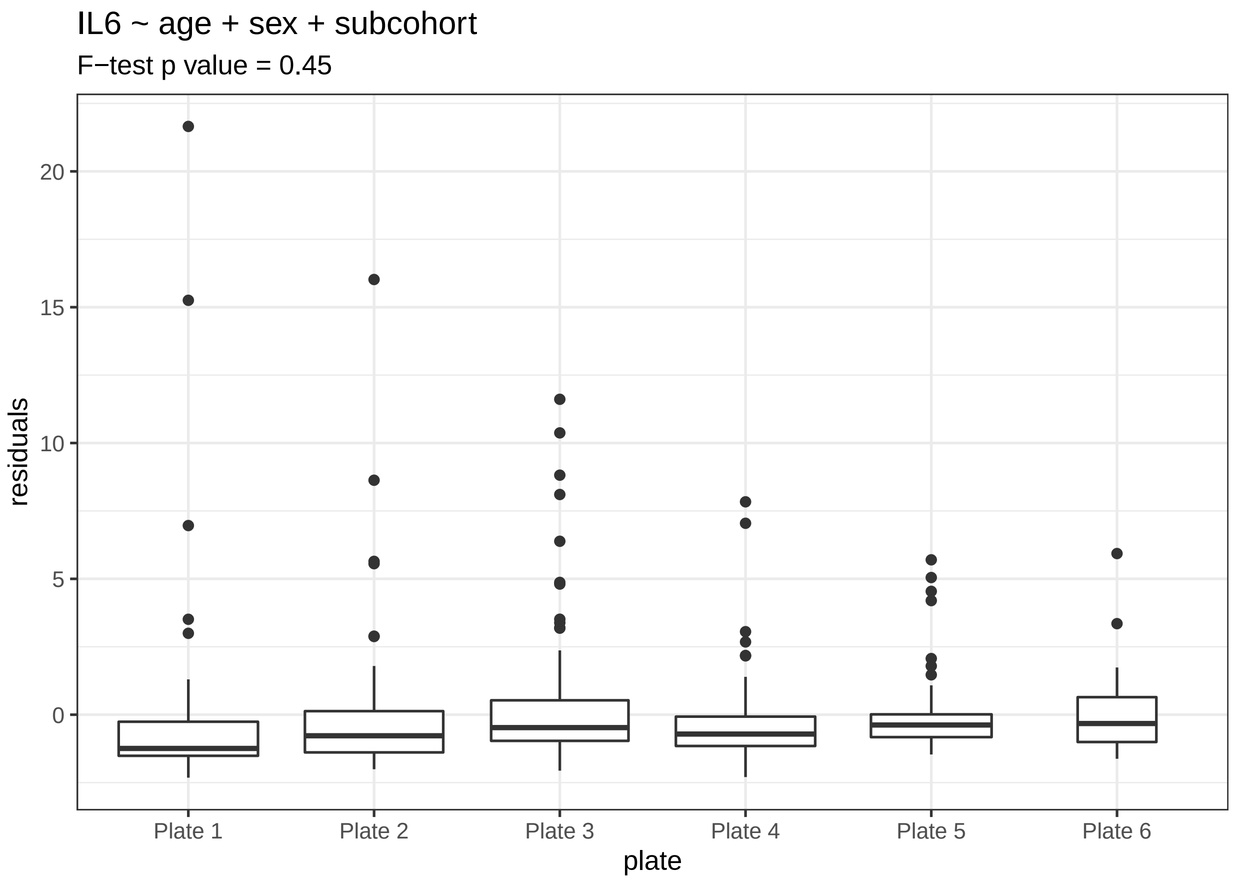

Figure S1: Distribution of IL6 by assay plate in the after adjustment for age, sex, and subcohort for FHS.

###### S3.3.4 Step 3 example: cimt_2

Candidate variables were inspected for missingness. In cases where the individual measures of cIMT were missing, cimt_2 was calculated using any available non-missing values. The range of cIMT was calculated for each study to identify biologically implausible values (i.e. positive numbers and not substantially greater than the usual width of a carotid artery, 6-7 mm), but none were found. For the CHS study, which provided original and reread measurements for some participants, we checked for consistency between both sets of measurements.

##### S3.4 Step 4: Construct harmonization algorithms

When processing study variables, it is often necessary to work with data from a specific subset of participants within a study due to that study's data collection or organization in dbGaP phenotype files. We define the set of component variables and the algorithm used for a single study or subset of participants within that study as a "harmonization unit." Participants are grouped into the same harmonization unit if their data can be treated similarly. Practically, this often translates to participants whose data are available in the same dbGaP datasets. In some cases, the same variables and algorithm to transform those variables can be used for participants in different datasets within a study. More complex studies often require multiple harmonization units, depending on study design, data organization, and heterogeneity within the study. For example, FHS comprises multiple subcohorts (e.g., Original Cohort, Offspring Cohort, Third Generation Cohort, etc.); for some phenotypes, data for multiple subcohorts are available in a single phenotype dataset (e.g., pht002891 containing IL6 measurements for the Offspring, New Offspring Spouse, and Omni 1 subcohorts), while other datasets contain only variables for a single subcohort (e.g., pht000161 containing IL6 data for the Offspring cohort only). Due to this varying structure, the set of subcohorts that can be included in a single harmonization unit is often different for different harmonized phenotype variables, even for the same study. When using harmonized variables as components for a new harmonized variable, we use a single multi-study harmonization unit because the component variables are already comparable across studies.

Once the harmonization units have been established for the harmonized variable, we implement the harmonization algorithm for each harmonization unit as an R function. This R function accepts the component variable(s) in a specific format as an argument to the function, processes them for harmonization, and returns a data frame with columns “topmed_subject_id” (the unique participant identifier), the name of the phenotype being harmonized (without the post-appended concept variant number, which is assigned automatically when the final harmonized variable is added to the database), and “age”. For demographic phenotypes with no associated age, the “age” column is not included. The function handles any missing or incorrect values and harmonizes the component variables to fit the harmonized variable definition. We include comments for each step to give a general explanation of how the data are being processed. Missing values were generally not imputed, unless otherwise described in the harmonization comments for a harmonized variable. Examples of one harmonization algorithm for each of the four example traits is shown in the sections below. The harmonization functions are included in the JavaScript Object Notation (JSON) documentation provided in our GitHub repository (https://github.com/UW-GAC/topmed-dcc-harmonized-phenotypes).

The data are provided to the harmonization function as an R list with a specified format. If all component variables are from dbGaP study accessions (i.e. not DCC-harmonized), this list has one top-level element named “source_data”. The “source_data” element is also a list containing dbGaP study data (e.g, Web Box S1. The elements are named by the dbGaP accession for each dataset (e.g., pht012345), and each of those elements is a data frame in which the columns are “topmed_subject_id” and the selected component variable names in dbGaP from that dataset. If the component variables are DCC-harmonized variables (e.g., when using harmonized height and weight to calculate BMI), the list has a different top-level element named “harmonized_data”. The “harmonized_data” element is also a list containing one element for each component harmonized variable (e.g., Web Box S2. Each of those elements is a data frame whose columns are “topmed_subject_id”, the component_harmonized variable name, and the age at measurement of that variable.

Because each harmonization function produces a single harmonized variable, along with paired age at measurement, there is no need to retain a participant record with a missing value and such records are removed from harmonized data frame by the harmonization function.

##### S3.4.1 Step 4 example: ever_smoker_baseline_1

The harmonization function in Web Box S3 was used for participants from the Cleveland Family Study. In this case, age at measurement is stored in the same dbGaP dataset as the component variables. The function first subsets the data to the appropriate visit for each participant. It then converts variables from character type (as they are stored in the DCC database) to numeric type and recodes them as necessary for calculating the ever_smoker_baseline_1 variable. Once the harmonized variable is calculated, the function returns a data frame with no missing values and columns “topmed_subject_id”, “ever_smoker_baseline”, and “age”. Note that the concept variant number of this variable (“1”) is not part of the “ever_smoker_baseline” column name, since it is assigned automatically by the code when the final harmonized variable is added to the database. Second, the column representing age at measurement is called “age” in this function but is renamed to “age_at_ever_smoker_1” in results distributed to users.

The full set of harmonization functions for all harmonization units for the ever_smoker_baseline_1 variable

Box S1: An example of the `phenlist` R data structure with simulated data for three component study variables in from dbGaP datasets. Because "height" and "weight" appear in the same dbGaP dataset, they are in the `phtXXXXXX1` element together.

```
phen_list$source_data
phen_list$source_data$phtXXXXX1
topmed_subject_id height  weight
                1     69     160
                2     68     138
                3     59     185
                4     69     155
                5     71     152
phen_list$source_data$phtXXXXX2
topmed_subject_id age
                1  71
                2  43
                3  23
                4  52
                5  25
```

Box S2: An example of the `phenlist` R data structure with simulated data for two component harmonized variables.

```
phen_list$harmonized_data
phen_list$harmonized_data$height_1
  topmed_subject_id height_1 age_at_height_1
1                 1      177              45
2                 2      180              37
3                 3      165              22
4                 4      188              59
5                 5      170              41
phen_list$harmonized_data
phen_list$harmonized_data$weight_1
  topmed_subject_id weight_1 age_at_weight_1
1                 1       81              45
2                 2       95              37
3                 3       63              22
4                 4      102              59
5                 5       76              41
```

are given in the publicly-available documentation.

Box S3: The harmonization function used to harmonize ever smoker status for CFS.

```
harmonize <- function(phen_list) {
  library(dplyr)

  df <- phen_list$source_data$pht001902 %>%

    # Subset to baseline visit. Some respondents baseline is visit 5
    filter(visit %in% c("1", "5")) %>%
    group_by(topmed_subject_id) %>%
    arrange(topmed_subject_id, visit) %>%
    filter(row_number(topmed_subject_id) == 1) %>%
    ungroup() %>%

    # Convert variables to numeric
    mutate_if(is.character, as.numeric) %>%

    # Recode encoded values and NA as 0
    mutate(AGESMOK = ifelse(AGESMOK %in% c(-1, -2, NA), 0, AGESMOK),
        AVGSMOK = ifelse(AVGSMOK %in% c(-1, -2, NA), 0, AVGSMOK),
        MONSMOKE = ifelse(MONSMOKE %in% c(-1, NA), 0, MONSMOKE),
        NOWSMOKE = ifelse(NOWSMOKE %in% c(-1, NA), 0, NOWSMOKE),
        # code ever_smoker_baseline as 1 if any smoking variables are positive
        ever_smoker_baseline = as.numeric(as.logical(
            SMOKED + AGESMOK + AVGSMOK + MONSMOKE + NOWSMOKE
        ))) %>%

    # Select only ID, age and phenotype
    select(topmed_subject_id, age, ever_smoker_baseline) %>%

    # Exclude incomplete records
    na.omit() %>%
    return()
}
```

##### S3.4.2 Step 4 example: bp_systolic_1

The harmonization function shown in Web Box S4 was used for participants in the Jackson Heart Study. This function works with variables from two different dbGaP datasets, one that includes the systolic, diastolic, and zero reading blood pressure component variables (pht001974) and one that has information about participant age (pht001949). The function sets an encoded character-type "NA" value in the study variables to missing and converts the data type of the blood pressure measurements to numeric for future processing. It then corrects each blood pressure measurement for the random-zero instrument by subtracting the zero readings to the measured values. The next step is to perform the QC step of setting both SBP and DBP measurements in a pair to missing when the SBP measurement is less than the DBP measurement or when the value for one measurement in the pair is missing. Finally, the function calculates the average SBP value using the paired readings and returns the harmonized data values in a data frame with columns "topmed_subject_id", "bp_systolic", and "age".

Box S4: The harmonization function used to harmonize SBP for JHS.

```
harmonize <- function(phen_list){

  # Get dataset.
  dataset <- inner_join(phen_list$source_data$pht001949,
                        phen_list$source_data$pht001974,
                        by = "topmed_subject_id")

  # Substitute the value of 'NA' to missing.
  dataset$SBPA13[dataset$SBPA13 %in% 'NA'] <- NA
  dataset$SBPA14[dataset$SBPA14 %in% 'NA'] <- NA
  dataset$SBPA15[dataset$SBPA15 %in% 'NA'] <- NA
  dataset$SBPA16[dataset$SBPA16 %in% 'NA'] <- NA
  dataset$SBPA17[dataset$SBPA17 %in% 'NA'] <- NA
  dataset$SBPA18[dataset$SBPA18 %in% 'NA'] <- NA

  # Convert character values to numeric.
  dataset <- mutate_if(dataset, is.character, as.numeric)

  # Calculate random-zero corrected BP readings.
  dataset <- mutate(dataset,
                    sbp1 = SBPA13 - SBPA15,
                    dbp1 = SBPA14 - SBPA15,
                    sbp2 = SBPA16 - SBPA18,
                    dbp2 = SBPA17 - SBPA18)

  # Set systolic BP to NA when systolic BP is less than diastolic BP from the same reading
  # or when diastolic BP from the same reading is NA.
  dataset <- mutate(dataset,
                    sbp1 = ifelse(sbp1 >= dbp1, sbp1, NA),
                    sbp2 = ifelse(sbp2 >= dbp2, sbp2, NA))

  # Calculate the average systolic BP.
  dataset$bp_systolic <- rowMeans(dataset[, c("sbp1", "sbp2")], na.rm = TRUE)

  # Rename and select the output variables.
  dataset <- rename(dataset, age = AGE01) %>%
             select(topmed_subject_id, bp_systolic, age)

  # Remove records with NAs from dataset.
  dataset <- na.omit(dataset)

  return(dataset)
}
```

##### S3.4.3 Step 4 example: il6_1

The function shown in Web Box S5 was used to harmonize data from participants in the "Coronary Artery Risk Development in Young Adults" (CARDIA) study. This function uses variables from two datasets, one with IL-6 measurements (pht001862) and the other with age information for that time point (pht001851). The function sets the encoded character-type "NA" value to missing. It then handles measurements outside the upper LOD for this assay by setting them to the upper LOD, for consistency across studies. Because no values below the lower LOD were observed for participants in this harmonization unit, correction for the lower LOD was not necessary in this harmonization function. Finally, it selects the appropriate set of data frame columns ("topmed_subject_id", "il6", and "age"); converts them to the proper data type (numeric); removes missing records; and returns the data frame with harmonized values.

Note that CARDIA did not provide an indicator of whether an assay failed for each participant. For studies that did provide this variable, the harmonization function included an additional step to remove participants with failed assays from the harmonized variable.

Box S5: The harmonization function used to harmonize IL-6 for CARDIA.

```
harmonize <- function(phen_list){
  library(dplyr)

  # Get dataset and rename variables.
  dataset <- inner_join(phen_list$source_data$pht001862,
                        phen_list$source_data$pht001851,
                        by = "topmed_subject_id") %>%
             rename(age = EX6_AGE, il6 = FL6IL6)

  # Substitute the value of 'NA' to missing.
  dataset$age[dataset$age %in% 'NA'] <- NA
  dataset$il6[dataset$il6 %in% 'NA'] <- NA

  # Set IL6 values above the upper limit of detection to the upper limit of detection.
  dataset$il6[dataset$FL6IL6CM == 'High > 12'] <- 12

  # Select the output variables.
  dataset <- select(dataset, topmed_subject_id, il6, age)

  # Convert character values to numeric.
  dataset <- mutate_if(dataset, is.character, as.numeric)

  # Remove records with NAs from dataset.
  dataset <- na.omit(dataset)

  return(dataset)
}
```

##### S3.4.4 Step 4 example: cimt_2

The harmonization function shown in Web Box S6 was used to harmonize cimt_2 for two subcohorts from the Multi-ethnic Study of Atherosclerosis (MESA) study. The data for these subcohorts are stored in different datasets, but the structure and organization of these datasets are similar enough that the variables can be harmonized together. For other phenotypes, this is generally not the case, as subcohorts within a study are typically processed in two different harmonization units.

This function first renames variables from the two datasets so that they can be combined into one data frame. It then converts the data values to numeric types so that the cimt_2 values can be calculated as a mathematical average of the four measurements of maximum carotid intima media thickness. The appropriate columns are selected ("topmed_subject_id", "cimt", and "age") before removing missing records and returning the final data frame.

Box S6: The harmonization function used to harmonize cimt_2 for participants in the MESA Classic and MESA Family subcohorts.

```
harmonize <- function(phen_list){
  library(dplyr)
  source_data <- phen_list$source_data

  # Rename variables in Family Exam dataset to match Classic.
  source_data$pht001121 <- rename(source_data$pht001121, age1c = agefc,
                                  rcfwmax1 = rcfwmaxf, rcnwmax1 = rcnwmaxf,
                                  lcfwmax1 = lcfwmaxf, lcnwmax1 = lcnwmaxf)

  # Bind dataframe row-wise.
  harmonized <- bind_rows(source_data) %>%
    # Convert character vectors to numeric.
    mutate_if(is.character, as.numeric) %>%
    # Specify calculations will be row-wise.
    rowwise() %>%
    # Select and rename necessary variables, calculate mean cimt.
    transmute(topmed_subject_id, age = age1c,
              cimt = mean(c(lcfwmax1, lcnwmax1, rcfwmax1, rcnwmax1), na.rm = TRUE)) %>%
    # Exclude rows with missing data.
    na.omit()

  return(harmonized)
}
```

#### S3.5 Step 5: Produce and QC multi-study harmonized phenotype

Once component traits and harmonization algorithms are completed for all harmonization units, we combine the harmonized values, perform QC, and write the harmonized variable to the database. The first step in this process is to generate a configuration file that contains metadata about the harmonized variable, as well as the component variables and algorithm for each harmonization unit. We then run a series of python and R scripts that accept this configuration file as an input, process the information, and produce an interim harmonized variable for further QC. If the QC process reveals issues with the harmonized variable, we either revise the harmonization algorithm, choose new component variables, or exclude the study from harmonization. Once any QC issues have been resolved, we use the internal scripts to add the finalized harmonized variable to the database.

The configuration file required by the scripts is an Extensible Markup Language (XML) file that includes all information necessary to produce the harmonized variable. The configuration file for one harmonized variable contains three child nodes:

1. A "metadata" node, which specifies information such as the variable name, description, data type, and any encoded values. It also includes a path to a file containing the harmonization comments (described below) and a term from the Unified Medical Language System (UMLS) metathesaurus (6) that best fits

the phenotype being harmonized, which allows future investigators to more easily identify the meaning of the harmonized variable.
2. The "input" node, which contains information about each harmonization unit comprising the harmonized variable. This node has child nodes for each harmonization unit, which specify the internal database identifiers for each component variable used and the path to a file containing the definition of the harmonization function for that unit.
3. The "output" node specifies where the interim output files should be written on disk.

An example configuration file is shown in Web Box S7. For brevity, we have removed all but two "input_unit" nodes.

Next, we run internal R scripts to produce the harmonized variable using the configuration file as input. These scripts retrieve the component variables from the database, run the harmonization algorithms for each unit, and combine the harmonized data values from each unit into one data frame. The following set of automated checks are run during this process:

1. No component variables are from outdated study versions.
2. All component variables for a harmonization unit come from the same study accession.
3. Required metadata exists in the configuration file.
4. Component traits are not from outdated study versions or different studies.
5. The data type specified in the metadata is consistent with the data type of the harmonized values.
6. There is only one record per participant.
7. The order of the input component variables and data values does not change the output harmonized phenotype values.

The initial runs of the script produce data files containing harmonized values for each participant, which are used to perform additional, interactive QC.

The general QC process involves checking for differences in the distribution of the harmonized values by study, harmonization unit, and study-subcohort. Analysts also fit a linear model that adjusts for age, sex, and ancestry group, and re-check distributions of residuals from this model. If applicable, analysts also inspect the harmonized values by additional grouping variables that could affect the phenotype, such as medication use. Specific QC steps performed for the four example variables are described in the examples below.

Once any adjustments to the harmonization units are made and any QC issues have been resolved, DCC analysts write a free-text summary of the harmonization in Markdown format with important notes for users of the data. These notes include a more detailed description of the phenotype definition than can be given in the metadata description, plus any general or study-specific issues that were encountered. An example for ever_smoker_baseline_1 is shown in Web Box S8.

The last step in harmonization is to run the harmonization scripts with a flag that adds the harmonized variable to the database. At this point, the concept variant number and database identifiers are assigned automatically and harmonization for this variable is considered complete. The "age" variable is also renamed to the harmonized variable name prepended with "age_at_" (e.g., age_at_ever_smoker_baseline_1).

###### S3.5.1 Step 5 example: ever_smoker_baseline_1

Specific QC checks for the ever_smoker_baseline_1 variable include inspecting the frequency of smokers by harmonization unit and subcohort (described in the main text). We also verified the consistency with a related harmonized variable indicating current smoking status (current_smoker_baseline_1) to ensure that all participants who are current smokers also were labeled as ever smokers.

The harmonization comments for this phenotype are shown in Web Box S8.

Box S7: An example configuration file for the ever_smoker_baseline_1 harmonized variable. Only two harmonization units are shown for brevity.

```
<config>
  <metadata>
    <target>
      <name>ever_smoker_baseline</name>
      <description>Indicates whether subject ever regularly smoked cigarettes.</description>
      <data_type>encoded</data_type>
      <encoded_values>
        <value code="0">Never a cigarette smoker</value>
        <value code="1">Current or former cigarette smoker</value>
      </encoded_values>
      <ontology>
        <record>
          <source>UMLS</source>
          <version>2018AB</version>
          <code>C1519384</code>
          <relationship>Comparable</relationship>
        </record>
      </ontology>
    </target>
    <update>
      <harmonized_trait_set_id>21</harmonized_trait_set_id>
    </update>
    <qc_document>analyst_comments.md</qc_document>
  </metadata>
  <input>
    <input_unit unit_id="ARIC">
      <source_trait_id>376622</source_trait_id>
      <source_trait_id>376623</source_trait_id>
      <source_trait_id>376624</source_trait_id>
      <source_trait_id>376629</source_trait_id>
      <source_trait_id>376630</source_trait_id>
      <age_trait_id>373913</age_trait_id>
      <custom_function>function_def_ARIC.R</custom_function>
    </input_unit>
    <input_unit unit_id="CARDIA">
      <source_trait_id>279458</source_trait_id>
      <age_trait_id>278698</age_trait_id>
      <custom_function>function_def_CARDIA.R</custom_function>
    </input_unit>
  </input>
  <output>
    <output_directory>output</output_directory>
    <output_prefix>output</output_prefix>
  </output>
</config>
```

Box S8: Abbreviated harmonization comments for the ever_smoker_baseline_1 harmonized variable.

```
When available, we used component variables from smoking history questionnaires
to harmonize this trait, rather than derived variables, to promote
reproducibility and for handling inconsistencies.  In the case of contradictory
information, as a general approach, any positive indication that a subject
smoked regularly will cause them to be coded as an "ever smoker" (e.g. they
respond that they have never smoked, but _smoked a positive number of cigarettes
per day_ when they did smoke).

#### HVH

There are multiple observations for many subjects in the HVH phenotype file. In
these instances, we used the earliest observation for harmonization. Although
this harmonized phenotype is designated as "baseline", the concept of "baseline"
does not apply to HVH based on its study design. Consult the study documentation
for more details (phs001013).
```

##### S3.5.2 Step 5 example: bp_systolic_1

Specific QC checks for the bp_systolic_1 variable include inspection of density plots by age, ancestry group, sex, study, and antihypertensive medication status. We also fit a linear model that adjusted the bp_systolic_1 values for age, sex, and ancestry group, and checked the residual distributions by the same groups. No notable differences between studies were present after adjustment, so all of the initial variables and studies identified to be included in harmonization were kept after the QC checks (Web Figure S2).

Because SBP and DBP were harmonized as separate variables but with related QC, we compared bp_systolic_1 values with its paired harmonized variable for DBP, bp_diastolic_1, to confirm that no participants had SBP values that were smaller than the bp_diastolic_1 values. While this handling was implemented in the algorithms for each variable separately, we verified that it had been correctly applied after harmonization of both variables.

The harmonization comments for bp_systolic_1 are shown in Web Box S9. In particular, the harmonization algorithms for some units differed from the original definition due to data availability. These harmonization units were retained in the final dataset, and the differences were noted in the harmonization comments.

##### S3.5.3 Step 5 example: il6_1

The QC process for the harmonized variable il6_1 is discussed in the main text..

The harmonization comments in Web Box S10 include general information about how measurements outside an assay's LOD were handled. The comments also include one table detailing which exam was used for each included study-subcohort; a second table specifying information about the assay used for each study-subcohort; and a third table indicating the specimen type on which the assay was run, if known.

##### S3.5.4 Step 5 example: cimt_2

QC checks for cimt_2 included inspection of the distribution of values by study and harmonization unit as well as the residuals after adjusting for age, sex, and ancestry group (Web Figure S3). Before adjustment, the phenotype values for cimt_2 look notably different for some studies, but they are much more similar after adjustment for these factors. Even after adjustment, one study (study D) had lower values on average compared to other studies. We consulted with the study and were not able to find an explanation for this

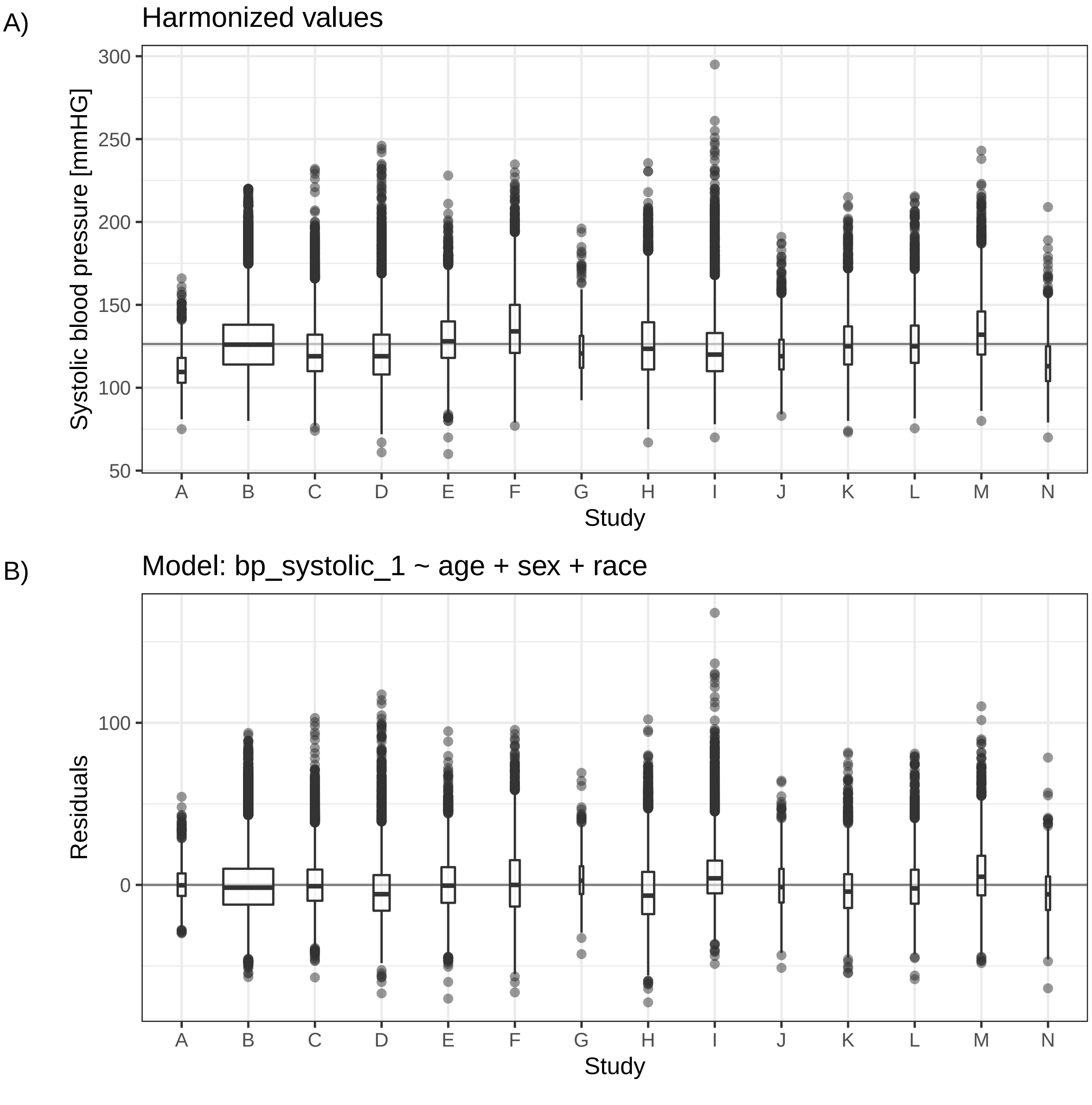

Figure S2: A) Distribution of bp_systolic_1 by study. The mean value across all studies is shown as the solid horizontal line. B) Distribution of bp_systolic_1 by study after adjusting for age, sex, and race. The solid horizontal line shows the y = 0 line, around which these residuals should be centered.

Box S9: Abbreviated harmonization comments for the bp_systolic_1 harmonized variable.

This variable was harmonized by taking the average of two systolic blood pressure (BP) measurements collected at a single clinic visit. When more than two measurements were collected, the average was calculated using the second and third measurements. In cases where either of the measurements was missing, the average was calculated discarding the missing value. If a study used a random-zero sphygmomanometer and the variables representing the zero readings were available in dbGaP, the zero reading adjustments were applied in the harmonization. In cases where the individual BP measurements were not available in dbGaP, a mean systolic BP variable derived by the study was used for harmonization. For paired systolic and diastolic BP measurements, if one of the paired measurements was missing or the systolic BP was less than the diastolic BP, the values for both systolic BP and diastolic BP for that pair were set to missing. This harmonized variable was not adjusted for antihypertensive medication status.

###### COPDGene

Only one blood pressure measurement was available for each subject at baseline, so an average systolic BP value could not be calculated. The single measurement was used for harmonization of systolic BP.

###### FHS

Because antihypertensive medication was not recorded before Exam 4 for the Original cohort, systolic BP values from Exam 4 were used for harmonization.

###### GOLDN

Only one blood pressure measurement was available for each subject at baseline, so an average systolic BP value could not be calculated. The single measurement was used for harmonization of systolic BP.

###### Instrumentation

The instruments used for BP measurements were different among studies, including standard manual sphygmomanometers, random-zero sphygmomanometers, and automated digital blood pressure monitors.

Box S10: Abbreviated harmonization comments for the il6_1 harmonized variable.

```
This variable was harmonized by converting the component study variables to the
appropriate unit of measure as needed and, when possible, accounting for
measurements outside an assay's limits of detection (LOD). If the information
was available, measurements below the lower limit of detection (LLOD) were set
to the LLOD and measurements above the upper limit of detection (ULOD) were set
to the ULOD unless otherwise indicated in the study-specific sections below.
Some studies identified subjects with measurements outside the LOD; see table
below for more details. The assay(s) used to measure IL6 concentration from
serum or plasma differed by study and/or subcohort.

#### Exam visit for IL6 measurements

| Study or subcohort | Visit |
|---------|----------|
| CARDIA | Year 15/Exam 6 |
| CFS | Visit 5 |
| CHS_Original | Baseline visit |
| CHS_AfricanAmerican | Baseline visit |
| FHS_Offspring | Exam 7 |
| FHS_NewOffspring Spouse | Exam 1 |
| FHS_Gen3 | Exam 1 |
| FHS_Omni1 | Exam 3 |
| MESA_Classic | Exam 1 Main |

#### Assay and limits of detection for IL6 measurements

| Study or subcohort | Assay | LLOD | ULOD | Differentiated^1^ |
|---------|----------|---------|---------|---------|
| CARIDA | ELISA | 0.10 pg/mL | 12 pg/mL | Yes |
| CFS | ELISA | 0.08 pg/mL | 15 pg/mL | Yes |
| CHS | ELISA  | < 0.7 pg/mL | 300 pg/mL | No |
| FHS_Offspring | ELISA | < 0.7 pg/mL | 300 pg/mL | No |
| FHS_Gen3 | ELISA | 0.039 pg/mL | NA  | No |
| FHS_NewOffspringSpouse | ELISA |  0.15 pg/mL | NA  | No |
| FHS_Omni1 | ELISA | 0.15 pg/mL | NA  | No |
| MESA_Classic | ELISA  | 0.09 pg/mL | 13.0 pg/mL | Yes |

1.  The study included information indicating which measurements were below or
above the limit of detection. If "Yes", measurements outside the LOD can be
identified using component study or subcohort variables.

#### Specimen type for IL6 measurements

Table includes studies or subcohorts with known specimen types only.

| Study or subcohort | Specimen |
|---------|----------|
| CHS | Serum |
| FHS | Serum |
```

difference. Given that the difference between this study and others was small, we retained this study in the final harmonized variables, which allows investigators either to use or to remove this sample set.

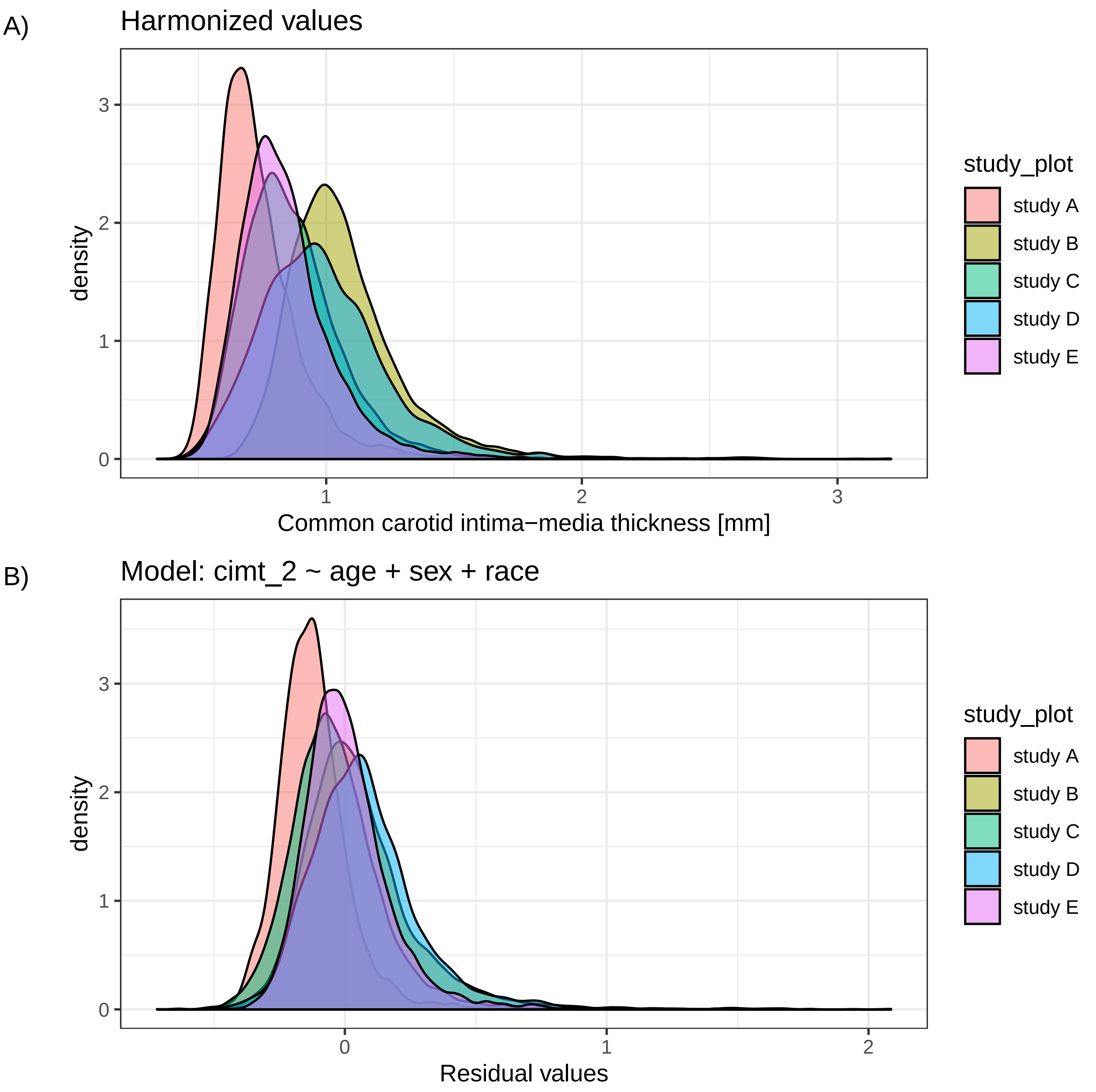

Figure S3: A) Distribution of cimt_2 values by study. B) Distribution of cimt_2 values by study after adjustment for age, sex, and race.

The harmonization comments for cimt_2 are shown in Web Box S11. For this variable, we report small differences from the original harmonization plan so that users of the data can choose whether or not to use those studies' data values. We also provide the instrument used to measure cIMT values for each study, which allows users to select which studies to include in analysis or to adjust for different instruments if desired.

Box S11: Abbreviated harmonization comments for the cimt_2 harmonized variable.

```
This variable was harmonized by taking the mean of the following four
measurements of common carotid intima-media thickness (IMT): maximum left near
wall IMT, maximum left far wall IMT, maximum right near wall IMT and maximum
right far wall IMT. In cases where values for individual measures of IMT were
missing, mean IMT was calculated ignoring the missing values. Where possible,
this variable was derived with component measures of IMT, but in cases where the
components were not available in dbGaP, mean-of-max IMT variables derived by the
studies were used for harmonization.

#### CHS

Baseline carotid ultrasound scans for the Original cohort were reread due to
reader drift. Reread measurements of *_CHS_* subjects were used for
harmonization.

#### FHS

Measurements of *_FHS_* subjects were taken in systole and diastole.
Measurements in diastole were used for harmonization.

#### Instrumentation

Studies used different instruments at their carotid ultrasound exams:

| Study | Instrument                 |
|-------|----------------------------|
| ARIC  | Biosound 2000 II SA        |
| CHS   | Toshiba SSA-270A           |
| FHS   | Toshiba SSH-140A           |
| JHS   | Hewlett Packard SONOS 4500 |
| MESA  | GE Logiq 700               |
```

#### S4 Updating harmonized variables

When updating a previously-harmonized variable, analysts create a configuration file for the harmonized variable being updated using the information stored in the database, and then modify it to incorporate updates. To add phenotype values from new studies, harmonization units for those studies are constructed and added to the configuration file. Updates to the harmonized variable for a previously included study require updating the component variables to their most recent versions, which can be done automatically using the existing versioning in the dbGaP study and variable accession numbers (e.g., phs000007.v29 vs. phs000007.v30 for the FHS study accession). Even though the same QC processes applied to the original harmonized variable are also applied to the updated version, the updating process is generally much faster than producing a new harmonized phenotype. The DCC generally updates all harmonized variables in a dataset at the same time.

When a variable is updated, the updated variable has the same name and concept variant number , but a new record with an incremented version number for the updated variable is added to the database.

#### S5 Distributing harmonization results to the scientific community

After a group of related harmonized variables have been added to the database, we produce a dataset containing those variables for distribution to the scientific community. This process first consists of creating a record for the dataset (e.g., "Lipids") and the version of that dataset (e.g., v1) to be released. For updated versions, only a record for the new version of the dataset with an incremented version number is created (e.g., v2). Next, DCC staff link that dataset to the included harmonized variable versions. Once these records have been entered into the database, the dataset is created by running a function in the internal R package that creates all files for that dataset version using information stored in the database for distribution to National Institutes of Health (NIH) data repositories.

The eight datasets listed in Main text Table 2 have been submitted to two NIH data repositories, dbGaP (https://www.ncbi.nlm.nih.gov/gap/) and BioData Catalyst (https://biodatacatalyst.nhlbi.nih.gov/). Each dataset contains multiple harmonized phenotype variables and consists of (1) a data file and (2) documentation about the harmonization process. In the data file, we provide harmonized data values, age at measurement, and the harmonization unit used for each combination of participant and harmonized variable in the dataset. We also provide the dbGaP study accession and version that were used to harmonize each participant's data for a given harmonized phenotype variable, which allows the harmonized data to be linked to their consent value in that accession. For documentation, we provide a data dictionary with definitions and data types for each harmonized variable as well as Portable Document Format (PDF) documentation containing the harmonization comments for each variable, plus the list of component variables and the harmonization function for each contributing harmonization unit.

Histograms of the distributions of the harmonized variables presented in this paper are shown in Web Figure S4.

#### S6 Harmonized phenotype documentation and reproducibility

We provide full documentation for all harmonized phenotype variables in a GitHub repository (https://github.com/UW-GAC/topmed-dcc-harmonized-phenotypes). The repository contains one JSON documentation file for each harmonized phenotype variable, which includes the following information:

1. harmonized phenotype variable metadata such as name, description, measurement units, etc.;
2. the version number of the harmonized variable;
3. any controlled vocabulary terms attached to this harmonized phenotype variable;
4. the harmonization comments;

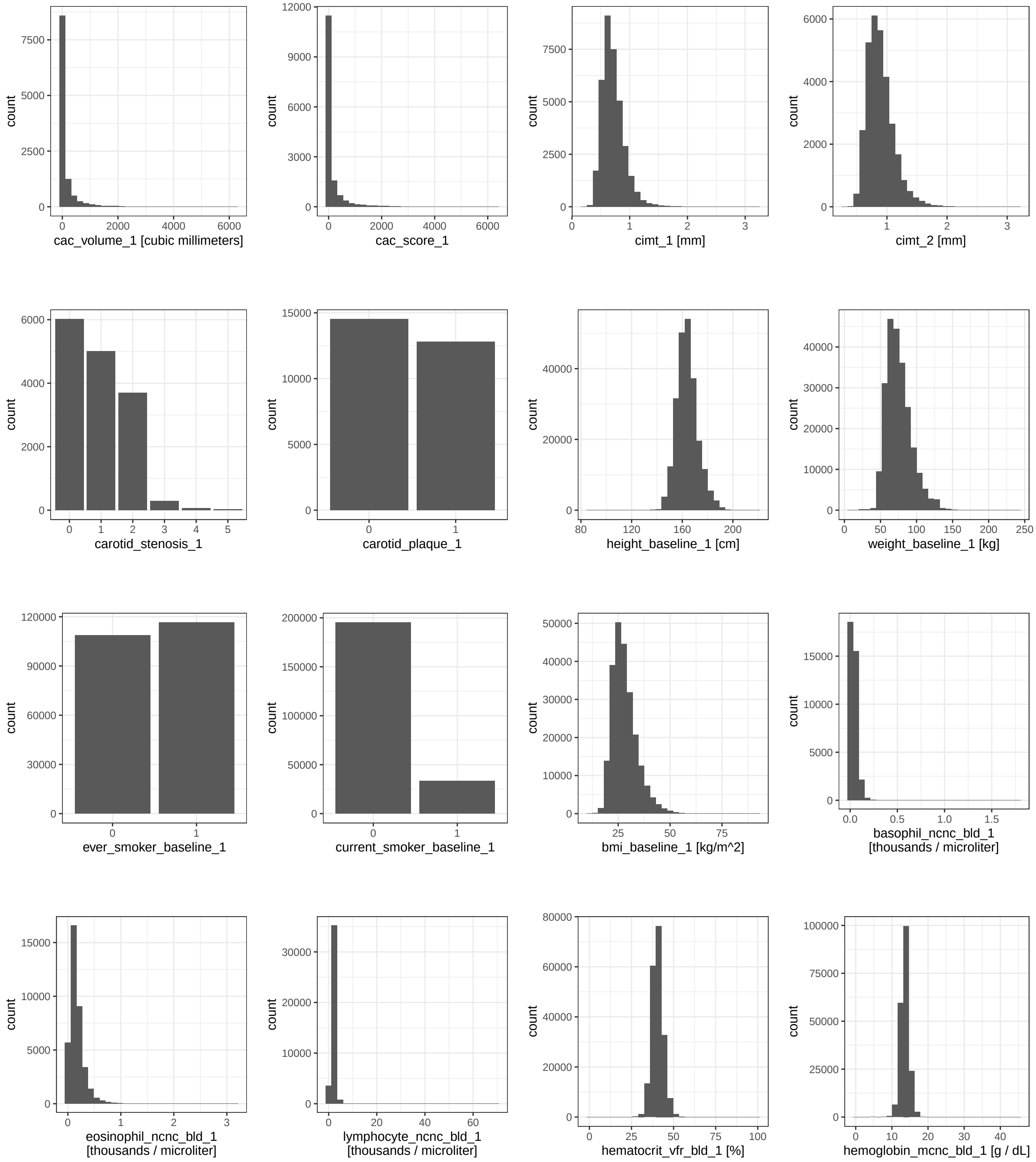

Figure S4: Histograms of the harmonized variables presented in this paper. For categorical variables, the ratio of some categories can be different than expected from the general population due to study size and recruitment strategy. For example, the sex ratio shown in the histogram for annotated_sex_1 indicates an excess of females, which is mainly due to the inclusion of the large, all-female WHI study. Due to the large number of categorical values in geographic_site_1 and subcohort_1, histograms for these variables are not shown.

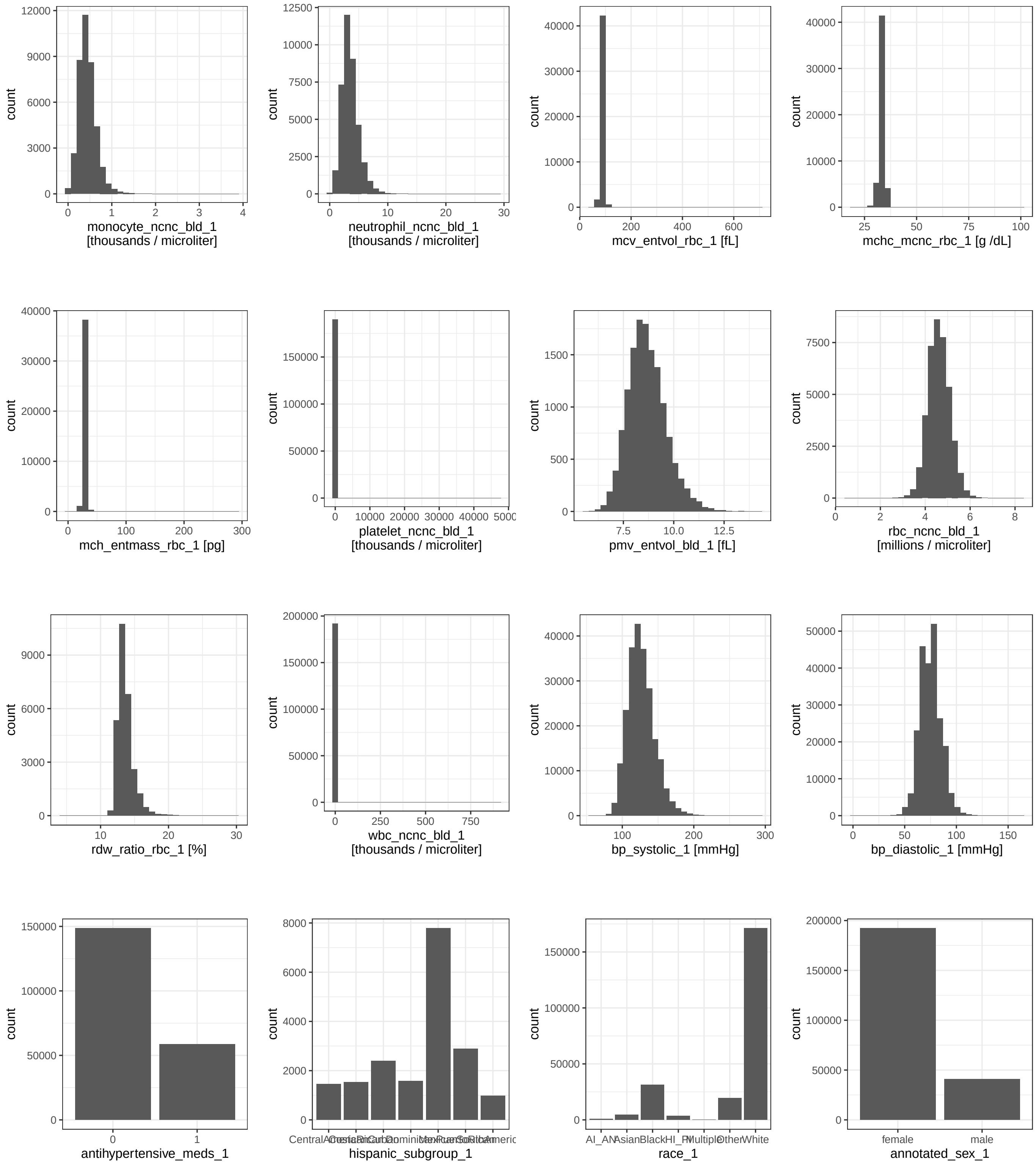

Figure S4: (cont.) Histograms of the harmonized variables presented in this paper.

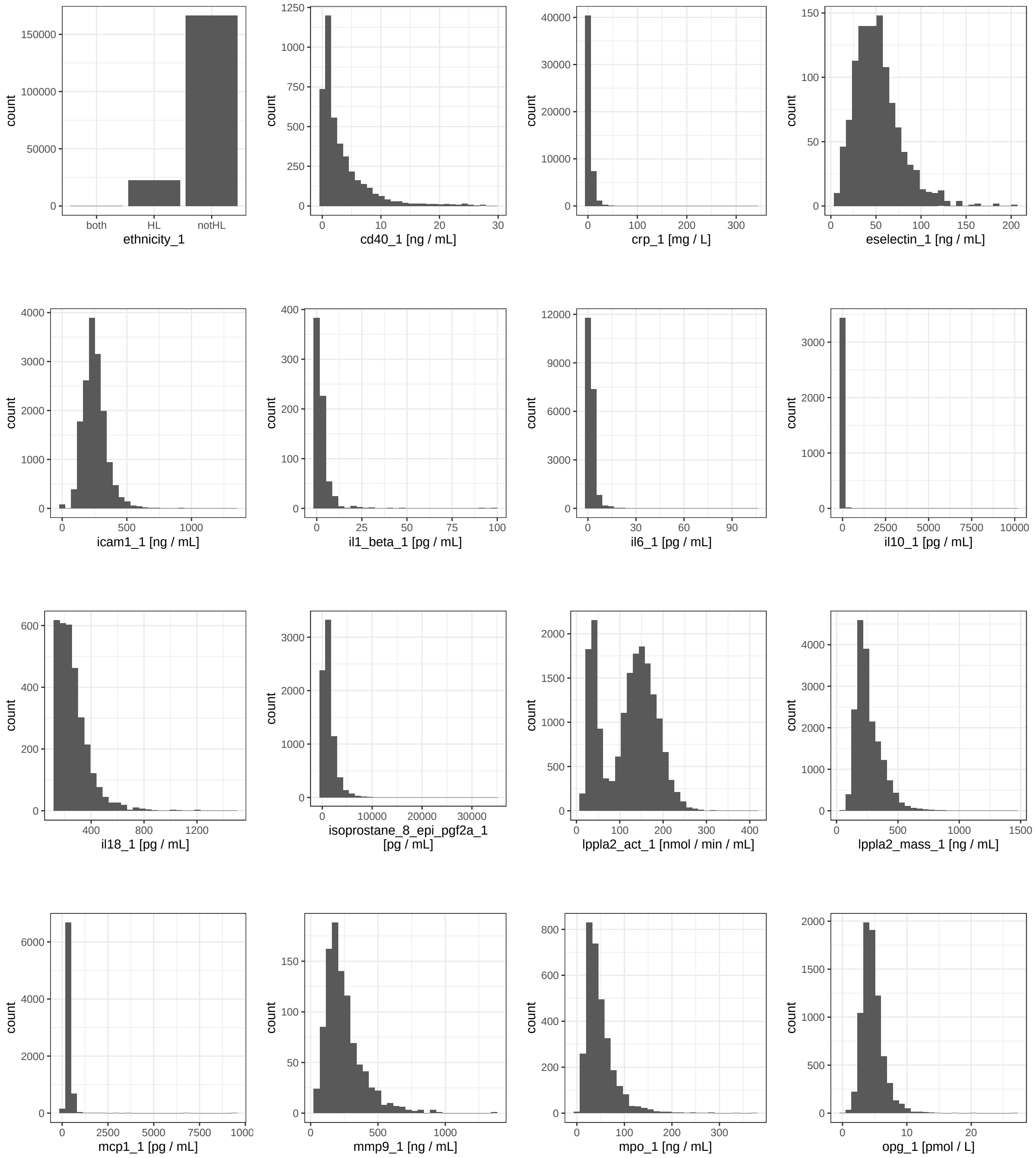

Figure S4: (cont.) Histograms of the harmonized variables presented in this paper.

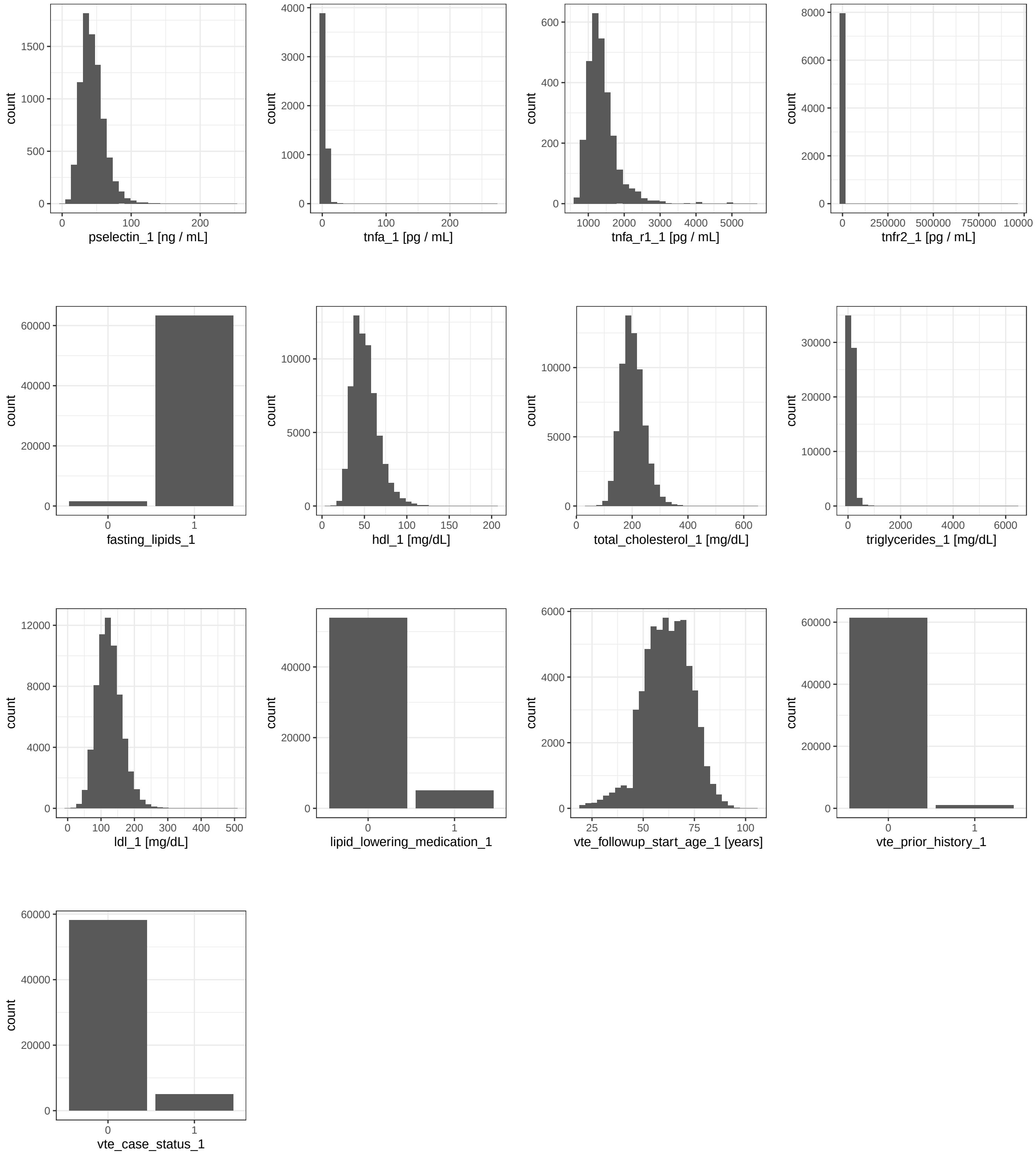

Figure S4: (cont.) Histograms of the harmonized variables presented in this paper.

5. the set of component variables and harmonization function used for each harmonization unit included in the harmonized phenotype variable.

Once an investigator obtains access to the component study variables on dbGaP, the documentation allows them to recreate each harmonized variable exactly. It also enables them to customize the harmonized variable by modifying the harmonization functions and component variables; excluding some harmonization units; using a different definition; or using study variables from a different time point. The repository also includes a reproducible example that shows users how to use the documentation to recreate an example harmonized variable using simulated dbGaP data.

When the DCC updates a harmonized phenotype variable, the documentation files in this repository are updated to reflect the new version.

### S7 Phenotype tagging detailed methods

#### S7.1 Motivation

As detailed above, the process for producing a quality-controlled harmonized variable is very involved. However, there is a need for additional harmonized variables beyond the capacity of the DCC's harmonization team. There are also many reasons that an investigator might want to perform additional harmonization of phenotype concepts for which we have already produced a harmonized variable:

- To use different component variables
- To use a different harmonization algorithm
- To use a different phenotype definition
- To use component variables from a different time point
- To include additional time points
- To include additional non-TOPMed studies

To support additional harmonization efforts by the scientific community, we identified step 2 of the harmonization process ("Identify candidate phenotype variables across contributing studies") as a very time-consuming step that future harmonization efforts would need to repeat. There are many reasons that finding dbGaP variables to harmonize is time-consuming:

- There may be tens of thousands of variables in a single study
- Variable names are often related to the data collection form, rather than to the phenotype content of the variable
- Variable descriptions may not be fully informative and you may need additional information from the data collection forms or the dataset and encoded value documentation to determine the phenotype content of the variable
- Multiple synonyms may be used for the same phenotype
- Phenotype terminology may change over time
- There are variables for multiple measurements from different timepoints and/or clinic visits to select from

In order to reduce the amount of time spent on step 2 for future harmonization efforts, we set out to tag TOPMed dbGaP study variables with controlled vocabulary terms to indicate the phenotype they represent. Just as you might tag your friend's face with their name in a photo on Facebook, we sought to tag dbGaP variables with a label for the relevant phenotype concept. For example, the variable "MF67" (phv00000539) from Framingham Heart Study with the variable description "HEIGHT: FULL INCHES, EXAM 1" can be labeled with the tag "Height". This allows future researchers to easily find all of the TOPMed dbGaP variables tagged "Height", speeding along step 2 for a project to harmonize height according to a different harmonization method. The result is an increase in findability within the TOPMed dbGaP phenotype data, one of the FAIR (Findable, Accessible, Interoperable, and Reusable) guiding principles for scientific data management and stewardship (7).

See separate Excel file.

Table S7: Title, legend, and table provided separately in Excel.

#### S7.2 dbGaP phenotype variables for tagging

We prioritized tagging for dbGaP phenotype variables from the seven large cohort studies included in TOPMed and available to us via dbGaP at the time: ARIC, CARDIA, CHS, FHS, JHS, MESA, and WHI (see Web Table S1 for study abbreviations). These seven studies contained 131,563 dbGaP phenotype variables to consider for tagging. We worked with phenotype data experts from these studies, funded via subcontracts, to complete the tagging.

Ten additional TOPMed studies were also available for tagging. Members of our TOPMed DCC phenotype team completed tagging for 4,409 dbGaP phenotype variables from these smaller studies. In total, 135,972 dbGaP phenotype variables from 17 studies were considered for tagging. Main text Table 3 summarizes the dbGaP phenotype data available for tagging and total number of tagged variables.

#### S7.3 Defining phenotype concepts

With input from NHLBI TOPMed program officers, the TOPMed phenotype harmonization committee, and DCC clinical experts, we developed a list of 65 high priority phenotype concepts with which we wanted to tag variables. We then worked with domain experts from the TOPMed WGs and the TOPMed phenotype harmonization committee to develop clear and concise definitions of each of the phenotype concepts. We also worked with domain experts to develop detailed instructions for which kinds of variables to tag with each phenotype concept. Wherever possible, the instructions we developed included examples of the kinds of variables to include in the tag, and also examples of the kinds of variables that should not be included. Every effort was made to keep the definitions and detailed instruction consistent across phenotype tags.

We attempted to identify an existing phenotype ontology or controlled vocabulary to use, rather than developing our own phenotype concepts and definitions. However, we could not identify a single system that could accommodate all 65 of the phenotype concepts we had determined to capture. Many existing ontologies, such as LOINC (8), SNOMED CT (9), and PhenX (10), were too specific for the task we were trying to accomplish. For example, LOINC has different terms for different lab assays taking the same kind of measurement, whereas we wanted to capture all of the different assays for one kind of measurement in one tag. Other systems, such as MedGen (11), didn't have existing terms for all 65 of the high-priority phenotype concepts we had identified. Others were missing terms for non-disease state measurements; for example, the Human Phenotype Ontology (HPO) (12) has terms for "abnormality of body height", "short stature", and "tall stature", but no term for a quantitative measure of height.

We mapped our 65 detailed phenotype concepts to terms from the UMLS (6) in order to connect our phenotype concepts to existing controlled vocabularies. UMLS is a metathesaurus linking terms across many controlled vocabularies and ontologies, including LOINC, SnoMed, and HPO. Using our mappings to UMLS terms, similar phenotype terms can be linked across multiple vocabularies. Because UMLS contains terms from multiple vocabularies, we were able to find matching terms for all of our 65 phenotype concepts. Web Table S7 provides definitions of the phenotype concepts, as well as their corresponding UMLS terms.

#### S7.4 Tagging user interface

To provide a convenient interface for TOPMed DCC phenotype team members to search and browse the database containing TOPMed dbGaP study phenotype variables, we developed a web application, Phenotype Inventory Explorer (PIE). PIE is written with the Python web framework Django, with templates built on Twitter's Bootstrap HTML, CSS, and JS toolkit (3.3.6). dbGaP study variable metadata are imported from the phenotype harmonization relational database to a separate MariaDB database serving as the PIE backend. Only publicly available metadata (i.e. none of the controlled-access dbGaP phenotype data values) are imported into PIE.

We added tagging functionality to PIE, incorporating several permission, versioning, and data validation features that would not otherwise have been possible. The PIE site administrator created tag objects for each of the 65 phenotype concepts defined in the previous section. We granted permission for tagging dbGaP study variables on a per-study basis according to the study or studies each user is affiliated with. A user with permission may apply a tag to a dbGaP study variable from the variable's detail page, from a tag's detail page, or from a form allowing selection of a tag and entry of multiple study variable accession numbers. Study variables to be tagged may be located via a search page with advanced search filters, or by browsing datasets and variables per study.

When a tag is applied to a study variable, a tagged variable object is created in the backend database, tracking the creator of the tagged variable and a creation timestamp. Data is validated upon entry via PIE, ensuring that the following conditions are met before a tagged variable is created:

- The dbGaP study variable accession is valid
- The tag name is valid
- The study variable to be tagged is from the latest version of the dbGaP study
- The tagged variable is not a duplicate of previously existing tag-study variable pairs

All of the form fields for selecting tags or dbGaP study variables to tag are enabled with string autocompletion to prevent data entry errors and make the tagging process as efficient as possible.

After the creation of tagged variables, PIE allows browsing tagged variables by study and by tag. Tag detail pages display summary counts of the number of tagged variables per study and study detail pages display summary counts of the number of tagged variables per tag. Study variable detail pages display any tags linked to the variable.

#### S7.5 Tagging process

We provided training webinars with demonstrations of tagging on PIE to train the study data experts who participated. Some of the larger studies had more than one phenotype data expert involved, resulting in 11 phenotype data experts from 7 TOPMed cohort studies. We guided the phenotype data experts through completing the tagging on a six month timeline with three intermediate milestone goals.

We set up a mailing list for study data experts to submit questions that might come up during the tagging process. DCC phenotype team members answered technical questions about the tagging functionality on PIE as well as conceptual questions about the interpretation of instructions for specific tags. The questions we received were often asking for additional guidance on whether a tag should be applied to specific dbGaP study variables. DCC phenotype team members answered all of these questions within one or two days, and in some cases consulted with clinical domain experts to provide an answer. The mailing list archive proved a valuable resource for finding related questions and their answers. DCC phenotype team members regularly reviewed the archive of answered questions to ensure consistent application of the tags across studies and across similar kinds of phenotype concepts. We used the feedback from these questions to modify the tagging instructions for clarity, often including additional examples of the kinds of study variables to include or not include in the application of a given tag.

In a few rare cases we revised the tag definition and instructions more substantially in response to the questions we received. For example, we received several questions about the "systolic blood pressure" and "diastolic blood pressure" tags that we initially defined. Based on these questions we determined that our initial phenotype concept definition and instructions didn't account for the wide variety of instruments for blood pressure measurement or the multiple conditions in which blood pressure is routinely measured. In response, we changed these tags to the more specifically named "resting arm diastolic blood pressure" and "resting arm systolic blood pressure" to indicate that we wanted to include only measures of blood pressure taken from the arm at a resting state. Our initial instructions excluded measurements by Doppler/ultrasound and mentioned only sphygmomanometer as a measurement device to include, but our revised instructions stated to include blood pressure "measured by any device, including mercury or other manometer, aneroid gauge, oscillometric device, or Doppler/ultrasound". Note that tag definitions are often broader than those for DCC-harmonized variables. For example, the definition of the DCC-harmonized variables for blood pressure specified that measurements must be collected using a sphygmomanometer.

#### S7.6 Tagging review process

In order to ensure consistency across studies and across tags for similar phenotype concepts, we performed quality review as part of the tagging process. The functionality to accomplish quality review of the tagging data was added to PIE with a straightforward and easy to use interface. The quality review process consisted of up to three rounds of review:

1. Initial review by the DCC phenotype team
2. Opportunity for response from the study phenotype experts
3. Final decision by the DCC phenotype team

In step 1, members of the DCC phenotype team inspect each tagged variable to assess whether it is consistent with the tag (description and instructions) and the study variable (variable description, dataset description, and any available documentation or data collection forms). The review page on PIE displays all of this information on one screen, along with links to more detailed information available on dbGaP, to allow for speedy, accurate, and easy review. After inspecting the tag and study variable information displayed, the DCC phenotype team member either confirms the accuracy of the tagged variable by clicking on a "Confirm" button, or flags the tagged variable for further review by clicking on a "Require study followup" button and providing a brief comment describing why the study variable should not have the tag applied to it. Tagged variables that are confirmed in step 1 require no further review.

In step 2, the study phenotype data experts inspect each of the tagged variables that are flagged for further review in step 1. As in step 1, a review page on PIE displays all relevant information on the tag (description and instructions) and the dbGaP study variable (variable description, dataset description, and links to detailed information on dbGaP). This step 2 review page also includes the comment provided by the DCC in step 1 explaining why the tagged variable is flagged for further review. From here, the study phenotype data expert either agrees to remove the tagged variable or provides a comment explaining why they think it should not be removed. To ask that the tagged variable not be removed, the study phenotype data expert clicks an "Explain why not to remove" button and provides a comment. To agree to removal of the tagged variable, the study phenotype data expert clicks a "Yes, remove tag" button. If a study phenotype data expert agrees to remove a tagged variable during this review step, the tagged variable is not deleted, but archived, preserving all of its related data in the PIE database. These archived tagged variables are not displayed on PIE or included in any counts of tagged variables, and they are excluded from tagging data exports. Tagged variables that are archived in step 2 required no further review.

In step 3, members of the DCC phenotype team inspect each of the tagged variables that are not archived in step 2. A review page on PIE displays all relevant information shown in the previous review steps, along with a timeline showing the actions taken in steps 1 and 2 of the review process. A DCC phenotype team member reviews all of this information and may consult with phenotype domain experts, clinical data experts, other phenotype team members, and the study phenotype data expert before coming to a final decision on whether or not to keep the tagged variable. To keep the tagged variable, the DCC phenotype team member clicks on a "Confirm" button and provides a comment explaining why they decided to keep the tagged variable. To remove the tagged variable, the DCC phenotype team member clicks on a "Remove" button and provides a comment explaining why. Tagged variables that are marked for removal in step 3 are archived as described for step 2. Detail pages for each tagged variable object, even those that have been archived, display the entire history of the review process for that tagged variable.

For tagged variables from the ten studies initially tagged by DCC phenotype team members, the quality review process consisted of a single step. A different DCC phenotype team member than the one who created the tagged variable inspects the tagged variable and its detailed information on a PIE review page and either confirms the tagged variable or flags it for removal. Tagged variables flagged for removal in this step are immediately archived as described above, along with a comment explaining why.

#### S7.7 Tagging review results

15,912 of 17,063 tagged variables passed the review process. The majority of these tagged variables were created by study data experts and reviewed by members of the DCC phenotype team in a 3-step review process. Roughly 13% of the tagged variables (1,194 of 17,063) were initially created by members of the DCC phenotype team and reviewed by another team member in a 1-step review process.

Rates of review decisions were compared across reviewers, studies, and phenotype tags and no notable differences were observed, with one exception. The "AHI" phenotype tag had a high proportion of tagged variables fail review (~48%) (Web Figure S5). Investigation determined that this was attributed to a very large number of study variables with nearly identical variable names and variable descriptions representing multiple clinic visits in a single study. These highly similar variables were all tagged by the study data expert as "AHI", but determined by the DCC phenotype team not to agree with the phenotype concept definition and instructions. Therefore this high proportion of tagged variables failing review could be explained by a single differing interpretation of tagging instructions, repeated over many similar study variables. The "Carotid IMT" phenotype tag also had a somewhat high (~19%) proportion of tagged variables fail review for a similar reason.

#### S7.8 Tagging results

We tagged dbGaP study variables with UMLS terms representing 65 phenotype concepts in 16 domains. A total of 16,671 dbGaP phenotype variables from 17 studies are now tagged with relevant UMLS phenotype terms. Because some study variables may be tagged with multiple phenotype terms, there are 17,063 unique pairings of dbGaP study variable and UMLS phenotype term. Main text Table 3 shows the proportion of study variables tagged in each study, along with the total number of study variables available per study. The proportion of dbGaP study variables tagged is generally proportional to the number of dbGaP study variables that were available for tagging in each study. Studies with a larger number of dbGaP study variables (e.g., FHS) had a much smaller proportion of study variables tagged. For these studies, the 65 prioritized phenotype concepts included in the tagging process represent a fraction of the phenotype concepts for which data have been collected.

The number of study variables tagged for each phenotype concept presents an overview of the variety of phenotypes collected for TOPMed studies (Web Table S8). For example, "Medication/supplement use", "Cigarette smoking", and "Carotid IMT" have the greatest number of study variables tagged, indicating an abundance of data collected for these phenotypes. This could also be a good way to identify new genetic analysis opportunities in TOPMed - phenotype concepts with large numbers of tagged variables, but few published analyses, could be prioritized.

We can also examine the number of studies with at least one study variable tagged for each phenotype concept. Web Figure S6 shows the cumulative frequency of this number. For example, 31 phenotype concepts have at least 8 studies represented in tagged variables for that concept. Web Figure S6 shows a steady increase in cumulative frequency. About half of the studies are represented in tagged variables for about half of the phenotype concepts.

The results of the tagging project have already served as an invaluable resource for identifying candidate component variables for new DCC harmonization projects. Rather than compiling the results from multiple searches of key terms in study variable names, variable descriptions, and encoded values, DCC harmonization team members can instead pull up a list of all of the study variables tagged with a particular phenotype concept. This set of tagged variables was produced by data experts and carefully quality reviewed with input from domain experts, and therefore can be used as a gold standard to use for training, testing, and validation

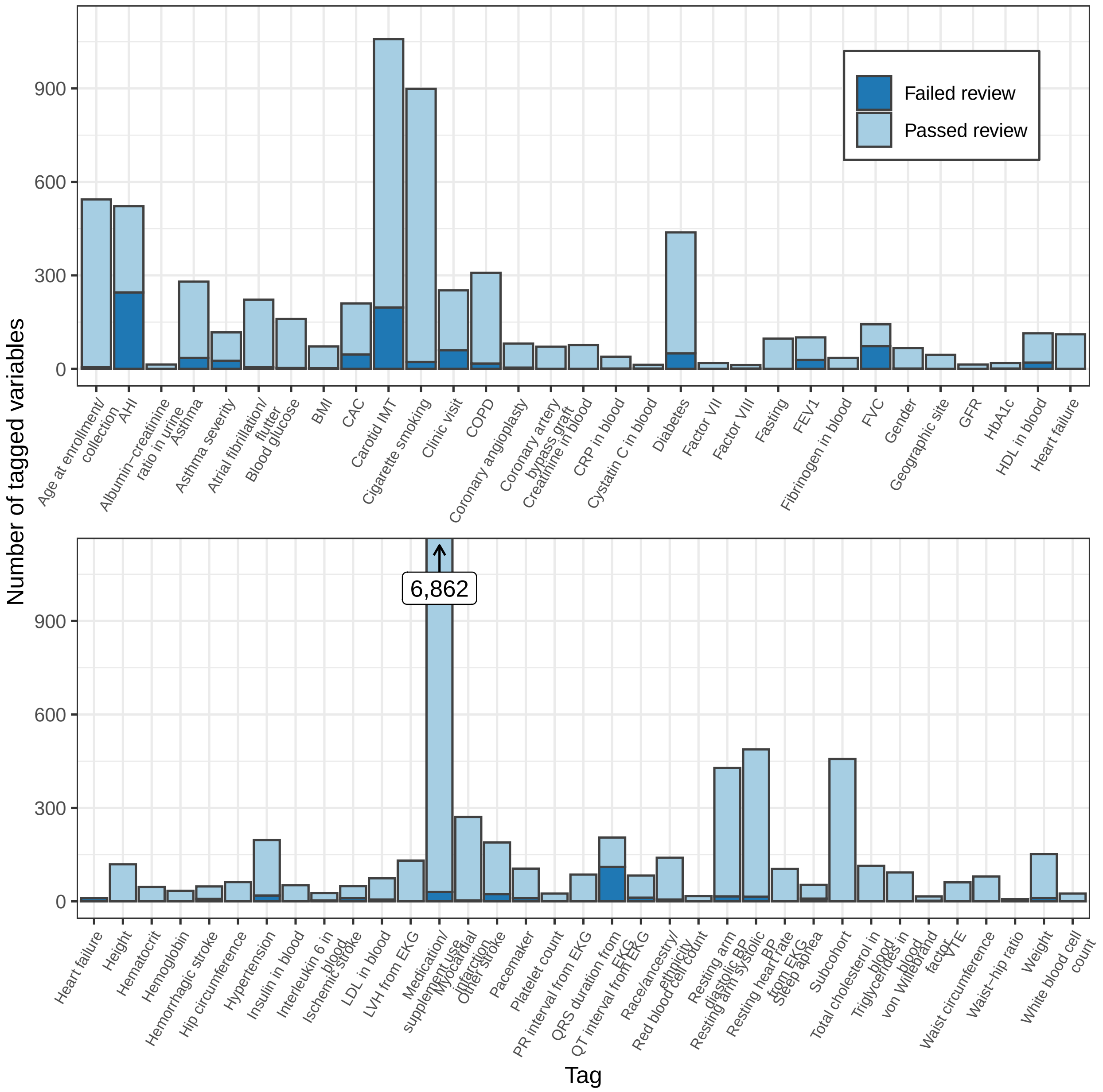

Figure S5: Tagging and quality review results by phenotype concept. Arrow and label for “Medication/supplement use” bar indicates the very high number of study variables tagged for this phenotype concept.

of Natural Language Processing methods for automated tagging. We are already working in coordination with developers from the NHLBI BioData Catalyst platform to develop automated tagging solutions.

Versioning tools in PIE enable the automatic tagging of new versions of study variables that were previously tagged, so that as new versions of TOPMed study accessions are released the tagged variables can remain connected to the most recent version of a dbGaP study variable. As new studies are added to TOPMed, PIE can be used to tag study variables for the 65 prioritized phenotype concepts. New phenotype concept tags can also be added to PIE to expand the project scope.

Table S8: Count of study variables tagged with each phenotype concept by study.

| Phenotype tag | MESA | Amish | JHS | ARIC | CHS | Samoan | CFS | WHI | FHS | HCHS/SOL | CARDIA | GENOA | GOLDN | COPDGene | CRA | HVH | Mayo_VTE | Total | N studies |
|---|---|---|---|---|---|---|---|---|---|---|---|---|---|---|---|---|---|---|---|
| LDL in blood | 12 | 1 | 3 | 10 | 7 | 2 | 2 | 5 | 19 | 1 | 6 | 0 | 0 | 0 | 0 | 0 | 0 | 68 | 11 |
| HDL in blood | 10 | 1 | 3 | 9 | 4 | 2 | 3 | 5 | 42 | 1 | 6 | 7 | 1 | 0 | 0 | 0 | 0 | 94 | 13 |
| Triglycerides in blood | 9 | 1 | 3 | 9 | 4 | 2 | 2 | 5 | 43 | 1 | 6 | 7 | 1 | 0 | 0 | 0 | 0 | 93 | 13 |
| Total cholesterol in blood | 10 | 1 | 3 | 9 | 14 | 2 | 2 | 5 | 54 | 1 | 6 | 7 | 0 | 0 | 0 | 0 | 0 | 114 | 12 |
| Resting arm systolic BP | 43 | 1 | 14 | 34 | 11 | 2 | 20 | 6 | 207 | 3 | 85 | 45 | 1 | 1 | 0 | 0 | 0 | 473 | 14 |
| Resting arm diastolic BP | 35 | 1 | 11 | 31 | 11 | 2 | 16 | 6 | 172 | 1 | 83 | 41 | 1 | 1 | 0 | 0 | 0 | 412 | 14 |
| Height | 13 | 1 | 1 | 6 | 5 | 1 | 1 | 3 | 62 | 1 | 13 | 8 | 0 | 1 | 1 | 1 | 1 | 119 | 16 |
| Weight | 16 | 1 | 1 | 10 | 12 | 1 | 2 | 4 | 65 | 1 | 16 | 8 | 0 | 1 | 1 | 1 | 1 | 141 | 16 |
| BMI | 14 | 1 | 1 | 5 | 5 | 1 | 2 | 3 | 28 | 1 | 6 | 0 | 1 | 1 | 1 | 0 | 0 | 70 | 14 |
| Waist circumference | 8 | 1 | 1 | 5 | 6 | 1 | 2 | 2 | 32 | 1 | 14 | 7 | 0 | 0 | 0 | 0 | 0 | 80 | 12 |
| Hip circumference | 8 | 0 | 0 | 5 | 4 | 1 | 2 | 1 | 20 | 1 | 13 | 7 | 0 | 0 | 0 | 0 | 0 | 62 | 10 |
| Waist-hip ratio | 0 | 1 | 0 | 4 | 0 | 0 | 0 | 1 | 0 | 1 | 0 | 0 | 0 | 0 | 0 | 0 | 0 | 7 | 4 |
| Ischemic stroke | 6 | 0 | 0 | 10 | 1 | 0 | 0 | 4 | 11 | 0 | 5 | 0 | 0 | 0 | 0 | 2 | 0 | 39 | 7 |
| Hemorrhagic stroke | 3 | 0 | 0 | 14 | 1 | 0 | 0 | 4 | 11 | 0 | 5 | 0 | 0 | 0 | 0 | 2 | 0 | 40 | 7 |
| Other stroke | 7 | 1 | 14 | 65 | 8 | 1 | 4 | 0 | 25 | 0 | 19 | 18 | 0 | 1 | 0 | 2 | 1 | 166 | 13 |
| Age at enrollment/collection | 31 | 3 | 31 | 8 | 14 | 1 | 2 | 99 | 314 | 1 | 21 | 9 | 1 | 1 | 1 | 1 | 1 | 539 | 17 |
| Gender | 12 | 1 | 10 | 5 | 10 | 1 | 0 | 0 | 5 | 2 | 13 | 2 | 1 | 1 | 1 | 1 | 1 | 66 | 15 |
| Race/ancestry/ethnicity | 29 | 0 | 0 | 4 | 8 | 0 | 8 | 17 | 42 | 2 | 10 | 2 | 0 | 2 | 1 | 2 | 7 | 134 | 13 |
| CAC | 46 | 1 | 1 | 0 | 1 | 0 | 0 | 0 | 26 | 0 | 86 | 3 | 0 | 0 | 0 | 0 | 0 | 164 | 7 |
| Carotid IMT | 215 | 1 | 46 | 439 | 112 | 0 | 0 | 0 | 28 | 0 | 20 | 0 | 0 | 0 | 0 | 0 | 0 | 861 | 7 |
| Myocardial infarction | 12 | 1 | 25 | 52 | 9 | 1 | 5 | 0 | 124 | 2 | 19 | 15 | 0 | 1 | 0 | 1 | 1 | 268 | 14 |
| Coronary angioplasty | 8 | 0 | 11 | 10 | 5 | 0 | 2 | 8 | 15 | 0 | 2 | 15 | 0 | 1 | 0 | 0 | 0 | 77 | 10 |
| Coronary artery bypass graft | 4 | 0 | 11 | 3 | 5 | 0 | 1 | 9 | 20 | 0 | 2 | 15 | 0 | 1 | 0 | 0 | 0 | 71 | 10 |
| Heart failure | 4 | 0 | 11 | 4 | 7 | 0 | 4 | 25 | 43 | 1 | 11 | 0 | 0 | 1 | 0 | 0 | 0 | 111 | 10 |
| Hypertension | 24 | 0 | 14 | 24 | 15 | 5 | 7 | 5 | 25 | 0 | 33 | 24 | 0 | 1 | 0 | 1 | 0 | 178 | 12 |
| Blood glucose | 14 | 1 | 3 | 9 | 10 | 2 | 4 | 5 | 92 | 2 | 9 | 5 | 1 | 0 | 0 | 0 | 0 | 157 | 13 |
| Insulin in blood | 5 | 1 | 2 | 4 | 4 | 1 | 4 | 6 | 9 | 2 | 8 | 5 | 0 | 0 | 0 | 0 | 0 | 51 | 12 |
| HbA1c | 3 | 0 | 2 | 0 | 0 | 0 | 0 | 4 | 8 | 1 | 0 | 0 | 0 | 0 | 0 | 0 | 0 | 18 | 5 |
| Diabetes | 48 | 1 | 18 | 23 | 36 | 4 | 10 | 4 | 188 | 6 | 33 | 15 | 0 | 1 | 0 | 1 | 0 | 388 | 14 |
| Atrial fibrillation/flutter | 27 | 0 | 0 | 53 | 36 | 0 | 0 | 14 | 85 | 1 | 0 | 0 | 0 | 0 | 0 | 1 | 0 | 217 | 7 |
| QRS duration from EKG | 5 | 1 | 14 | 12 | 12 | 0 | 1 | 1 | 47 | 1 | 0 | 0 | 0 | 0 | 0 | 0 | 0 | 94 | 9 |
| QT interval from EKG | 4 | 1 | 1 | 5 | 12 | 0 | 2 | 2 | 42 | 2 | 0 | 0 | 0 | 0 | 0 | 0 | 0 | 71 | 9 |
| PR interval from EKG | 9 | 1 | 12 | 4 | 11 | 0 | 1 | 1 | 45 | 1 | 0 | 0 | 0 | 0 | 0 | 0 | 0 | 85 | 9 |
| Resting heart rate from EKG | 4 | 1 | 1 | 19 | 11 | 0 | 0 | 1 | 67 | 0 | 0 | 0 | 0 | 0 | 0 | 0 | 0 | 104 | 7 |
| LVH from EKG | 8 | 0 | 1 | 37 | 17 | 0 | 0 | 3 | 64 | 0 | 0 | 0 | 0 | 0 | 0 | 0 | 0 | 130 | 6 |
| Pacemaker | 15 | 0 | 2 | 23 | 3 | 0 | 0 | 2 | 50 | 0 | 0 | 0 | 0 | 0 | 0 | 0 | 0 | 95 | 6 |
| Hematocrit | 2 | 1 | 1 | 4 | 4 | 0 | 0 | 6 | 26 | 1 | 1 | 0 | 0 | 0 | 0 | 0 | 0 | 46 | 9 |
| Hemoglobin | 3 | 1 | 1 | 4 | 4 | 0 | 0 | 6 | 13 | 1 | 1 | 0 | 0 | 0 | 0 | 0 | 0 | 34 | 9 |
| Platelet count | 2 | 1 | 1 | 4 | 4 | 0 | 0 | 5 | 6 | 1 | 1 | 0 | 0 | 0 | 0 | 0 | 0 | 25 | 9 |
| Red blood cell count | 2 | 1 | 1 | 2 | 0 | 0 | 0 | 4 | 5 | 1 | 1 | 0 | 0 | 0 | 0 | 0 | 0 | 17 | 8 |
| White blood cell count | 2 | 1 | 1 | 4 | 4 | 0 | 0 | 5 | 6 | 1 | 1 | 0 | 0 | 0 | 0 | 0 | 0 | 25 | 9 |
| Fibrinogen in blood | 4 | 1 | 0 | 2 | 4 | 0 | 0 | 5 | 11 | 0 | 5 | 3 | 0 | 0 | 0 | 0 | 0 | 35 | 8 |
| Factor VII | 0 | 0 | 0 | 2 | 4 | 0 | 0 | 6 | 3 | 0 | 4 | 0 | 0 | 0 | 0 | 0 | 0 | 19 | 5 |

Table S8: *(continued)*

| Phenotype tag | MESA | Amish | JHS | ARIC | CHS | Samoan | CFS | WHI | FHS | HCHS/SOL | CARDIA | GENOA | GOLDN | COPDGene | CRA | HVH | Mayo_VTE | Total | N studies |
|---|---|---|---|---|---|---|---|---|---|---|---|---|---|---|---|---|---|---|---|
| Factor VIII | 3 | 0 | 0 | 1 | 1 | 0 | 0 | 4 | 0 | 0 | 2 | 0 | 0 | 0 | 0 | 0 | 0 | 11 | 5 |
| von Willebrand factor | 1 | 0 | 0 | 1 | 0 | 0 | 0 | 4 | 3 | 0 | 5 | 0 | 0 | 0 | 0 | 0 | 0 | 14 | 5 |
| VTE | 0 | 0 | 0 | 4 | 4 | 0 | 0 | 4 | 36 | 0 | 10 | 0 | 0 | 0 | 0 | 2 | 1 | 61 | 7 |
| CRP in blood | 7 | 1 | 1 | 0 | 4 | 0 | 8 | 4 | 7 | 0 | 3 | 3 | 0 | 0 | 0 | 0 | 0 | 38 | 9 |
| Interleukin 6 in blood | 5 | 1 | 0 | 0 | 1 | 0 | 0 | 4 | 12 | 0 | 1 | 0 | 0 | 0 | 0 | 0 | 0 | 24 | 6 |
| Creatinine in blood | 7 | 0 | 2 | 9 | 8 | 0 | 3 | 5 | 31 | 1 | 3 | 7 | 0 | 0 | 0 | 0 | 0 | 76 | 10 |
| Cystatin C in blood | 2 | 0 | 0 | 0 | 0 | 0 | 2 | 4 | 2 | 0 | 0 | 2 | 0 | 0 | 0 | 0 | 0 | 12 | 5 |
| Albumin-creatinine ratio in urine | 6 | 0 | 2 | 0 | 1 | 0 | 2 | 0 | 0 | 1 | 2 | 0 | 0 | 0 | 0 | 0 | 0 | 14 | 6 |
| GFR | 13 | 0 | 1 | 0 | 0 | 0 | 0 | 0 | 0 | 0 | 0 | 0 | 0 | 0 | 0 | 0 | 0 | 14 | 2 |
| FVC | 3 | 1 | 4 | 2 | 7 | 0 | 0 | 0 | 19 | 2 | 29 | 0 | 0 | 3 | 0 | 0 | 0 | 70 | 9 |
| FEV1 | 3 | 1 | 4 | 2 | 7 | 0 | 0 | 0 | 21 | 1 | 30 | 0 | 0 | 3 | 0 | 0 | 0 | 72 | 9 |
| Asthma | 32 | 0 | 17 | 14 | 36 | 0 | 6 | 8 | 93 | 5 | 26 | 0 | 0 | 7 | 1 | 0 | 0 | 245 | 11 |
| Asthma severity | 19 | 0 | 0 | 0 | 25 | 0 | 2 | 0 | 31 | 3 | 10 | 0 | 0 | 1 | 0 | 0 | 0 | 91 | 7 |
| COPD | 32 | 0 | 5 | 17 | 32 | 0 | 6 | 12 | 149 | 1 | 18 | 6 | 0 | 13 | 0 | 0 | 0 | 291 | 11 |
| Sleep apnea | 3 | 0 | 0 | 3 | 2 | 0 | 6 | 1 | 25 | 0 | 0 | 0 | 0 | 4 | 0 | 0 | 0 | 44 | 7 |
| AHI | 0 | 0 | 0 | 7 | 8 | 0 | 7 | 0 | 254 | 1 | 0 | 0 | 0 | 0 | 0 | 0 | 0 | 277 | 5 |
| Cigarette smoking | 67 | 1 | 24 | 53 | 55 | 10 | 8 | 34 | 364 | 9 | 192 | 35 | 0 | 17 | 6 | 1 | 1 | 877 | 16 |
| Subcohort | 3 | 0 | 1 | 0 | 15 | 0 | 0 | 13 | 425 | 0 | 0 | 0 | 0 | 0 | 0 | 0 | 0 | 457 | 5 |
| Clinic visit | 0 | 0 | 39 | 5 | 13 | 0 | 2 | 78 | 37 | 2 | 4 | 9 | 0 | 2 | 0 | 1 | 0 | 192 | 11 |
| Fasting | 8 | 0 | 4 | 10 | 18 | 1 | 0 | 3 | 34 | 1 | 13 | 5 | 0 | 0 | 0 | 0 | 0 | 97 | 10 |
| Geographic site | 15 | 0 | 1 | 4 | 9 | 1 | 0 | 1 | 8 | 1 | 2 | 0 | 1 | 1 | 0 | 0 | 1 | 45 | 12 |
| Medication/supplement use | 752 | 3 | 359 | 562 | 1319 | 3 | 198 | 535 | 2403 | 61 | 498 | 106 | 0 | 32 | 0 | 0 | 1 | 6832 | 14 |
| Total | 1717 | 40 | 740 | 1680 | 2020 | 48 | 359 | 1011 | 6154 | 132 | 1412 | 441 | 9 | 99 | 13 | 20 | 17 | 15912 | NA |

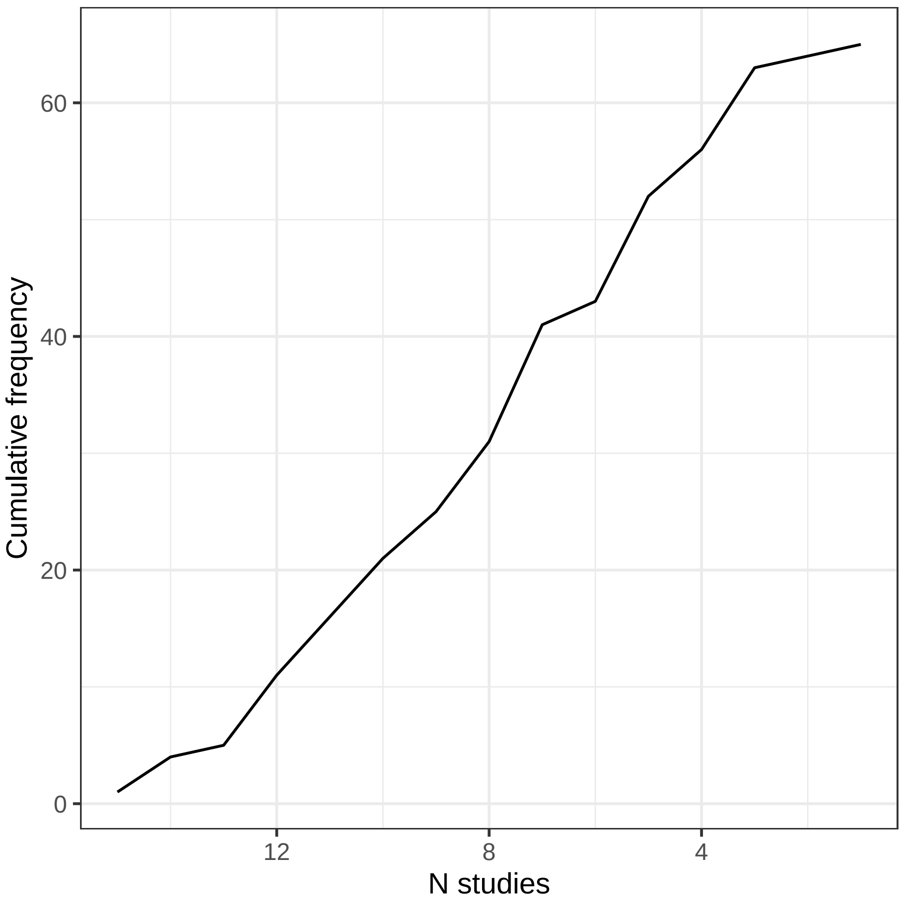

Figure S6: The cumulative frequency of phenotype concepts with at least one study variable tagged for that phenotype concept.

#### S7.9 Availability on dbGaP

We worked with dbGaP scientists to make the tagging information available in dbGaP searches and visible on dbGaP study variable pages. dbGaP users can search for study variables by UMLS term (using the UMLS Concept Unique Identifier, CUI) in either the Entrez search or faceted search. Consult Web Table S7 for UMLS CUIs to use as search terms.

Instructions and a video demo of searching for the tagged variables on dbGaP are available at https://www.nhlbiwgs.org/dcc-pheno.

2. MariaDB 10.2.11 Release Notes. *MariaDB KnowledgeBase*.

3. Friedewald WT, Levy RI, Fredrickson DS. Estimation of the concentration of low-density lipoprotein

cholesterol in plasma, without use of the preparative ultracentrifuge. *Clinical Chemistry.* 1972;18(6):499–502.

4. Dawber TR, Meadors GF, Moore FE. Epidemiological approaches to heart disease: The Framingham Study. *American Journal of Public Health and the Nation's Health.* 1951;41(3):279–281.

5. Bis JC, Kavousi M, Franceschini N, et al. Meta-analysis of genome-wide association studies from the CHARGE consortium identifies common variants associated with carotid intima media thickness and plaque. *Nature Genetics.* 2011;43(10):940–947.

6. Bodenreider O. The Unified Medical Language System (UMLS): Integrating biomedical terminology. *Nucleic Acids Research.* 2004;32(Database issue):D267–D270.
